# Quantitative essentiality in a reduced genome: a functional, regulatory and structural fitness map

**DOI:** 10.1101/2025.02.06.636790

**Authors:** Samuel Miravet-Verde, Raul Burgos, Eva Garcia-Ramallo, Marc Weber, Luis Serrano

**Author notes:** These authors contributed equally to this work (Samuel Miravet-Verde, Raul Burgos).

## Abstract

Essentiality studies have traditionally focused on coding regions, often overlooking other small genetic regulatory elements. To address this, we obtained a high-resolution essentiality map at near-single-nucleotide precision of the genome-reduced bacterium *Mycoplasma pneumoniae*, combining transposon libraries containing promoter or terminator sequences. By integrating temporal transposon-sequencing data, we developed a novel method of essentiality assessment based on k-means unsupervised clustering, which provides dynamic and quantitative information on the fitness contribution of different genomic regions. We compared the insertion tolerance and persistence of the two engineered libraries, assessing the local impact of transcription and termination on cell fitness. Essentiality assessment at the local base-level revealed essential protein domains and small genomic regions that are either essential or inaccessible to transposon insertion. We also identified structural regions within essential genes that tolerate transposon disruptions, resulting in functionally split proteins. Overall, this study presents a nuanced view of gene essentiality, shifting from static and binary models to a more accurate perspective. Additionally, it provides valuable insights for genome engineering and enhances our understanding of the biology of genome-reduced cells.

## INTRODUCTION

Genes are the basic units that define the genome of an organism. Nonetheless, other genomic elements are also expected to contribute to cell function. These may include structural regions governing chromosome replication, or transcriptional and translational regulatory elements that allow coordination of gene activity. Notably, a comprehensive analysis of a genome-reduced organism like *Mycoplasma pneumoniae* has shown the importance of alternative regulatory mechanisms of transcriptional activity that do not depend on transcription factors [1], such as for example RNA degradation, the importance of the first nucleotide of the transcript, supercoiling, or the influence of chromosomal structure [2]. Hence, studies that aim to fully understand how genetic information impacts the fitness of an organism should consider other genomic features aside from genes.

Transposon mutagenesis is a powerful genetic tool to produce random mutations in genomes [3]. The appearance of ultra-deep sequencing technologies has allowed the screen of large pools of transposon mutants simultaneously in a given growth condition [4–8], opening the possibility to perform diverse genome-wide analyses of gene essentiality in a high-throughput manner [9]. In a typical transposon-insertion sequencing (Tn-Seq) experiment, transformed cells with insertions in genes required for growth are lost after a short period of growth selection. This results in a transposon integration map where essential regions remain free of transposon insertions, allowing the identification of essential (E) and non-essential genes (NE). However, NE genes can be classified in subgroup categories depending on how the disruption of these genes causes a competitive defect under a defined set of conditions. These genes, typically known as fitness (F), may be classified as NE or E depending on the growth conditions and rounds of selection. In addition, essential genes may tolerate insertions in specific locations such as N- and C-terminal regions that generally do not form part of the functional unit.

Traditionally, essentiality studies have focused at the gene level with few studies looking at the essentiality of other genomic features, including small genetic entities such as regulatory signals (including promoters and terminators) and non-transcribed regions of the chromosome [10–13]. This is due to the high transposon insertion density required to provide sufficient statistical evidence on the essentiality of small DNA segments (*e.g.*, a promoter region). Ideally, this resolution should approach one insertion per base, with careful consideration given to the design of the transposon used in the analysis. For example, the mariner transposable elements depend on TA dinucleotide targets, limiting the resolution that can be obtained, especially in high GC-content genomes [13]. Additionally, the presence of promoter and terminator sequences within the transposon can bias the preference of insertion depending on the transcriptional context. This can be particularly important when assessing the essentiality of large operons constituted by NE genes followed by E ones, or when gene overlaps happen. In this respect, approaches have been developed placing outward facing promoters in the transposon vector to induce the expression of E genes and minimize polar effects when an operon is disrupted [14].

In this study, we leveraged the small genome of *M. pneumoniae* (816 kb), its high transformation efficiency, and the extensive available -omics datasets [2,15–23], to define a comprehensive and high-resolution fitness map of its genome. We achieved this by designing and transforming the bacterium with two different Tn4001-based transposons. One presents outward-facing promoters at both ends, aimed at minimizing polar effects and exploring the influence of transcriptional changes in fitness. The other transposon features rho-independent terminators at each end, allowing us to investigate the impact of termination or reduced transcription in different genomic loci. By combining both datasets, we found 453,897 unique insertions covering ∼55% of the entire genome, reaching a transposon coverage close to absolute saturation for NE genes (92.4% of coverage, ∼1 bp resolution). This resolution, along with the comparison of the two transposon libraries, enabled us to evaluate the significance of genomic regions beyond genes, such as regulatory signals and structural elements. It also revealed the influence of genome accessibility on transposon sequencing data. Additionally, we assigned essentiality at the protein domain level and identified structural regions within E genes that tolerate disruptions, leading to functional split proteins. Furthermore, we assessed the fitness impact of gene disruptions and transcriptional perturbations by tracking insertion decay after serial passaging selection. This approach allowed us to develop a novel methodology for assessing essentiality, using a k-means unsupervised clustering method that treats essentiality as a dynamic phenomenon.

Collectively, this study provides a resource for understanding the biology of a genome-reduced cell, offering insights on the minimal functions required to sustain life and their regulation, while also providing guidance for the design of proteins or entire biological systems for biotechnological applications.

## RESULTS

### Design and generation of Tn-Seq libraries of *M. pneumoniae* at 1 bp resolution

We designed two transposon vectors based on the mini-transposon pMTnCat [24], which contains a mobile *cat* resistance marker enclosed by inverted repeats (IR), and a transposase derived from Tn4001 [25]. One vector (pMTnCat_BDPr) was engineered to contain two outward facing promoters (P438) adjacent to the two IR (Fig. 1A). This configuration was implemented to minimize transcriptional polar effects on neighboring genes regardless of transposon orientation. The P438 promoter comprises 22 bases and includes two Pribnow boxes promoting constitutive and strong transcription of leaderless mRNAs [26]. Considering most mRNAs in *M. pneumoniae* lack Shine-Dalgarno sequences, translation initiation at the first methionine of the transcript seems the preferent mechanism for protein production [27,28]. Hence, this transposon design provides a system to evaluate the essentiality of potential ORF products as long as the P438 promoter inserts close to a translation initiation codon. A complementary vector (pMTnCat_BDter) containing outward facing intrinsic rho-independent terminator sequences overlapping part of the two IR sequences was also constructed (Fig. 1A). In contrast to pMTnCat_BDPr, this genetic configuration aims to terminate transcription of any transcript originating from, or passing through the transposon sequence once inserted in the genome. For this purpose, we used an endogenous rho-independent terminator (ter625) that silences the expression of the MPN626 sigma factor (Fig. S1A). Termination activity of the ter625 sequence was assessed in two different genetic contexts using gene reporters expressed as polycistronic or fusion transcripts (Fig. S1B). In both cases, we detected a reduction of expression of the reporter in the presence of ter625 as compared to a mutated version affecting the terminator hairpin structure. These results confirmed the functionality of the ter625 sequence.

**Figure 1.**
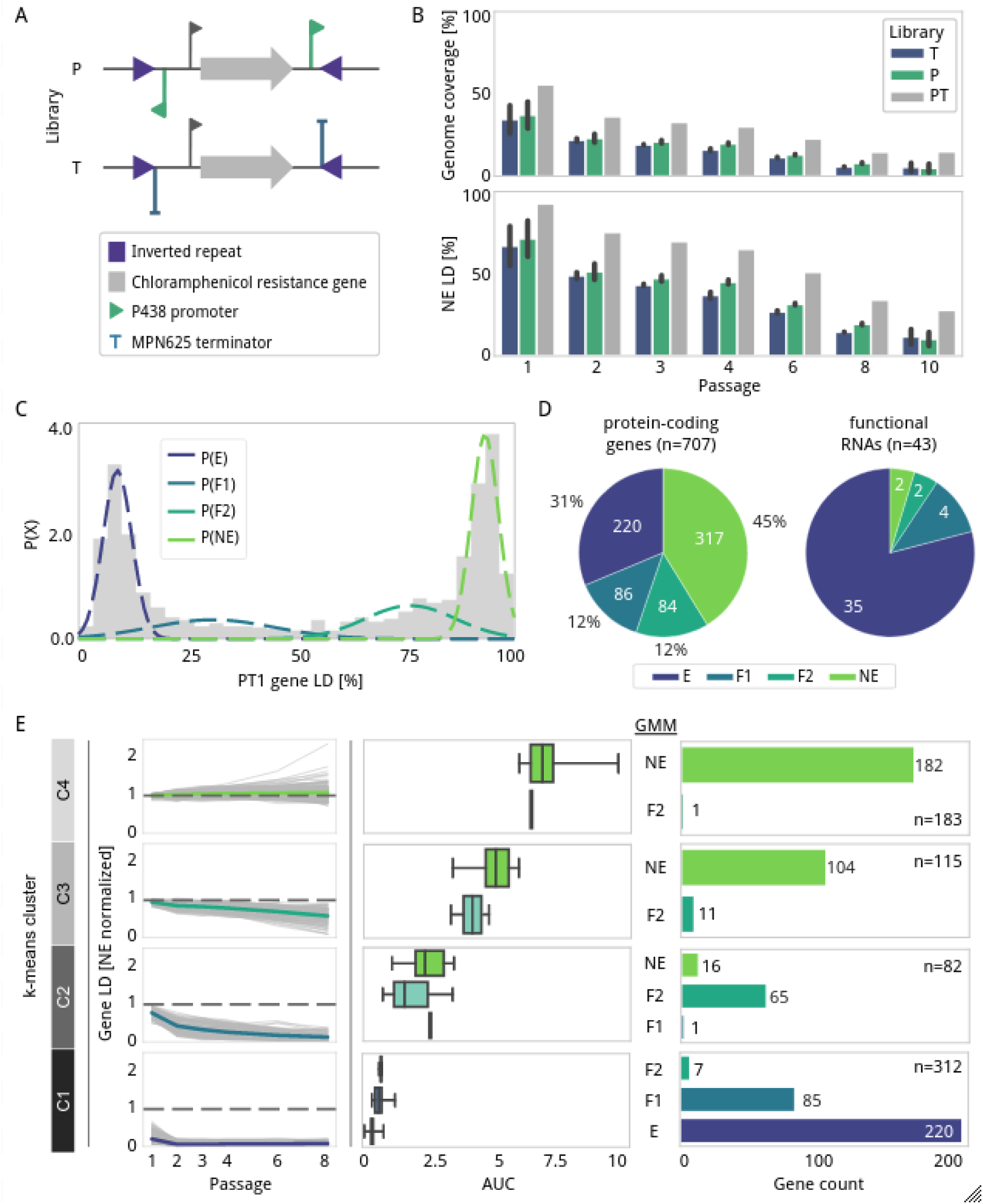
Transposon library design and gene essentiality analysis. **A)** Schematic of the transposon mobile region in the two libraries designed here. P (top) containing outward facing P438 promoters (green flags); and T (bottom) containing ter625 terminators (blue flags). Both constructs carry the chloramphenicol resistance as a marker (gray arrow) flanked by inverted repeat (IR) sequences (purple triangle). Notice that in the case of T libraries the terminator partially overlaps with the IR sequence ‘*TACGGACTTTATC’*. **B)** Bar plots comparing the transposon coverage (Y-axis) obtained along different serial passages (X-axis) for P (green), T (blue) and the combined P and T PT libraries (gray). The top plot shows the coverage of the full genome, whereas the bottom plot considers only known NE genes. Except for PT, which are the result of merging independent samples, the height of the bar represents the average obtained between replicates of the same condition. Error bars represent the standard deviation (n=2). **C)** Gaussian Mixture Model (GMM) of the distribution of linear density (LD) associated with annotated genes (X-axis) against the frequency (Y-axis) in PT1 samples. Dashed overlying lines represent the probability distribution for each essentiality category: E (dark blue), F1 (blue), F2 (turquoise), and NE (green). **D)** Pie charts showing the essentiality category of annotated genes, including protein-coding genes (left) and functional RNAs (right). **E)** Comparative analysis between k-means clusters, area under the curve (AUC) values and GMM categories for protein-coding genes. The first left column shows the decay trajectories (gray lines) in LD (Y-axis) measured from samples PT along passages 1 to 8 (X-axis). For relative comparison purposes, LD decays were normalized to a gold set of known NE genes. The mean trajectory is plotted with a specific color in comparison to a gray dashed line centered at 1 corresponding to the average LD for a gold set of NE genes. The center column represents the AUC values (X-axis) assigned to the genes in each cluster and essentiality category (Y-axis) as defined in Fig, 1C. The right column counts the number of genes in each cluster and the essentially category estimated by GMM. Note that for the clustering analysis we excluded 15 genes with a high percentage of sequence repetition, as these interfere with the clustering prediction (see Methods).

Using the two mini-transposons vectors described above, we constructed two independent libraries of transposon mutants in *M. pneumoniae* M129 strain (‘P’ for pMTnCat_BDPr, and ‘T’ for pMTnCat_BDter). To eliminate death cells that could interfere with the analysis and to define more precisely the contribution of NE genes in cell fitness, we completed up to ten consecutive passages equivalent to approximately ten cell divisions each (samples P1 or T1, to P10 or T10). Two biological replicates per sample were processed for next-generation sequencing and insertion sites were then identified using FASTQINS [29] (Supplementary Data S1). To ensure mapping accuracy, we used as a reference the genome sequence obtained from our laboratory strain, which was re-annotated based on available transcriptomic and proteomic data (see Methods; Supplementary Data S2). In terms of linear density (LD: number of insertions normalized by length of the region considered) and RPKM (reads mapping an insertion per kilobase per million reads), we observed a good correlation between biological replicates for P and T libraries across passages (R^2^ ≥ 0.87 or R^2^ ≥ 0.91 in all conditions, respectively; Fig. S2), suggesting the high transposon coverage in the libraries circumvents possible sampling batch effects.

At genome level, the P library showed an average transposon insertion coverage in P1 of 37.1±8.1% (Fig. 1B), or 46.3% when aggregating insertion events from the two independent P1 datasets (Supplementary Data S3). For the T1 library, the coverage was slightly lower, 34.4±8.4% and 44.2%, respectively. In total, we detected 285,015 insertions in common for both libraries, while 93,344 and 75,538 insertions were uniquely detected in the P and T libraries, respectively. The combination of both libraries, referred here as ‘PT’, resulted in a total genome coverage of 55.6% (453,897 unique insertions, with an average of 267.1 reads per insertion), and 92.4% when considering a set of known NE genes [29] (20,385 unique insertions along 22,047 bp; Fig. 1B). This represents ∼1 transposon insertion at every base. As expected, the high LDs observed at P1 and T1 samples decreased with increasing rounds of growth selection, highlighting the loss of mutants with reduced fitness (Fig. 1B).

### Gene essentiality estimation in highly saturated transposon libraries

To assign essentiality categories to genes annotated in *M. pneumoniae* (Supplementary Data S2), we used an unsupervised clustering approach based on a Gaussian Mixture Model (GMM) [29]. For this, we used the combined datasets obtained at passage 1 (PT1) since they had the maximal LDs. The GMM analysis highlighted a model fitting in 4 categories (Fig. 1C and Fig. S3). As expected, the LD distributions showed two dominant gene populations in the left- and right-ends representing E and NE, respectively. The high coverage of the datasets made possible a classification of the population in the middle (defined as F genes) into two categories, referred herein as F1 (quasi-E) and F2 (quasi-NE).

Considering all the annotated regions of the genome including protein-coding genes (707), functional RNAs (6), and tRNAs (37), we found a total of n_E_=255, n_F1_=90, n_F2_=86, and n_NE_=319 (Fig. 1D, Supplementary Data S4). Functional RNAs such as ribosomal RNAs (MPNr01, MPNr02, MPNr03), MPNs01 (4.5S RNA), MPNs03 (RNaseP) and MPNs04 (tmRNA), they were all essential for survival. Most annotated tRNAs (n=37) were classified as E (n_E_=29, n_F1_=4, n_F2_=2, n_NE_=2), with the exception of MPNt08, MPNt16, MPNt23 and MPNt24 (F1), MPNt28 and MPNt36 (F2), and MPNt26 and MPNt15 that exhibited transposon densities consistent with an NE category (Supplementary Data S2). Although MPNt26 appears annotated as a serine TCG tRNA, sequence analysis using ARAGORN [30] predicts lack of tRNA structure, suggesting it represents a pseudo-tRNA of MPNt25, with whom MPNt26 shares an identity of 89.9%. This is supported as well by the fact MPNt26 is the only tRNA that does not classify as E after eight passages (Supplementary Data S4). MPNt15 seems to be dispensable out of all the tRNAs decoding arginine codons, consistent with the fact that MPNt37 may complement it due to the wobble effect [31], but not the other way around (Supplementary Data S2). A similar explanation seems true for MPNt28 lysine decoding tRNA that can be replaced by MPNt20, and MPNt36 leucine decoding tRNA that could be replaced by MPNt19 (Supplementary Data S4). Overall, these results highlight the accuracy of the transposon data to predict different degrees of fitness contribution. Additionally, we looked at the essentiality of the 186 ncRNAs annotated in *M. pneumoniae* [15]. Of these, 18% and 12% were classified as E and F1, respectively. Except ncMPN336 that is classified as F1 and overlaps with a F2 gene (MPN415), the other E/F1 ncRNAs overlap with E/F1 genes (Supplementary Data S4). Considering ncRNAs are mostly NE and originate from transcriptional noise in *M. pneumoniae* [32], these overlapping ncRNAs are likely to be dispensable for the cell as well. Regarding protein-coding genes we found a classification consisting in n_E_=220, n_F1_=86, n_F2_=84, n_NE_=317 (Fig. 1D). Among the 220 E coding genes, 185 were found conserved in the minimal synthetic bacterium JCVI-syn3.0 (531,490 bp; 438 protein-coding genes) [33] (Supplementary Data S4). Also, we found conserved in JCVI-syn3.0 other 165 F1/F2/NE genes of *M. pneumoniae*, 105 of them presenting at least 50% of LD, suggesting that some of these genes could be dispensable in the synthetic genome, or if deleted, lead to a viable organism but with reduced growth (Fig. S4 and Supplementary Data S4).

The assignment of two F categories revealed sets of genes that can be disrupted but with different impacts on the viability of the cell. To further explore this, we measured the apparent decay in LD for each gene along passages. This data was first analyzed by clustering the gene LD trajectories using a k-means algorithm (Fig. 1E and Fig. S5), which revealed at least four different clusters (C1 to C4). These clusters aligned well with the predicted essentiality using the GMM model, in which gene essentiality was predicted from the single time point PT1 (Fig. 1E). For example, cluster C1 was characterized by constant E LDs from passage 2 ahead, and comprised almost exclusively E and F1 genes. In contrast, cluster C4 showed stable LDs up to passage 6, mostly including NE genes. However, this analysis revealed additional layers of complexity for F1, F2 and NE genes. In particular, we identified genes that while starting at similar LDs and having a similar GMM category, exhibited different decays and cluster assignments.

### Establishing a method for quantitative assessment of essentiality

The above observations prompted us to define a quantitative method of essentiality. For this, we calculated the area under the decay curve (AUC) for each gene, as an indicator of the cell fitness contribution of each gene (Fig. 1E). Using this approach we observed a good correlation of AUC independently on the passages selected (Fig. S6). We identified, for instance, 72 F2 and 16 NE genes with high LDs at passage 1, but exhibiting low AUC values (belonging to cluster 1 and 2; Fig. 1E) owing to the low persistence of their insertions. This observation suggests that although these genes are classified as NE/F2 at first passage, they have in fact a major contribution to cell fitness. These decay differences could be explained in part by the fact that NE/F2 genes clustered in C2 encode proteins with higher copy numbers, as compared to genes clustered in C3 and C4 (Fig. S7). These differences in protein abundance could affect the fitness effect of genes after disruption, as a certain degree of cellular function may be still present during the first passages until the protein is diluted and/or degraded (average protein half life in *M. pneumoniae* is above 60 hours) [18,22].

In summary, we present in Supplementary Data S4 two complementary essentiality analysis approaches and metric scores that provide a dynamic quantitative assessment of essentiality. This represents an improvement of classification performance compared to previous studies relying on 2 or 3 static categories of essentiality [9,23], thus providing a more accurate view on the specific contribution of each gene to cell fitness.

### Essentiality analysis of transcriptional and translational regulatory elements

We then examined the essentiality of potential transcriptional and translational regulatory sequences. For this analysis, we took into consideration different genomic elements that define transcriptional and translational units (Supplementary Data S2). These include predicted promoters associated to transcription start sites (TSS) experimentally determined [1,34], predicted intrinsic transcription termination sites (TTS), defined 5’ and 3’ untranslated regions (5’UTR and 3’UTR, respectively), and ribosome binding site (RBS) predicted motifs (see Methods). Regions located between two non-overlapping genes and expressed from the same operon were also considered as possible regulatory regions (named here as inter-genic-intra-operon, or “iGiO”). For the analysis, we discarded elements smaller than 5bp, as well as those overlapping with E/F1 genes or containing repetitive sequences, as these are known to interfere with the transposon mapping [29]. In total, 1,050 out of 3,189 regulatory regions remained for downstream analyses.

The majority of the 1,050 selected regulatory regions analyzed exhibited a transposon coverage consistent with a NE nature in the PT1 library (Fig. S8A and Supplementary Data S5). These results indicate that genes in *M. pneumoniae* do not normally require strict regulation either at the transcriptional or translational level, consistent with the limited transcriptional variation observed for this organism [1]. The GMM model predicted 43 regions consistent with E and F1 categories (Fig. 2A). After applying a more stringent cut-off criteria of LD≤0.4, we kept 25 essential regulatory regions associated with 21 different genes. These genes with apparent essential regulation, including 6 tRNAs, were mostly related to translation and transcription (Supplementary Table S1). The 5’UTR and iGiO elements were the most frequent regulatory features associated with these E regions (Fig. 2A and Supplementary Table S1), suggesting that translation initiation, transcript stability, and/or processing, are important contributor factors to the regulation of these genes [35]. For example, we identified essential iGiO sequences separating rRNAs and tRNA genes, which are probably required for transcript processing and maturation [36]. Similarly, we detected depletion of insertions in the promoter and downstream sequences of the mature tmRNA (MPNs04), consistent with the requirement of these sequences to complement a tmRNA null mutant [37]. We also identified cases of E 3’UTR regions, yet these elements often overlap with other genomic features making it difficult to ascertain their specific contribution (Fig. S9). One example is the 3’UTR of *mpn517* (uncharacterized protein), which overlaps with the 5’UTR of *mpn516*, encoding the RNA polymerase subunit β. We also observed a depletion of insertions in the TTS and 3’UTR regions of *mpn061* (Fig. S9). This E gene encodes the signal recognition particle and is oriented in the opposite direction of *mpn060*, an E gene encoding the S-adenosylmethionine synthetase (MetK). The *mpn061* terminator could play a role in preventing collisions during convergent transcription, and avoid transcriptional interference. Alternatively, sequences present in the 3’UTR may be crucial to modulate mRNA stability or the *mpn061* function [38].

**Figure 2.**
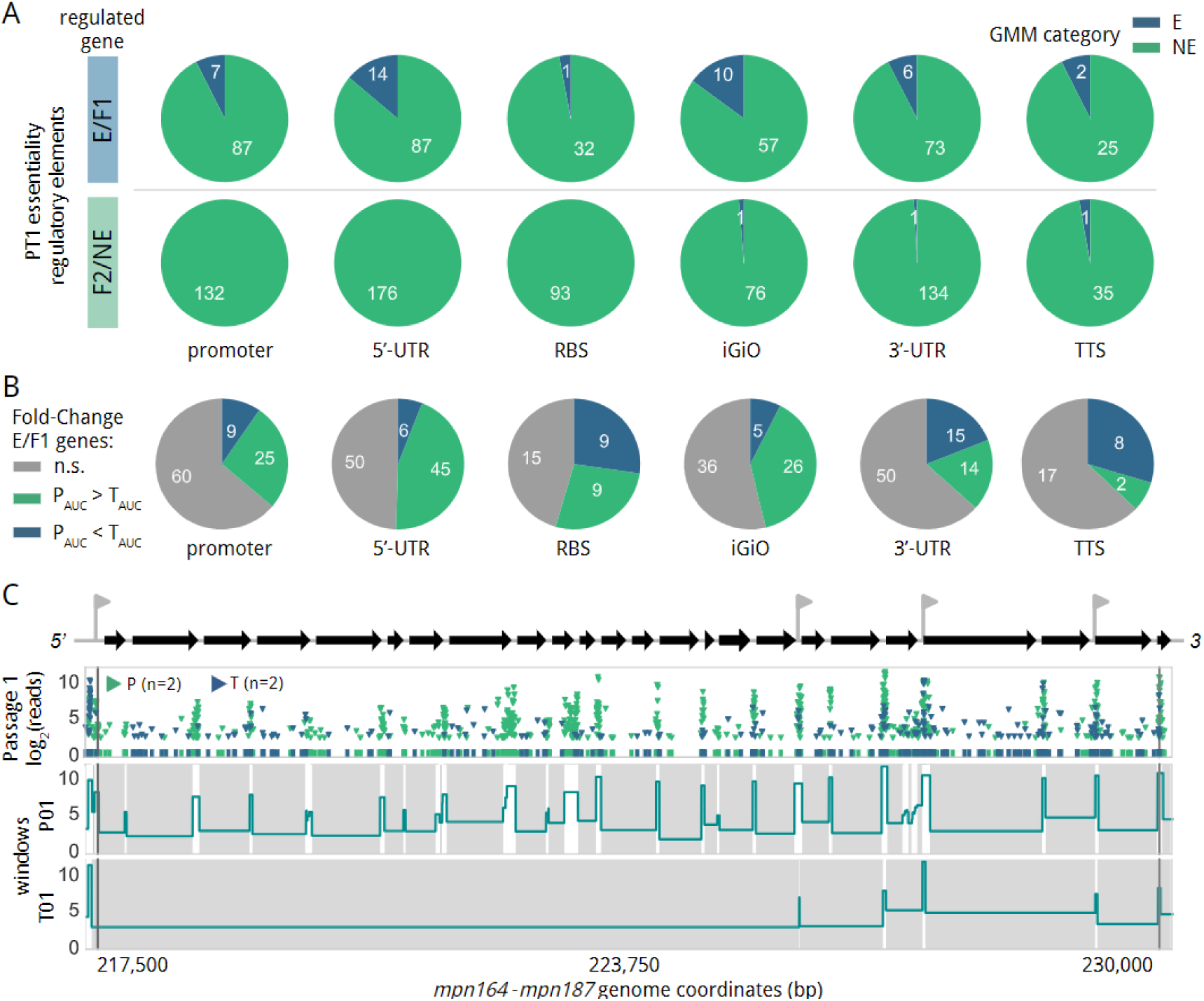
Essentiality analysis of putative regulatory elements. **A)** Essentiality assignment of regulatory elements based on a GMM model. Pie charts represent the number of E (blue) and NE (green) elements for each type of regulatory element. The top row shows regulatory elements associated with E/F1 genes, while the bottom row shows those associated with F2/NE genes. Only elements of at least 5 bp, non-repeated and not overlapping with E/F1 genes are represented. **B)** Fold-changes in AUC comparing the persistence of transposons containing promoters (P) versus terminators (T) in different types of regulatory elements. Pie charts represent the distribution of regulatory elements according to the AUC enrichment tendency. In gray, no significant (n.s), AUC enrichment for P in green, and AUC enrichment for T in blue. **C)** Transposon insertion profile at passage 1 across an operon containing 23 genes (*mpn164* to *mpn187*; top gray arrows) mainly encoding ribosomal proteins. Each transposon insertion and its read count (Y-axis, in log_2_) is represented by an inverted triangle, with transposons containing promoters or terminators labeled in green or blue, respectively. The top scheme also shows predicted endogenous promoters (gray arrows). Below the scheme, the window essentiality domain at first passage highlights the differential insertion bias for the P and T libraries at passage 1, (P01 and T01, respectively). Gray and white areas represent E and NE regions, respectively. The height of each domain corresponds to the log2 transform of the average of read count per insertion in each.

Next, we looked for possible enrichments of promoters versus terminators transposon insertions in regulatory regions (Supplementary Data S5). As expected, no significant differences were found between P and T libraries in regulatory elements exclusively associated with F2 and NE genes (Fig. S8B). In contrast, we detected significant and persistent enrichment of promoter insertions compared with terminators in upstream regulatory regions of genes belonging to E and F1 categories, including promoter, 5’UTR and iGiO sequences; and also for RBSs at later passages (Fig. S8C). We did not find statistically significant differences between the two libraries in TTS or 3’UTR regions, in which terminator enrichments could be expected (Fig. S8B and S8C). However, when looking at the persistence of the insertions across passages, our AUC analysis showed that while promoters persisted longer than terminators in regulatory elements upstream of E/F1 genes, this trend was inverted in 3’UTR and intrinsic terminators (Fig. 2B). These results suggest that transcription termination in *M. pneumoniae* has a minor influence in cell fitness, but could be important for achieving optimal growth. One example is the enrichment of transposons containing terminators in the endogenous *mpn625* terminator sequence (Fig. S1A), consistent with the fact that overexpression of the MPN626 ortholog in *M. genitalium* is toxic [39].

We also detected a particular preference of P library insertions close to the TSS and N-terminus of E protein coding regions at nucleotide level (Fig. S10). As an example, Fig. S2C shows a large transcription unit mainly constituted by E ribosomal genes, in which intergenic regions mainly accept transposons containing promoters. Remarkably, this pattern was also almost completely maintained up to the 6th passage (Fig. S11). In contrast, we found that terminator sequences are better tolerated in the intersections delineating sub-operon structures, where internal promoters can rescue transcription (Fig. S12A).

Despite these general observations, our genome global analysis revealed that 44.7% of 5’UTR (n=105) and iGiO elements (n=39) that are associated with E genes, and lack clear downstream promoters to rescue transcription, tolerate insertions of the T library with relatively high AUC values (≥2). These observations suggest either that transcription termination is leaky depending on the context [40], or that weak unpredicted promoters allow the expression of these genes. In fact, the *M. pneumoniae* sigma 70 factor recognizes a lax consensus promoter sequence, being the Pribnow box the only required element [15]. Furthermore, we found that terminator sequences inserted in regions lacking apparent mechanisms of transcriptional rescue tended to be associated with highly expressed transcripts (Fig. S12B), suggesting that transcription termination may not be totally efficient in these particular cases, as we have shown experimentally (Fig. S1B). Interestingly, we also observed a progressive and fastest depletion after subsequent passages of the transposons located closer to the 5’ end of polycistronic mRNAs, indicating that disturbances occurring earlier in the transcript induces growth defects to a greater extent (Fig. S11 and Fig. S13).

Altogether, the results show that *M. pneumoniae* generally lacks essential regulatory features and its transcriptome is quite resistant to genomic perturbations. Despite this, there is still a need for preserving transcription and translation of E genes when no additional regulatory elements can rescue their expression after disruption. In addition, the results also highlight that the maintenance of proper levels of gene expression is important to achieve optimal fitness.

### Mapping local-level essentiality using an annotation-independent approach

To expand our essentiality analysis to other non-annotated regions of the genome, we used an annotation-independent approach based on a sliding window analysis. For this, we examined transposon insertion densities of informative sliding window fragments of 31 bp. This analysis segmented the genome in 4,255 regions, defining contiguous E segments surrounded by NE (Supplementary Data S6). After filtering out segments containing repeated sequences and shorter than 5 bp (see Methods), we detected in the PT1 library samples a total of 914 E segments with an average size of 328 bp, extending to a maximum of 5,594 bp (Supplementary Data S6). The majority of these DNA segments (n=879) overlapped with annotated coding regions, especially F1 and F2 genes, consistent with the discontinuous insertion profile typically observed in these genes (Fig. 3A, Supplementary Data S7). As expected, the longest E fragments mainly overlapped with E genes and tended to delineate precisely their sequence boundaries as seen for the aforementioned ribosomal operon (Fig. S11). This highlights the accuracy of the algorithm to define E regions and its usefulness to assist in the annotation of genomes.

**Figure 3.**
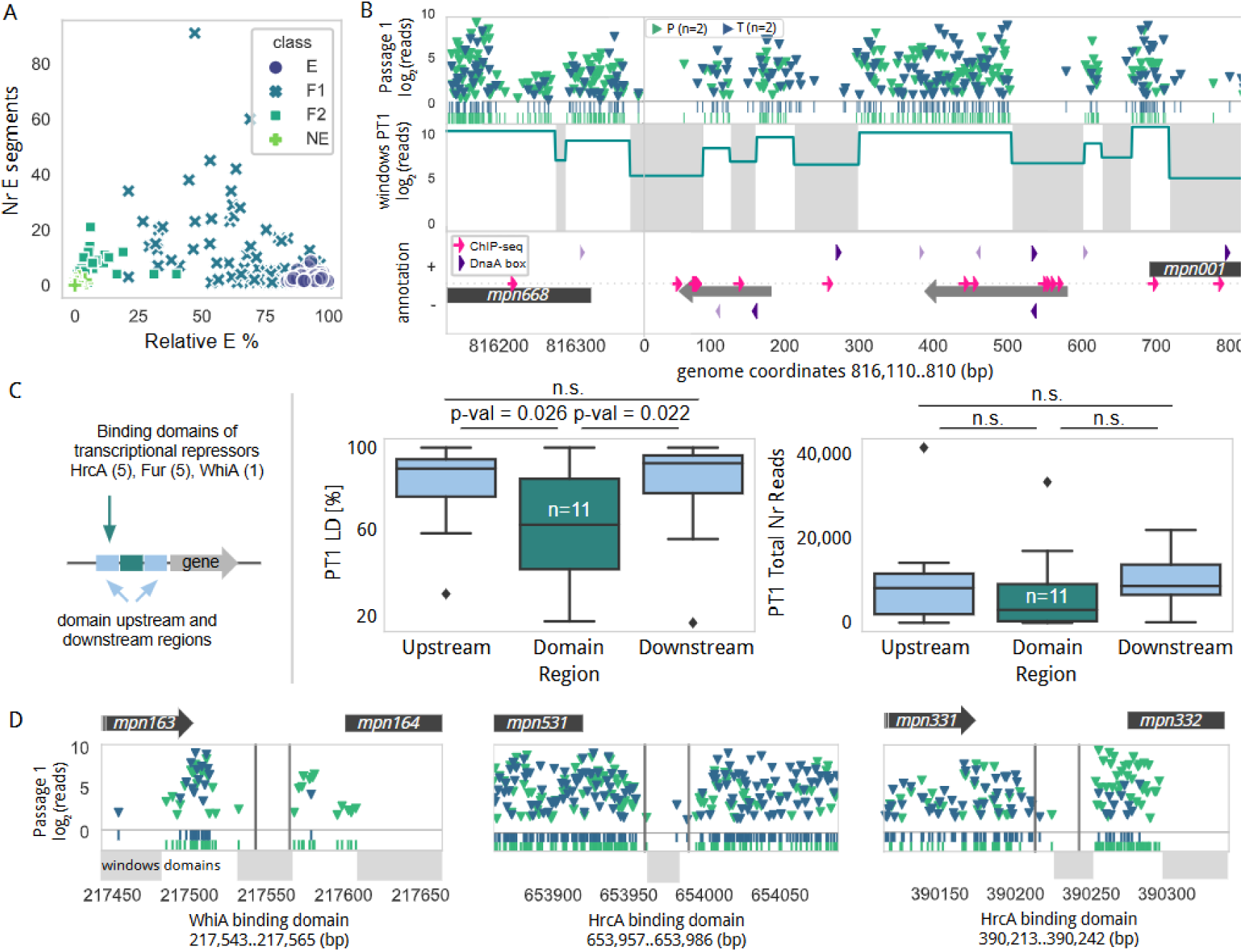
Local-level essentiality analysis using a sliding window approach. **A)** Scatter plot relating the percentage of each gene being essential by our sliding windows approach (X-axis) and the number of E segments (Y-axis) with different coloring and shape based on the essentiality category assigned in the GMM analysis. **B)** Transposon insertion profile in the promoter (P) and terminator (T) libraries at passage 1 (PT1) for the *oriC* genome region. Transposon insertions and read count (Y-axis, in log2) are represented (promoters or terminators labeled in green or blue, respectively). Below, the window essentiality domain highlights the differential segments (gray - E and white - NE). Last row shows the genomic context in both orientations, with gray arrows representing ncRNAs and black representing the N-termini of *mpn001* and *mpn668*. ChIP-seq peaks are represented as pink arrows while the purple triangles label putative DnaA boxes that are darker when overlapping an E domain. **C)** Left - Schema of the analysis performed. Center - Box plot analysis of the preferential insertions using the PT1 library in the domain region where a transcription repressor is bound (n=11), compared with the equivalent size upstream and downstream regions. Right - same representation including the total number of insertion reads. Underlying data points are all included in Supplementary Data S9. **D)** Transposon insertion profile as shown in panel A for DNA binding regions of transcriptional repressors. Windows essentiality domains (gray - E and white - NE) are represented below.

Consistent with the compact genome organization of *M. pneumoniae*, we only detected 35 E segments mapping to non-coding regions. Of these, 30 overlapped with putative gene regulatory elements, while 5 were mapped to other non-annotated areas. Of note, 4 of them are located in a non-coding region that separates *mpn688* (*soj*) and *mpn001* (*dnaN*) genes (Fig. 3B), which apparently contains the origin of replication (*oriC*) of *M. pneumoniae*, as this region together with short stretches of the flanking genes support replication of suicide plasmids [41]. Strikingly, only 5 out of the 10 putative DnaA boxes predicted in the *oriC* were found to overlap with E segments [41], suggesting that many of these predicted DNA boxes are either redundant or not functional, or the requirements for initiation of replication in *M. pneumoniae* are rather relaxed. In fact, two of the E DnaA boxes lie within the E coding sequences of *soj* and *dnaN*, thus making it difficult to ascertain their real essentiality. The discontinuous E profile of the *oriC* region could reflect the importance of the specific arrangement of individual DnaA boxes and their different affinity properties. In other words, insertions between specific DnaA boxes could alter the local distance, indirectly affecting the nucleation process and proper assembly of DnaA oligomers [42]. Some of the identified E segments may also be important for binding other essential proteins.

### Impact of genome accessibility on Tn-Seq data

Next, we wonder whether the insertion profile of small chromosome regions are affected by the binding of DNA proteins, which could prevent transposon accessibility. To evaluate this effect, we analyzed regions of the genome where we detected bound proteins as revealed by ChIP-seq and protein occupancy data [1]. In general we did not find differences in terms of the number of transposon insertions in these regions, but we found some exceptions (Fig. S14 and Supplementary Data S8). For example, two of the DNA segments in the *oriC* region found to be E could be protected as shown by DNA protection studies (POD) [1], or specifically methylated [21]. Additionally, ChIP-seq showed binding between nucleotides 1 and 100 for the SMC-ScpAB complex (crucial role in the organization and segregation of the chromosome [43]), the RNA polymerase complex (around position 550), and DnaC (around position 650) [1]. Thus we see that the E regions detected in the *oriC* seem to be bound by different protein complexes. This raises the question of whether some of these small E regions detected are truly essential or if they are merely protected from transposon insertion (see below for further discussion). To further explore whether DNA protection could affect transposition, we looked at the promoters regulated by the three known transcription repressors of *M. pneumoniae* (HrcA, Fur and WhiA) [1], as these are likely to bind in a more constitutive manner to their DNA target sites and some of them regulate NE genes. Inspection of a total of 11 binding sites for these transcriptional repressors, shows in general a significant reduction in LD and a decreased trend in total transposons reads, when compared to adjacent regions of the same length (Fig. 3C, Supplementary Data S9). This is especially clear for the main targets of HrcA (*mpn021*, *mpn332* and *mpn531*), for two main targets of Fur (*mpn043* and *mpn162*) and the one of WhiA (*mpn164*) (Fig. 3D). Of note, we have previously demonstrated that the upstream regulatory region of *mpn332* where HrcA binds can be replaced by an inducible system platform [22], demonstrating that this region is in fact NE under normal growth conditions as long as there is an active promoter transcribing *mpn322*, which encodes the essential Lon protease. The fact that *mpn531*, *mpn043* and *mpn162* encode NE proteins, also supports the conclusion that these regulatory regions are NE and that the Tn-seq data is sensitive to strong DNA-protein interactions.

Overall these results suggest that lack of transposon insertions in regulatory regions could be in some cases due to the tight binding of regulatory proteins. Therefore, genome accessibility must be considered when interpreting Tn-seq essentiality data of small regions.

### Detection of non-essential regions and domains in essential protein-coding genes

Next, we exploited our sliding-window analysis to identify possible NE regions within E and quasi-E (F1) protein-coding genes. We found that E genes tolerate insertions affecting 5 and 9 amino acids in average at the N- and C-terminal regions, respectively (representing 2% and 4% of the total protein, Supplementary Data S7). For F1 genes the tolerance increased to 13 and 18 amino acids (7% and 9%; Fig. 4A). Sequence conservation analysis of orthologous proteins in closely-related species revealed that in *M. pneumoniae* out of 345 E/F1 genes, 193 of them have acquired N- or C-terminal extensions through evolution (Supplementary Data S7). Notably, ribosomal proteins were highly represented among this group (Fig. S15), perhaps reflecting strategies for ribosome evolution. Although the selection process underlying these protein extensions is unclear, our data indicate that they are disruptable or at least less essential than the rest of the coding gene. For example, we detected within the *nusA* gene of *M. pneumoniae* (*mpn154*), a high tolerance of insertions across the 3’ end of this E gene, corresponding to a long C-terminal extension of 124 amino acids (Fig. S16A). As expected, when analyzing P and T libraries separately, transposons carrying promoters identified predicted longer N- and C-terminal (in the case of genes preceding an E or F genes in the same operon) NE extensions for both E and F genes (Fig. 4B). For example we o found cases like *mpn229* (Fig. S16B) and *mpn116* (in this case, a F2 gene; Fig. S16C), in which the NE extensions are covered almost exclusively by transposons containing promoters, which highlights the importance of preserving transcription of downstream genes. Importantly, although the protein extensions identified are apparently NE, our results indicate that the persistence of insertions in these regions is generally lower than those found in NE coding genes (Fig. 4C). These observations support the notion that these NE extensions do have a contribution to cell fitness in many cases.

**Figure 4.**
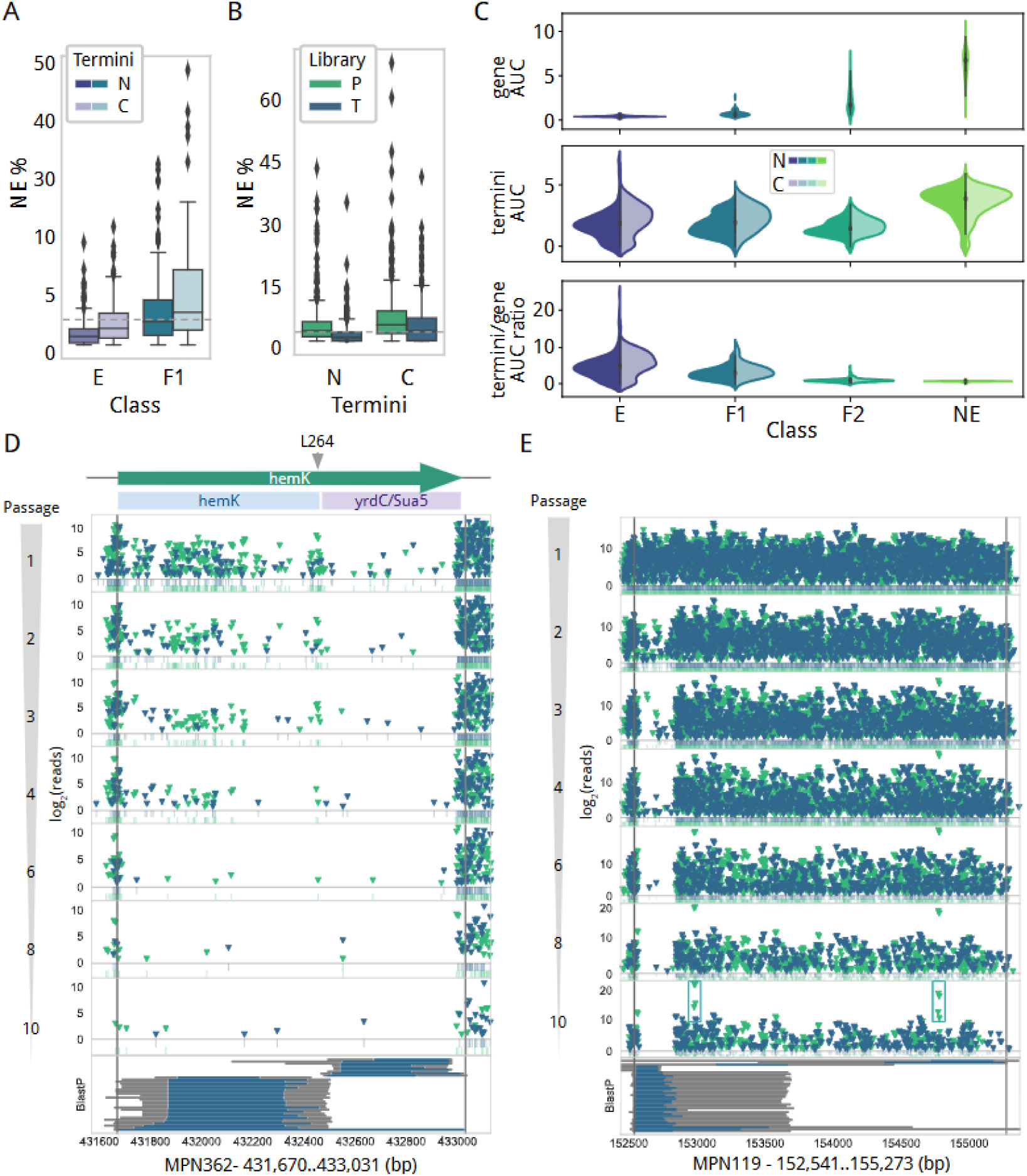
Analysis of regions with different essentiality in coding genes. **A)** Box plot comparing the percentage of E and F1 genes being disrupted by insertions when combining P and T libraries at first passage (PT1). The darker box indicates the percentage of the protein at the N-terminus that is found to be NE and the clear box the same for the C-termini. Notice a dashed gray line marking the 5%, generally used as threshold for permissive insertions in essentiality studies. **B)** Same representation as in C panel but combining E and F1 genes and coloring by the library (green for promoter (P) and blue for terminator (T), respectively). **C)** Violin plots comparing the AUC calculated for genes (top), N- and C-termini (middle) and the ratio of these two values (bottom) by essentiality category from the GMM model. In the termini and ratio panels, the solid (left) and transparents (right) colors of the splitted violin plots represent the AUC and ratio distributions calculated for N- and C-termini, respectively. **D)** Transposon insertion analysis of MPN362 reveals two domains with different essentiality (hemK and yrdC/Sua5). Insertion profiles for promoter (green) and terminator (blue) from passage 1 to 10 are displayed. Last row represents the BlastP results, in blue the aligned region and grey non-aligned sequence for the homologous hit. It can be noticed the two domains are commonly found splitted in other bacteria. **E)** Same representation as in panel D for MPN119, which encodes a DnaJ-like protein associated with the attachment organelle. Supported by BlastP analysis, it is shown a DnaJ-like conserved region at the N-terminus that accepts transposon insertions during the first two passages, but these are rapidly lost after consecutive passages, indicating a different fitness contribution as compared to the rest of the gene. Interestingly, two promoter regions (represented in the green boxes at passage 10) are selected with high efficiency, suggesting split variants with enhanced fitness. From the BlastP analysis, hits mapping each of the three splits suggested by promoter insertions can be retrieved in other bacteria. Note passages 8 and 10 Y-axis are in a different scale with insertion reaching a log_2_(reads) value over 20.

Then, we focused our attention on genes containing protein domains exhibiting distinct profiles of essentiality that could suggest modular functionality within the protein. Through visual inspection of the transposon data, we defined domain boundaries from a total of 16 genes that contained regions larger than 15% of the total protein length, with a transposon profile different from the rest of the coding sequence (Supplementary Table S2). Of note, many of the identified regions corresponded to known conserved domains assigned to COG or Pfam families. An example is *mpn362*, which encodes a N5-glutamine methyltransferase (HemK) and a C-terminal protein domain conserved in the YrdC/Sua5 protein family, which is responsible for the formation of the threonylcarbamoyladenosine (t(6)A) modification found at position 37 of ANN decoding tRNAs [44]. HemK in *E. coli* methylates the release factor RF1 and RF2 [45]. This post-translational modification has been shown to induce an important stimulatory effect on the release activity of RF2 [46]. Due to its unusual genetic code, *M. pneumoniae* only contains the RF1 protein that is encoded by the gene located upstream of *mpn362*. We found that the HemK domain tolerated transposon insertions while the YrdC/Sua5 domain remained mostly free, indicating that the essential nature of *mpn362* (classified as F1) is probably due to the activity of the YrdC/Sua5 domain (Fig. 4D and Supplementary Table S2). To validate these results, we used the SURE-editing genome-engineering tool [47] to replace the endogenous *mpn362* gene by N-terminal deletion variants containing only the YrdC/Sua5 domain (Fig. S17A). We identified two possible in-frame start codons close to the junction of both domains: TTG and ATG, encoding respectively residues L264 and M290. To express the YrdC/Sua5 domain alone, we tested both translation start codons under the control of the P438 promoter. Only positive gene replacements with the construct using the TTG start codon was obtained (Fig. S17B), consistent with the enrichment of transposons containing promoters detected just before this alternative start codon (Fig. 4D). Then, we assessed whether MPN362 is expressed as a domain fusion protein only, or YrdC/Sua5 can be expressed as a separate domain as well. To test this, we added a FLAG tag at the C-terminus of MPN362 and replaced the endogenous gene with this tagged variant (Fig. S17B). This FLAG-tagged variant was viable, and expressed only detectable levels of the full-length isoform based on Western blot analyses using anti-FLAG antibodies (Fig. S17C). Altogether, these results demonstrate that the Hemk domain of *M. pneumoniae* is dispensable for cell viability, yet it is expressed as a fusion protein with an essential YrdC/Sua5 domain. These results suggest that either an alternative modifying enzyme of RF1 exists, or *M. pneumoniae* RF1 may function independently of post-translational modifications.

Although differential domain essentiality was generally detected at passage 1, we also found cases in which this distinct behavior was not observed after several passages. For example, *mpn119* that encodes the ortholog of the MG200 gliding motility protein of *M. genitalium* [48,49] is fully disruptable across the whole coding sequence at passage 1. However, we detected a decrease in LD affecting only the first 100 amino acids, reaching a full essential LD profile at passage 3 (Fig. 4E). This N-terminal region encodes for a conserved DNAJ-like domain, indicating that in addition to the role of MPN119 in gliding motility, this domain may have an important contribution to cell fitness as well. However, we cannot rule out the possibility that the expression of a putative small ORF of 68 amino acids within this region may be responsible for the distinct essentiality profile observed, albeit we did not identify peptides by MS analysis [50,51]. Overall, these observations highlight the importance of assessing essentiality at longer growth time points.

### Structural essentiality and discovery of proteins that can be split

We then set out to characterize small NE internal regions of E coding sequences from which we could infer structural protein information. We identified 9 E genes that seem to maintain functionality when expressed apparently as split proteins (Supplementary Table S3). One of such examples is *mpn106*, which encodes the β subunit (PheT) of the Phenylalanyl-tRNA synthetase (PheRS). This essential enzyme catalyzes the covalent attachment of Phe to its cognate tRNA and has a complex tetrameric organization composed by two α subunits (PheS) and two β subunits in a (αβ)_2_ configuration [52]. As expected, both *mpn105* (*pheS*) and *mpn106* (*pheT*) exhibited a clean essential transposon profile across the gene length, except for a segment of about 34 bp in *pheT* that tolerated transposons containing promoters (23±5 insertions with 129±16 read count average; n=2), but rarely accepted transposons containing terminators (3.5±1.5 insertions with 10±5 reads; n=2). Notably, mutants with these transposon insertions were still tolerated after consecutive passages (Fig. 5A). Sequence alignment analysis indicated that this disruptable region coincides with a linker that separates the structural domains B1 and B3 (Fig. 5B) [52]. Inspection of this region revealed 3 possible in-frame alternative translation start sites, at residues L196, L202 and M208, suggesting that PheT may still be functional when expressed in two separate protein fragments. To corroborate these observations, we constructed a split *pheT* variant in which we introduced a premature stop codon after residue T207, followed by the P438 promoter sequence to drive the expression of a second PheT fragment starting at residue M208 (Fig. 5C). This would result in the expression of an N-terminal fragment of 207 amino acids corresponding to the structural domain B1-B2-B1, and a second fragment of 598 amino acids encompassing domain B3 to B8. Using the SURE-editing genome-engineering tool [47] we successfully replaced the endogenous *pheST* locus by the mutant variant, demonstrating that *pheT* can be split in two protein fragments without compromising *M. pneumoniae* viability, suggesting that the enzyme retains activity (Fig. 5D). Of note, the β subunit of PheRS contains recognition and binding sites of the tRNA [53,54], suggesting that the split fragments are capable to structurally accommodate the tRNA and interact with the catalytic α subunit.

**Figure 5.**
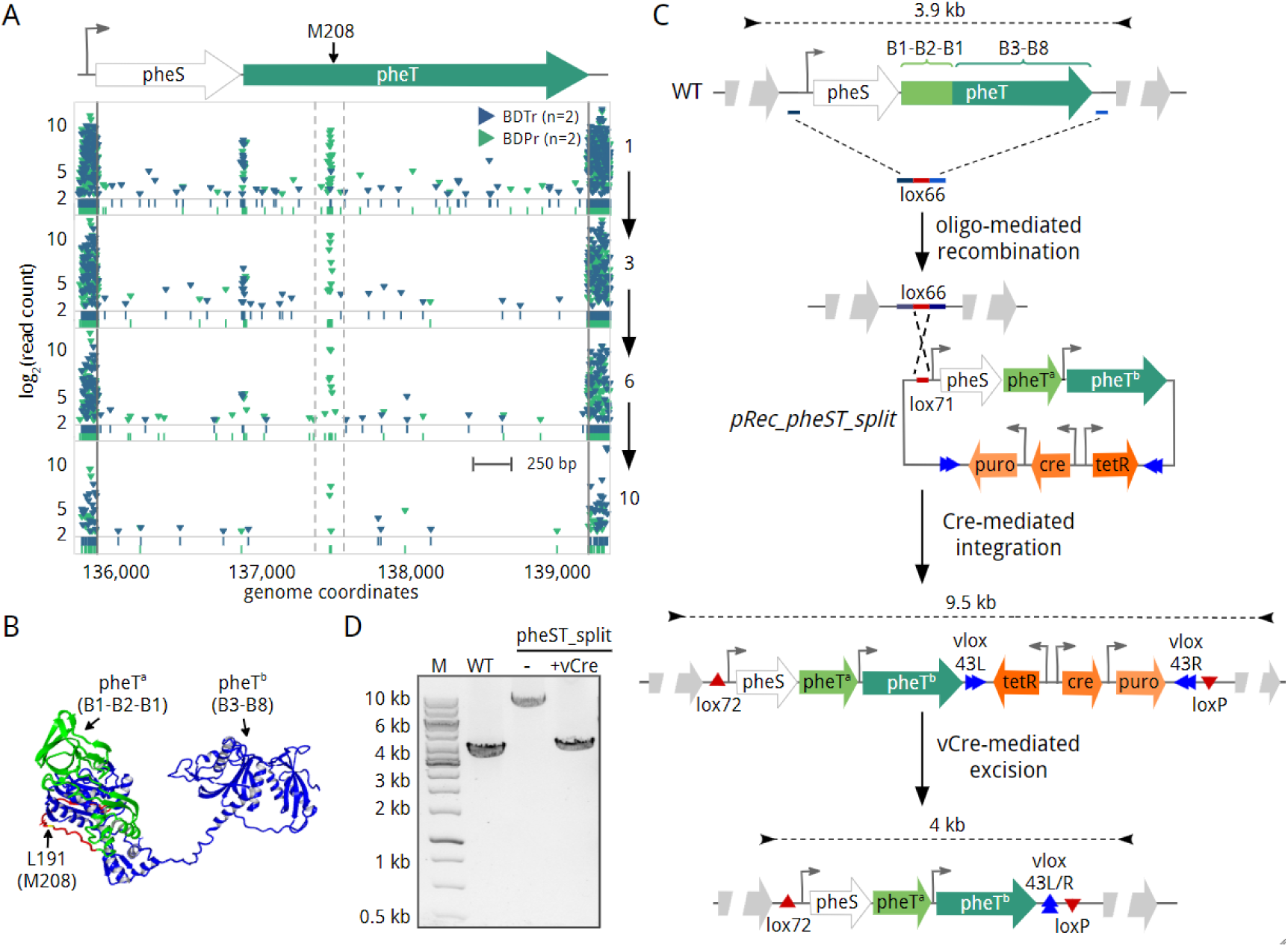
Identification and validation of a functional split protein. **A)** Transposon insertion analysis of the *mpn105* (*pheS*) and *mpn106* (*pheT*) genes. The four rows show the transposon insertion profile (green for promoter (P) and blue for terminator (T)) across these E genes at different passages (Left Y axis). The narrow region bounded by gray discontinuous lines shows preferential insertion of the P library upstream the M208 residue of the *pheT* coding-sequence, which could serve as an alternative translation start codon. **B)** Three dimensional structure of the phenylalanyl-tRNA synthetase beta subunit (PheT) from *Thermus thermophilus* (PDB 1PYS). Domains B1-B2-B3 (in green) and B3-B8 (in blue) are separated by a linker (in red) that accepts preferential insertions of the P library. Within this linker, we show residue L191 (in yellow), which corresponds to residue M208 of *M. pneumoniae* PheT protein based on sequence homology. **C)** Schematic diagram showing the constructs and SURE-editing procedure used to generate a *M. pneumoniae* strain with a splitted PheT variant. Briefly, deletion of the *pheST* endogenous locus is mediated by oligo-recombineering using an oligo containing a lox66 recombination site (in red) flanked by homologous regions (in blue). Gene complementation with the mutant variant is then mediated by Cre-mediated integration of plasmid pRec_pheST_split in the lox66 site, generating lox72 and loxP sites. Plasmid backbone (containing tetR repressor, Cre and puromycin resistance marker) is then removed from the genome by vCre-mediated recombination using the vlox sites present in the pRec_pheST_split plasmid sequence. Sizes of the PCR products expected for each intermediate strain are shown above. **D)** PCR analyses using genomic DNA of the WT and intermediate strains before (labeled as “-”) and after (labeled as “+”) vCre excision are shown. The size of the expected PCR products is shown in panel C.

These results support the suitability of transposon mutagenesis to identify structural regions and guide the design of protein-fragment complementation assays.

### Assessment of transposon insertion events improving cell fitness

The transposons used in this study are expected to change the transcription strength of the genes surrounding the transposon insertion. Although the majority of these transcriptional perturbations are expected to negatively impact cell fitness, we wonder whether there may be cases promoting an improvement. To identify these cases we computed the enrichment of normalized read values for both P and T libraries between passages 1 and 10. In order to constrain the analysis, only reproducible insertions between replicates were considered. We identified a total of 958 and 1605 positions whose read counts were significantly enriched from passage 1 to 10 exclusively in the P or T libraries, respectively. An additional set of 29 positions belonged to both libraries (Fig. 6A and Supplementary Data S10). To explore the potential effect of these insertions, we mapped them based on their distance and position to the corresponding gene and non-coding annotations As expected, the majority of enriched insertions overlapped to F2/NE genes (71.4% in P and 73.8% in T; Fig. S18). We then analyzed the RNA expression of the 147 genes showing enrichment of P transposon insertions in upstream positions (Supplementary Data S11). In general, we found that these genes presented lower expression, when compared to those genes lacking promoter enrichments in P transposon insertions (Fig. 6B). This observation may be indicative that the strong P438 promoter is selected in those positions where it could lead to an increase in expression that could improve fitness.

**Figure 6.**
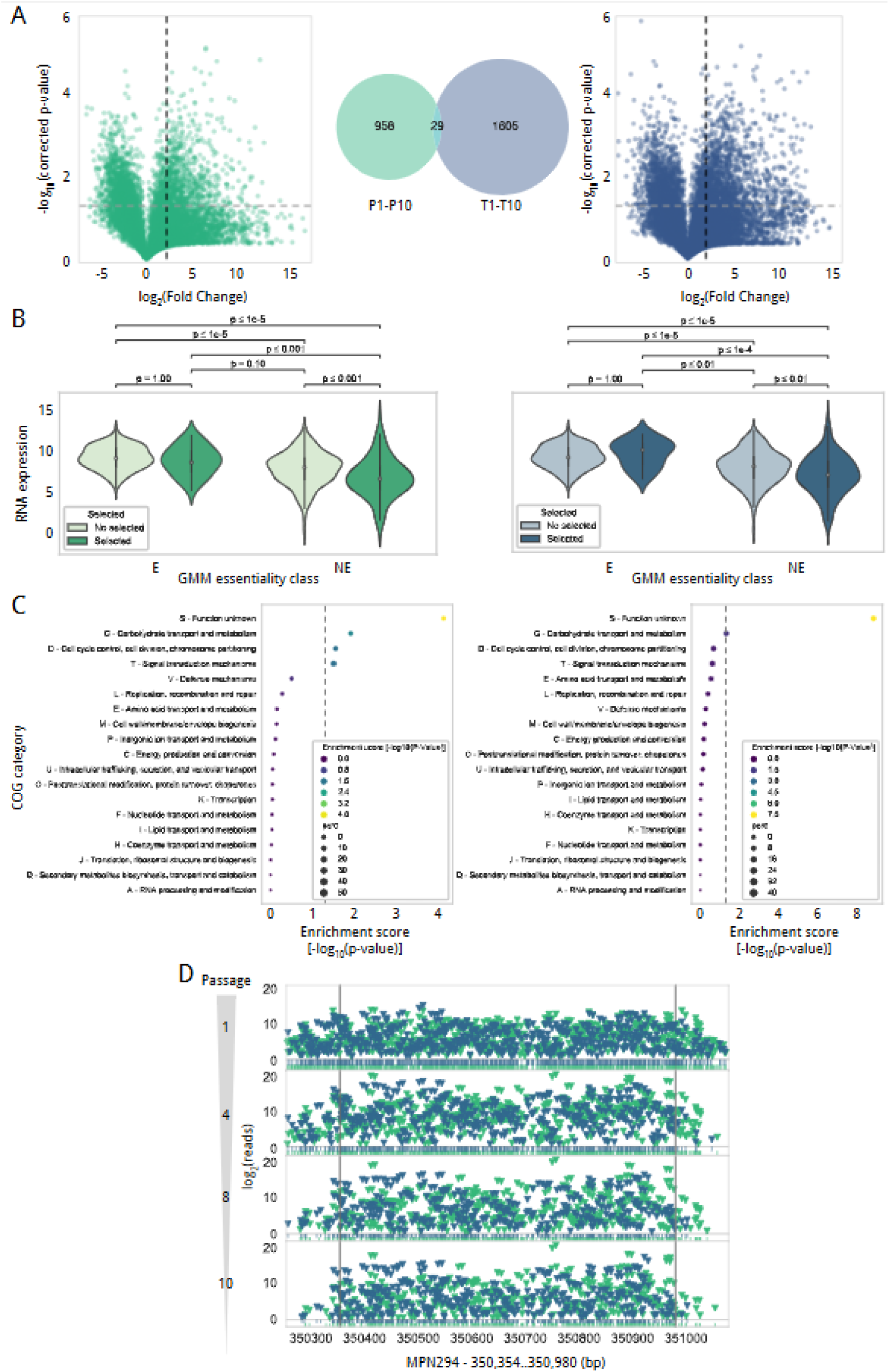
Transposon insertion enrichment analysis after 10 serial passages. **A)** Statistical analysis to identify insertion positions in the genome that are enriched after serial passages. We show the volcano plot analysis for the P (green) and T (blue) libraries (see Methods) as well as the number of positions enriched in both libraries and their intersection as a Venn diagram. **B)** RNA expression of the 147 genes showing enrichment of P transposon insertions in 5’upstream positions (green) compared with those not having it (blue). Genes with an insertion selected present a significantly lower expression, when compared to those genes lacking promoter enrichments in P transposon insertions (all statistical comparisons tested by two-tailed T-test). On the right, the same comparative for the T transposon library. **C)** Enrichment in COG categories for genes showing enrichment of P transposon insertions in 5’upstream positions for the P and T libraries (left and right, respectively). **D)** Transposon insertion profiles in *mpn294* gene across passages for promoter (green) and terminator (blue) libraries. Passages 1, 4, 8 and 10 are displayed. Note passages 8 and 10 Y-axis are in a different scale with insertion reaching a log_2_(reads) value over 20.

In parallel, we explored the function of these genes by means of a COG enrichment analysis (see Methods). We found an enrichment in genes belonging to the COG categories G - carbohydrate transport and metabolism, D - cell cycle control, cell division, chromosome partitioning; T - signal transduction mechanisms; and S - unknown function. When doing this analysis for genes showing enrichment of insertions containing terminators in upstream positions, we only detected COG enrichment for the S category (Fig. 6C and Supplementary Data S12). Apart from transcriptional perturbations, we also looked for gene disruptions that could improve cell fitness. In the absence of sampling effects during the passing process, we would expect in this case an increase of transposon reads across the whole coding sequence, instead of enrichments in specific positions. To discard cases where an increase of transposon reads could be derived from a few insertions, we subtracted the three insertions with higher reads at passage 10. After applying these criteria, only *mpn294* with homology to proteins belonging to the DJ-1/PfpI superfamily, was found to contain a consistent enrichment in transposon insertion reads across the whole coding sequencing (Fig. 6D).

Overall, these results highlight the power of transposon analysis to infer possible strategies to adjust cell fitness during the design of a bacterial chassis.

## DISCUSSION

This study presents a comprehensive analysis of transposon sequencing data using engineered transposons containing promoter, or terminator sequences, to explore complex genomic features. On one hand, we provide the essentiality map of a bacterial genome with the maximum coverage achieved in a study of this kind, close to 1 nucleobase resolution for NE genes. With this unprecedented high coverage, and using unsupervised clustering methods of analysis in combination with ten serial passages, we show a complex status of gene essentiality distant from simplistic models relying on static and binary classifications. This supports the concept that gene essentiality is a conditional trait that depends on the genetic and environmental context [55], in our case on the number of cell divisions at which essentiality is determined. The high transposon coverage obtained in this study allowed us to establish four essentiality categories based on a standard GMM model, corresponding to essential (E), non-essential (NE), quasi-essential (F1) and quasi-nonessential (F2). When we looked at the persistence of the transposons across passages, we defined 4 clusters that in general aligned well with the GMM essentiality categories. However, we noted that some genes exhibited different profiles of transposon persistence despite having the same essentiality category based on the GMM model. These results indicated that genes initially predicted with the same essentiality category can have in fact distinct degrees of fitness influence. Indeed, our analyses suggest that the initial concentration of the gene products can be a possible contributing factor explaining these differences.

These observations motivated us to implement an alternative estimation method of essentiality, based on the integration of temporal datasets by computing transposon decay curves over time. This approach offers a dynamic and quantitative estimation of the impact of loss or gain of function, thus enabling to distinguish the contribution of each gene, including those that are apparently dispensable for cell growth. This quantitative information can be particularly useful for engineering purposes and optimization of cellular processes. Additionally, this methodology could potentially provide insights on the roles of genes with unknown function, by comparing transposon persistence and selection across different genetic and environmental conditions [56–59].

Taking advantage of the 1-base resolution of the produced transposon data, we applied this methodology to perform a systematic interrogation of a bacterial genome to also infer essentiality information for less explored genomic features. To this end, we applied a sliding-window basis analysis to detect highly essential sequences across the genome [13]. Probably as a consequence of the compact genome organization of *M. pneumoniae*, we did not find many essential elements in non-annotated regions except defined sequences in few regulatory regions, mostly related to translation and transcription, and in the origin of replication. However, the essentiality of these small regions should be considered with caution since we found that tight binding of proteins to DNA can prevent transposon insertions. As an example, regions like the CIRCE element of the Lon protease (MPN332), which is targeted by the HcrA repressor, is wrongly assigned as essential in standard growth conditions based on the transposon sequencing data. Thus, these data should be integrated with ChIP-seq or POD data to clearly distinguish essential possible regulatory regions from those that cannot be defined without further experimentation.

When we compared the promoter and terminator libraries, we detected significant and persistent insertions of the P library in upstream regulatory regions of genes belonging to E and F1 categories, including promoter, 5’UTR and iGiO sequences; and also for potential RBSs at later passages. In contrast, we could not see a general enrichment of insertions of the T library at predicted TTS and 3’UTR regions. However, there are few examples where we clearly see preferential insertions of the terminator library at a TTS, like in the case of the TTS of *mpn625,* which prevents expression of MPN626. The importance of this termination site is consistent with the fact that overexpression of the MPN626 ortholog in *M. genitalium* is toxic [39]. When looking at the persistence of the insertions across passages, we found that while insertions from the P library persisted longer than those of T library upstream of E/F1 genes, this trend was inverted at 3’UTR and predicted TTS. This again illustrates the importance of analyzing multiple passages when analyzing essentiality of genomes and suggests that transcription termination in *M. pneumoniae* has a minor influence in cell fitness, but it is important in some genetic contexts for achieving optimal growth. It is also important to emphasize that while as described above we see very few cases for preferential insertions of the T library, we see many cases where there is a negative selection of terminators while P insertions are maintained. One strikingly clear case is the main and essential ribosomal operon, thus showing that the terminator sequence introduced in the T library transposon is functional.

When looking at annotated regions, we identified some protein-coding genes exhibiting different levels of essentiality across the coding sequence, suggesting they likely represent multifunctional genes that originated from the fusion of separate protein domains, or from elongation of the coding sequence. We could define non-essential N- and C-terminal extensions of essential genes, which were found to be quite common among ribosomal proteins. Although these extensions are not required for cell survival, our analysis across passages suggests that their presence confers in many cases a certain fitness advantage. In this regard, it has been suggested that some extensions of eukaryotic ribosomal proteins are involved in recruitment and binding of key initiation factors [60]. Additionally, some ribosomal proteins have acquired other functions beyond the ribosome [61,62], opening the possibility that these extensions may have evolved to fulfill these functions. For example, the transcription elongation factor NusA interacts with the RNA polymerase via its N-terminal domain, but it has been recently shown that it also interacts with ribosomes via its C-terminal region, suggesting that NusA could mediate transcription-translation coupling in *M. pneumoniae* [63]. Although this C-terminal extension was previously established as essential [63], our data indicates it is fully disruptable at passage 1, yet transposons are negatively selected in this region after consecutive passages. This suggests a contribution of this C-terminal extension to protein function, likely enhancing cell fitness. Remarkably, we could also infer structural information of essential proteins with a modular domain architecture, where one of the domains is essential while the other is not, confirming they have arised from gene fusion. An example that we validated experimentally is the *mpn362* gene, which contains a NE (Hemk) and an E (YrdC/Sua5) domain. Thus, this type of analysis shows its application when trying to define a minimal genome since it can identify NE functions present as fusion proteins. Finally, both the inclusion of promoter sequences in the transposon design and the 1 bp resolution of our transposon data, allowed us to identify regions in proteins where they could be splitted while maintaining their functionality. An example that we validated in this study is the Phenylalanyl-tRNA synthetase.

In addition, the use of engineered transposons bearing distinct transcription regulatory elements enabled us to assess the impact of transcription and translation disturbance on cell fitness. We found that gene expression in *M. pneumoniae* is mainly independent of essential regulatory regions and resistant to transposon perturbations. Despite this, the expression of essential genes is preserved in the mutant libraries, and most of the insertions are negatively selected after consecutive passages, emphasizing the importance of maintaining proper levels of expression to achieve optimal fitness. By contrast, we also identified insertions conferring competitive benefits in specific positions of the genome, especially of the promoter library in upstream positions of some low expressed genes, suggesting possible engineering strategies to obtain highly adapted strains. We also found that disruption of *mpn294* was highly selected in our growth competition assay. This gene encodes a protein of unknown function belonging to the DJ-1/PfpI superfamily, which includes some chaperones, proteases, and other stress response proteins [64]. In *Pseudomonas aeruginosa*, *pfpI* defective mutants exhibit higher spontaneous mutation rates and are more sensitive to different types of stress [65]. Thus, it is tempting to speculate that MPN294 disruption may cause increased mutation rates, leading to a larger repertoire of mutants with advantageous fitness rates.

In conclusion, we describe a novel methodology of essentiality assessment using a k-means unsupervised clustering method that in combination with multiple passages provides quantitative information about the contribution of different genetic elements in the overall fitness of a genome-reduced bacterium. The methodology described here can be applied to other model organisms, only limited by the transposon coverage that can be achieved in each particular cellular system. We show that the leverage of Tn-Seq data can be very informative and useful to guide genome annotations, in addition to serving as a resource for fundamental biology research and synthetic biology.

## METHODS

### Bacterial strains and growth conditions

Bacterial strains used in this study are summarized in Supplementary Table S4. Wild-type *M. pneumoniae* lab strain M129 (referred here as M129_LSR) and its derivatives were grown in a modified Hayflick medium at 37°C under 5% CO_2_ in tissue culture flasks. This medium was supplemented with puromycin (3 µg/ml), chloramphenicol (20 µg/ml), tetracycline (2 µg/ml) or gentamicin (100 µg/ml) for selection of transformants, and 0.8% agar when solid medium was required. To induce the Ptet promoter system we used anhydrotetracycline (aTc) at 5 ng/ml. For cloning purposes we used *Escherichia coli* strain Dh5α (New England Biolabs), which was grown at 37°C in LB broth or LB agar plates containing ampicillin (100 µg/ml).

### Molecular cloning

Plasmids used in this study are summarized in Supplementary Table S5 and were constructed by Gibson assembly using the primers listed in Supplementary Table S6 as follows: Vectors pMTnCat_BDPr and pMTnCat_BDter were designed to generate two distinct mycoplasma transposon libraries. Construction of pMTnCat_BDPr plasmid was previously described [29]. The pMTnCat_BDter transposon vector was obtained by amplifying the *cat* gene using ter_cat_F and ter_cat_R primers, and the resulting PCR product cloned by Gibson assembly into a pMTnCat vector [24] opened by PCR using primers p_ter_F and p_ter_R.

To assess the termination activity of the ter625 promoter, we used *cat* and *cherry* gene reporters expressed as polycistronic or fusion transcripts, respectively. To test the polycistronic context, we generated the pMTnTc_ter625_RBS_CAT plasmid that contains the *tetM438* gene followed by the ter625 terminator sequence, a RBS sequence and the CAT coding sequence. To obtain this plasmid we amplified the *cat* gene using two consecutive PCR reactions to include the ter625 and the RBS sequences. For the first reaction we used the pair of primers ter625_RBS_CAT_F and p_CAT_R and then primers p_ter625_F and p_CAT_R. The final PCR product was cloned by Gibson assembly into a digested *Bam*HI pMTnTetM438 vector [26]. We also generated the plasmid control pMTnTc_ter625mut_RBS_CAT, in which we mutated the ter625 sequence to disrupt the terminator hairpin structure. For this, we used the same cloning strategy but using the primer ter625_RBSmut_CAT_F instead of ter625_RBS_CAT_F primer. To test the fusion context, we generated the pMTnTc_P438_MP200_ter625_cherry plasmid. This construct carries an in-frame N-terminal fusion of a 29 amino acid peptide of mp200 ORF to the Cherry coding sequence [37], but separated by the ter625 terminator sequence, which was included in-frame with the fused coding sequences. To obtain this plasmid, we amplified the mp200 peptide using two consecutive PCR reactions, first with the pair of primers p438_MP200_F and ter625_MP200_R, and then with p_P438_F and ter625_MP200_R. The final PCR product was cloned by Gibson assembly into a digested *Eco*RV pMTnTetM438 vector together with the Cherry coding sequence, which was amplified with primers ter625_Ch_F and p_Ch_R. As a control, we generated the pMTnTc_P438_MP200_ter625mut_cherry plasmid, in which we mutated the ter625 sequence to disrupt the terminator hairpin structure. For this, we used the same cloning strategy but using the primer ter625mut_Ch_F instead of ter625_Ch_F primer.

To confirm the viability of a subset of transposon insertions, we performed *M. pneumoniae* genome editions using the SURE-editing tool [47] and plasmids pRec_ΔNt_L_hemK, pRec_ΔNt_M_hemK, pRec_hemK_FLAG and pRec_pheST_Stop_P438. All these plasmids are based on the selector plasmid pLoxPuroCre that allows the selection of edited cells after oligo recombineering [47]. Since the genome editions attempted in this study affected essential regions, we modified the pLoxPuroCre selector plasmid to include essential regions and the intended gene mutations to perform DNA replacements of large genomic regions. For example, the pRec_ΔNt_L_hemK selector plasmid contains the essential genes *mpn360*, *mpn361* plus an N-terminally truncated gene version of *mpn362* (previously annotated as *hemK*), in which L264 was used as start codon and its expression driven by the P438 promoter [26]. To obtain this plasmid, two PCR fragments were obtained using genomic DNA as a template, and the pair of primers p_MPN360_F / P438_MPN361_R and P438_ ΔNt_L_hemK_F / p_hemk_R. To obtain the final construct, the two PCR products were cloned by Gibson assembly into the pLoxPuroCre vector opened by PCR using primers p_pRec_F and p_pRec_R. A similar strategy was used to obtain the pRec_ΔNt_M_hemK plasmid, which contains instead an N-terminally truncated version of the *mpn362* gene starting at residue M290. In this case, the second PCR product was replaced and obtained by using primers P438_ ΔNt_M_hemK_F and p_hemk_R. Similarly, the pRec_hemK_FLAG plasmid contains the essential genes *mpn360*, *mpn361* plus a C-terminally FLAG tagged version of *mpn362*. To obtain this construct we performed two consecutive PCR reactions, first with the pair of primers p_MPN360_F and hemk_FLAG_R, and then with primers p_MPN360_F and p_hemk_R. The resulting PCR product was cloned by Gibson assembly into the pLoxPuroCre vector opened by PCR using primers p_pRec_F and p_pRec_R. Finally, the pRec_pheST_Stop_P438 plasmid contains the essential genes *mpn105* (pheS) and *mpn106* (pheT), with the exception that the PheT coding sequence was premature interrupted by placing a stop codon after residue T207, followed by the P438 promoter sequence to drive the expression of a second PheT fragment starting at residue M208. To obtain this construct, two PCR fragments were obtained using genomic DNA as a template, and the pair of primers p_pheST_F / pheT_StopP438_R and StopP438_pheT_F / p_pheST_R. To obtain the final construct, the two PCR products were cloned by Gibson assembly into the pLoxPuroCre vector opened by PCR using primers p_pRec_F and p_pRec_R.

### Validation of transposon data by constructing specific mutant strains

To assess the quality of the transposon data, we constructed specific mutants to mimic the effect of a subset of transposon insertions, and thus confirm the viability of these gene disruptions. As examples, we constructed *hemK* and *pheT* mutants using the SURE-editing tool, a genome-engineering method that combines genome edition by oligo recombineering and selection of edited cells by plasmid integration mediated by site-specific recombinases [47]. A major advantage of this combined strategy is that it allows the introduction of gene platforms, thus opening the possibility to perform precise gene replacements even in essential regions of the genome. We took advantage of this improvement to introduce specific mutations affecting *hemK* or *pheT* genes without disrupting the essential genomic context. To obtain the *hemK* mutant derivatives (M129_ΔNt_L_hemK, M129_ΔNt_M_hemK and M129_ hemK_FLAG) and the *pheT* mutant (M129_pheST_Stop_P438), we co-electroporated the M129-GP35 strain [47] with 0.5 nmols of editing oligo (ssDNA_hemK for hemK mutant derivatives or ssDNA_pheST for pheT mutant) and 2µg of the corresponding selector plasmid described above (pRec_ΔNt_L_hemK, pRec_ΔNt_M_hemK, pRec_hemK_FLAG and pRec_pheST_Stop_P438). The editing oligo contains two 40-nucleotide-long regions with homology to the adjacent sequences of the intended deletion area, and separated by a lox66 recognition-site that is used to select the edited cells via Cre-integration of the selector plasmid. To allow expression of the puromycin marker and induce Cre expression from the selector plasmid, cells were recovered after electroporation at 37°C for 2h in the presence of aTc. Transformed cells were then selected during 24h in 25 ml cultures supplemented with aTc and puromycin to ensure oligo recombineering and subsequent selection of the edited cells by the integration of the selector plasmid. As described above, the selector plasmid was engineered to contain a DNA platform aimed to complement the deletion of essential regions mediated by the editing oligo, and to introduce the desired mutations. Finally, puromycin-resistant colonies were isolated from agar plates supplemented with puromycin, and DNA rearrangements verified by Sanger sequencing and PCR analysis using primers listed in Supplementary Table S6. The backbone of the selector plasmid was further excised from the genome by electroporating the mutants with 2µg of pGentaVcre suicide vector [47], which contains the vCre recombinase. PCR analysis and Sanger sequencing were performed using the primers listed in Supplementary Table S6 to confirm the intended genome edition.

### Immunoblot analysis

Mycoplasma cells were washed with 1× PBS twice, scraped off from the flasks and centrifuged at 13,100 *g* for 10 min. The cell pellet was then resuspended in 1% SDS lysis buffer, and disrupted using a Bioruptor sonication system (Diagenode) with an On/Off interval time of 30/30 s at high frequency for 10 min. Cell lysates were quantified by using the Pierce ^TM^ BCA Protein Assay Kit and mixed with NuPAGE ^TM^ LDS loading buffer (Invitrogen). Ten µg of each cell extract was subjected to electrophoresis through NuPAGE^TM^ 4–12% Bis-Tris precast polyacrylamide gels (Invitrogen), and proteins transferred onto nitrocellulose membranes using an iBlot^TM^ dry blotting system (Invitrogen). For immunodetection, membranes were blocked with 5% skim milk (Sigma) in PBS containing 0.1% Tween 20 solution and probed with the following antibodies. For the assessment of the ter625 termination activity, we used rabbit polyclonal antibodies anti-CAT (abcam, 1:2,000), anti-DsRed (Takara, 1:2,000) or anti-RL-7 (kind gift of Dr. Herrmann, Heidelberg University, 1:5,000) as loading control. For the detection of putative HemK isoforms, we used the mouse monoclonal anti-FLAG M2 (Sigma, 1:5,000). Anti-mouse IgG (1:10,000) or anti-rabbit IgG (1:5000) conjugated to horseradish peroxidase (Sigma) were used as a secondary antibody. Blot signals were detected using the Supersignal™ West Pico or Femto Chemiluminescent Substrate Detection Kit (Thermo Scientific) and the LAS-3000 Imaging System (Fujifilm).

### Genome sequencing and assembly of the *M. pneumoniae* strain used in this study

To increase the accuracy of the transposon insertion mapping, we sequenced the genome of our *M. pneumoniae* lab strain (M129_LSR). For this, genomic DNA was extracted using the MasterPure^TM^ DNA Purification Kit (Epicentre, Cat. No. MCD85201). Libraries were prepared using the NEBNext® DNA Library Prep Reagent Set for Illumina® kit (ref. E7370L) according to the manufacturer’s protocol. Briefly, 500 ng of DNA was fragmented to approximately 600 bp and subjected to end repair, addition of “A” bases to 3′ ends, ligation of NEBNext hairpin adapter and USER excision. All purification steps were performed using AgenCourt AMPure XP beads (ref. A63882, Beckman Coulter). Library size selection was done with 2% low-range agarose gels. Fragments with average insert size of 660 bp were cut from the gel, and DNA was extracted using QIAquick Gel extraction kit (ref. 50928706, Qiagen) and eluted in 15 µl EB. The adapter-ligated size-selected DNA was used for final library amplification by PCR using NEBNext® Multiplex Oligos for Illumina. Final libraries were analyzed using Agilent DNA 1000 chips to estimate the quantity and check size distribution, and were then quantified by qPCR using the KAPA Library Quantification Kit (ref. KK4835, KapaBiosystems). Libraries were loaded at a concentration of 15 pM onto a flow cell together with other samples at equal concentration (half a run), and were sequenced 2 x 300 on Illumina’s Miseq. Reads preprocessing consisted of trimming reads using SeqPurge v0.1-478-g3c8651b [66] with a minimum read length of 80 and default parameters for base calling quality threshold. Genome assembly was performed with the resulting reads using SPAdes genome assembler v3.14.1 using default parameters [67]. Quality of the assembly was assessed using QUAST v5.0.2 (Quality Assessment Tool for Genome Assemblies) [68]. Then, both single nucleotide polymorphisms and indels variants were called using snippy v4.6.0 (https://github.com/tseemann/snippy) with default parameters. Snippy uses FreeBayes v1.3.2 (arXiv:1207.3907) internally to call variants, and subsequently filters variants with low support. All variants passing the default filters (minimum coverage of 10, minimum quality of 100, minimum fraction of reads supporting the alternative allele of 0.9) were considered as high confidence. Additional manual curation was then performed by aligning the resulting genome with the *M. pneumoniae* M129 reference genome available at NCBI under accession NC_000912.1. This was performed using NucDiff v2.0.3 [69], a tool which allows comparison of closely related sequences, and rigorous analysis of local differences and structural rearrangements. Raw sequencing files and the resulting assembled genome have been deposited in ENA under the study accession number PRJEB80886 and accessible in the link http://www.ebi.ac.uk/ena/data/view/PRJEB80886.

### Comprehensive genome annotation of the *M. pneumoniae* laboratory strain

The genome annotation was updated based on the new genome sequence (816,357 bp; Supplementary Data S2; deposited under accession PRJEB80886 in ENA), followed by manual curation to define the most likely protein coding regions. For this, we used gene orthology analysis and experimental data, including transcription start sites (TSS) and peptide data collected by multiple MS experiments [1,34,50]. This genome annotation includes 707 protein-coding genes and several types of functional RNAs (3 rRNAs, 1 tmRNA, 1 RNaseP RNA, 1 4.5S RNA, 37 tRNAs) and 186 ncRNAs [15].

For the analysis of the transposon data, we considered the annotation of other genomic features, such as operons (n=323, considering 2 or more protein-coding genes and/or functional RNAs in the same transcriptional unit), TSS (n=856) and predicted promoters (n=1,430) [1,34,51,70]. Transcription terminator sites (TTS, n=434) were also defined as the position where an intrinsic terminator is predicted in the 500 bp downstream of a gene stop codon [71]. The 5’ and 3’ untranslated regions (UTR5, n=483; and UTR3, n=323) were defined from the closest annotated TSS or TTS to the gene start or end, respectively. When no TTS was found, we tracked the 150 bp down-stream of the related gene. The regions in between non-overlapping genes and expressed from the same operon were also tracked and referred to as inter-genic-intra-operon regions (iGiO; n=240). Finally, we also included predicted ribosome binding sites (RBS, n=279; included as a binary value where 1 indicates the presence of any of the possible Shine–Dalgarno sequences known to act as an RBS [50]) located no more than 15 bp upstream of a start codon, regions with experimental signal for protein occupancy and chromatin immunoprecipitation sequencing (ChIP-seq) data for different transcriptional regulators [1] (n=2,454), and a set of genomic regions lacking any gene annotation after excluding all putative ORFs (‘noann’; n=134) [51]. Finally, 155 regions were annotated as ‘noexp’ sharing a low transcriptional signal in both strands (i.e. log_2_[CPM]<3 in both genomic strands).

All the annotations described above are detailed in Supplementary Data S2, including the genomic coordinates and functional information, such as conservation studies and proteomic information [50]. Motivated by the nature of this study, we also reanalyzed the RNA-sequencing data for the new version of *M. pneumoniae* M129_LSR genome using the experiment found at ArrayExpress database under accession identifier E-MTAB-6203 (two replicates after 6 hours of growth at 37°C) to associate expression values to each genomic region considered in this study. Finally, for each of these annotations, percentage repeated is calculated by taking as reference a set of 85,077 bp genome positions labeled as ‘repeated’. This is done by running a sliding window of 21 bp, extracting the sequence and looking for perfect matches in the direct and reverse-complementary genome sequence of *M. pneumoniae.* If the subsequence appears at least twice, the position of the window will be labeled as repeated and assumed they will present low and/or limited mapping quality [29] (Supplementary Data S1 and Supplementary Data S2).

### Transposon mutant libraries preparation

Transposon mutant libraries were prepared as previously described [29]. Briefly, *M. pneumoniae* wild-type cells were transformed by electroporation [72] with 2 μg of mini-transposon plasmid DNA pMTnCat_BDPr or pMTnCat_BDter. Mutant libraries (referred herein “P” and “T”) were selected in 5 ml cultures supplemented with chloramphenicol, and the resulting transformants scrapped off the flasks in 1 ml of fresh medium to obtain a cell stock referred to as passage 0 (P0 and T0, respectively). Transposon mutants were then serially cultured through ten consecutive passages to eliminate death cells and further assess the impact on growth fitness of the transposon insertions represented in the libraries. For this, we inoculated 25 μl of P0 or T0 in 5 ml of Hayflick medium supplemented with chloramphenicol. After 4 days of culture (approximately 10 cell divisions), cells were scraped off the flasks in the same culture medium to also recover non-adherent cells. One milliliter of this cell suspension (referred to as passage 1: P1 and T1) was used for genomic DNA isolation using the MasterPure^TM^ DNA Purification Kit (Epicentre, Cat. No. MCD85201), and 25 μl to inoculate the next passage. This procedure was repeated until passage 10 (P10 and T10), including sample collection for all the intermediate passages. To account for sampling batch effects, cell passaging and sample collection were performed in duplicate.

### Transposon insertion sequencing and insertion calling

Illumina transposon sequencing libraries were constructed from genomic DNA of “P” and “T” mutant pools extracted at different passages, using the NEBNext Ultra DNA Library Prep kit (#E7370L) according to previously published protocols [29]. Transposon libraries were sequenced on a HiSeq 2500 platform using HiSeq v4 sequencing chemistry and 2 × 125 bp paired-end reads. The raw data was submitted to the ArrayExpress database (http://www.ebi.ac.uk/arrayexpress) with the accession identifiers E-MTAB-8918 (P library) and E-MTAB-14533 (T library).

The software FASTQINS was used to define the genome positions and the read count coverage of each transposition event from raw sequencing files by looking for reads containing the IR sequence *TACGGACTTTATC* [29]. The insertion points are then determined by mapping to *M. pneumoniae* M129_LSR allowing 1 mismatch in the genomic sequencing. The final output consists of each genome base found inserted and the read-count measuring how many times each transposon insertion is mapped (Supplementary Data S1). General mapping was assessed by calculating each sample genome coverage as the total number of insertions found in each sample normalized by the genome length of *M. pneumoniae* M129_LSR (816,357 bp). Also, a set of known NE genes (n=29; 21,599 bp in total) was used to provide an average of the maximum saturation as the total number of insertions found mapping in these genes and divided by the region covered by these same genes (Supplementary Data S3) [23].

### Metric and essentiality estimation by a GMM model

A reference of gene essentiality was estimated using both P1 and T1 samples and merging the 2 replicates of each library (PT1). To normalize for differences in transposon saturation and the coverage of the sequencing technique, the top and bottom 5% of insertions in terms of read-count in each sample are not considered in the merging process. Then, gene linear densities (LD) are calculated as the number of insertions mapped to a given genomic annotation normalized by its base pair length. Repeated regions where the ambiguity prevents efficient mapping of insertion were ignored as done in previous studies [29]. The list of positions is referenced in Supplementary Data S1.

Essentiality assignment in essential (E), fitness (F1 and F2), and non-essential (NE) was performed by using the Gaussian Mixture Model from the ANUBIS package [29]. This approach applies a probabilistic estimation of groups in the data, assuming they are generated from a mixture of *k* number(s) of Gaussian distributions. To define *k* we took into consideration the evolution of the Akaike Information Criterion (AIC) and Bayesian Information Criterion (BIC) to find the best compromise between goodness of fit and number of parameters (Fig. S3). Also, this approach was shown to outperform other estimation methods and present the advantage of not requiring a reference ‘gold’ set of genes with known essentiality categories [29]. The relation between genomic annotations found in *M. pneumoniae* M129_LSR and their estimated essentiality category by analysis condition can be found at Supplementary Data S4.

For structural and regulatory regions, we performed the same GMM analysis using the PT1 combined sample with 2 and 4 components. As elements categorized as F1 always correspond to E in the 2-components model and the same occurs for F2 which were associated with NE, we decided to simplify the prediction to consider only E and NE classes. This is in fact expected due to the way the GMM algorithm minimizes differences between components (i.e. the most explanatory threshold with two components separates E from NE, with four E+F1 from F2+NE elements). For the manual curation and plotting in the analyses related to LD, we filtered out annotations shorter than 5 bp or having less than 25% of its sequence repeated. The complete list of selected elements, associated information and whether they were selected or not for downstream analyses is included in Supplementary Data S5.

### Essentiality decay exploration and clustering

Decays in LD associated with each genomic annotation were explored by a k-means clustering algorithm to group annotations based on their response to passaging [73]. As the number of decay groups is initially ignored, we define the minimum number of plausible clusters by inspecting the distortion associated while increasing the number of clusters. This distortion value is calculated for each given required number of clusters and it corresponds to the sum of squared distances between observations and their dominating associated centroids. As similarly done for assigning essentiality categories with a GMM, we selected four as the number of clusters to predict as this is the value that minimizes the distortion (*i.e.*, sum of squared errors) and thus expected to provide a balanced compromise between goodness of fit, observed decay shapes, and number of parameters without overfitting the prediction (Fig. S5A). For each explored annotation (*e.g.*, genes), we calculated a metric (*e.g.*, LD) and its value along subsequent passages to represent the persistence of insertions in a specific annotation, or fitness impact on cell growth (Fig. S5). This value tends to decay due to both cell death and sampling effects. Thus, we normalized values by the average found for NE positions to avoid the biases derived by the original insertion coverage rate at the sample-level. Consequently, values can range from 0 to values above 1 (*e.g.*, when an annotation is more enriched in insertions than NE regions in that same sample). In addition, the definite integral of normalized linear densities along passages 1 to 4, 6, 8 and 10, corresponding to the area covered by the decay curve (AUC), are approximated by the trapezoidal rule using the function ‘trapz’ from numpy [74]. This value represents an integrated metric to represent the observed decays in a quantitative manner and it is also representative of the cluster to which a gene will belong (Fig. S5B). We also looked for the different correlation in AUC values between all passage combinations from sample pairs to ensure this metric is robust and independent of the passage conditions. In fact, Pearson’s standard correlation coefficient between subsets was always greater than 0.85, suggesting that the same analysis could be efficiently performed with a smaller number of samples (Fig. S6).

To perform statistical assessments in the calculation of AUC differences between P and T libraries, we calculated the AUC for density/reads normalized by NE regions separating each replicate (1 and 2) and library type (P and T). This generates a couple of measures for each condition that are then compared by one-tailed T-test and assuming the differences between P and T are significant if *P* < 0.05, while a fold-change indicates the direction of the change (Supplementary Table S5). It is important to remark that ignoring repeated positions would inflate the apparent LDs for genes with higher percentage of repetition as very few numbers of positions, generally inserted, will be taken into account. Thus, the k-means cluster prediction algorithm overestimates the evolution of the trajectories of these genes affecting the definition of coherent clusters. To alleviate this, we discard these 15 genes during the clustering step: *mpn091*, *mpn128*, *mpn130*, *mpn203*, *mpn204*, *mpn367*, *mpn371*, *mpn410*, *mpn413*, *mpn463*, *mpn467*, *mpn468a*, *mpn486*, *mpn501*, *mpn502*.

### Local essentiality estimation by a sliding window approach

Essentiality was assessed in an annotation-independent manner by means of a sliding window to capture potential functional units and explore small genomic loci [13]. To this end, we defined a minimum sliding window length by comparing windows of different sizes (from 3 to 150) sampled from E and NE windows from genes with known essentiality, and using the Gaussian Mixture Model to differentiate between E and NE populations of windows in an unsupervised manner. Using windows sampled from the reference gold set of genes, we observed that the probability to wrongly assign the NE category to a window from an E gene (*i.e.*, false discovery rate) is less than 0.1%, when using a size of 31 bp. Then, we extracted different metrics associated with each 31 bp region in the *M. pneumoniae* M129_LSR genome and defined NE domains by considering contiguous windows together when they were classified as NE. Unassigned nucleobases were labeled as E (Supplementary Data S6). To make the analysis stricter, we only considered DNA domains covering ≥5 bp and with less than a 10% of the segment repeated.

To identify possible NE regions within E genes, we mapped predicted domains against different annotations to obtain the overlap (general) and the percentage they represent relative to gene annotations (Supplementary Data S7). DNA domains with different essentiality profiles were also predicted by separating P and T libraries, in addition to computing AUC values (passages 1 to 8) for the predicted domains. For measuring the extensions from an evolutionary perspective, we ran BlastP against 109 bacterial reference genomes [50] and counted the number of additional or reduced amino acids in each gene based on the alignment positions (Supplementary Data S7).

To assess possible differences in transposon coverage across DNA domains known to bind to transcriptional repressors or other DNA-binding proteins (POD data previously published) [1], we calculated different metrics (number of insertions, LD, total read count and average number of reads per insertion) for the length of domain under inspection and compared to the same number of bp before and after the domain (Supplementary Data S8 and S9).

### Transposon-insertion enrichment analysis

To identify insertions presenting a selected enrichment after 10 passages, we only considered insertions that were conserved in passage 1 and 10 in both replicates for each P and T condition. Then, we took the read count observed in each condition and replica and applied a TMM (Trimmed Mean of M-values) normalization to be able to group by passage and compare them. The TMM method is commonly used to normalize sequencing read counts in RNA-Seq experiments and to ensure that the overall distribution of read counts is comparable across samples by removing biases that may arise due to differences in library sizes and gene-specific effects. It is important to note that TMM normalization assumes that the majority of insertions are not differentially selected across samples. With the normalized read counts, we proceed by calculating the fold-change observed between last and first passage and assessing the statistical differences by the p-value given by a one-tail T-test and corrected by the Bonferroni correction method (Supplementary Data S10). Then, these insertion positions were analyzed in relation to the annotations in Supplementary Data S2 and assigned in one or more of the following categories: overlap E/F1 gene, overlap F2/NE gene, upstream E/F1 gene, upstream F2/NE gene, overlap UTR5, upstream UTR5, overlap with iGiO, overlap with no transcribed region, between two gene ends, and not classified (not present in any of the previous; Supplementary Data S10).

Finally, for insertions located upstream to a gene, we also added the information of expression recalculated for the new *M. pneumoniae* M129_LSR genome with the E-MTAB-6203 dataset and COG categories (Supplementary Data S11). For statistical tests in the COG enrichment, after grouping genes by COG category, we counted the number of genes that are affected and not affected by an enriched insertion located up to 250 bp upstream to their start codon. Then, we calculated the enrichment p-value for each COG category using a hypergeometric test and adjusted it using the Benjamini-Hochberg method (Supplementary Data S12).

## DATA AND CODE AVAILABILITY

Raw sequencing files corresponding to the promoter library (P) library can be accessioned in the ArrayExpress database at EMBL-EBI, under accession number E-MTAB-8918, and are accessible from the following link: http://www.ebi.ac.uk/arrayexpress/experiments/E-MTAB-8918. Raw sequencing files corresponding to the terminator library (T) have been deposited in the same database under accession number E-MTAB-14533 and are accessible from the following link: http://www.ebi.ac.uk/arrayexpress/experiments/E-MTAB-14533.

The genome assembly of *M. pneumoniae* M129_LSR strain can be found at the European Nucleotide Archive (ENA) under the study accession number PRJEB80886 (http://www.ebi.ac.uk/ena/data/view/PRJEB80886).

The bioinformatic functions required to reproduce the presented analyses have been implemented in our previous analysis framework that can be accessed by https://github.com/CRG-CNAG/anubis.

## ACKNOWLEDGMENTS

We acknowledge the support provided by the Genomics Unit at the Centre for Genomic Regulation sequencing the samples. This project has received funding from the European Research Council (ERC) under the European Union’s Horizon 2020 research and innovation programme MYCOCHASSIS (670216) and ERC LUNG-BIOREPAIR (101020135) ERC Advanced Grants (AdG). We also acknowledge support of the Spanish Ministry of Science and Innovation through the Centro de Excelencia Severo Ochoa (CEX2020-001049-S, MCIN/AEI /10.13039/501100011033), the Generalitat de Catalunya through the CERCA programme and to the EMBL partnership. We are grateful to the CRG Core Technologies Programme for their support and assistance in this work.

## AUTHOR CONTRIBUTIONS

S.M.V. and R.B. designed this research. S.M.V. conceptualized and performed the computational and statistical analyses. R. B. designed the transposon library, prepared the samples for sequencing with support from E.G.R, and designed and performed experimental validations. M.W. assembled the new genome version of *M. pneumoniae* and S.M.V. and R.B. curated the annotations. S.M.V. and R.B. conceptualized and prepared figures and supplementary material. L.S. provided direct supervision and resources. S.M.V., R.B. and L.S. interpreted results and wrote the manuscript. All authors read and approved the final manuscript.

## Supplementary Material

Supplementary figures

**Figure S1.**
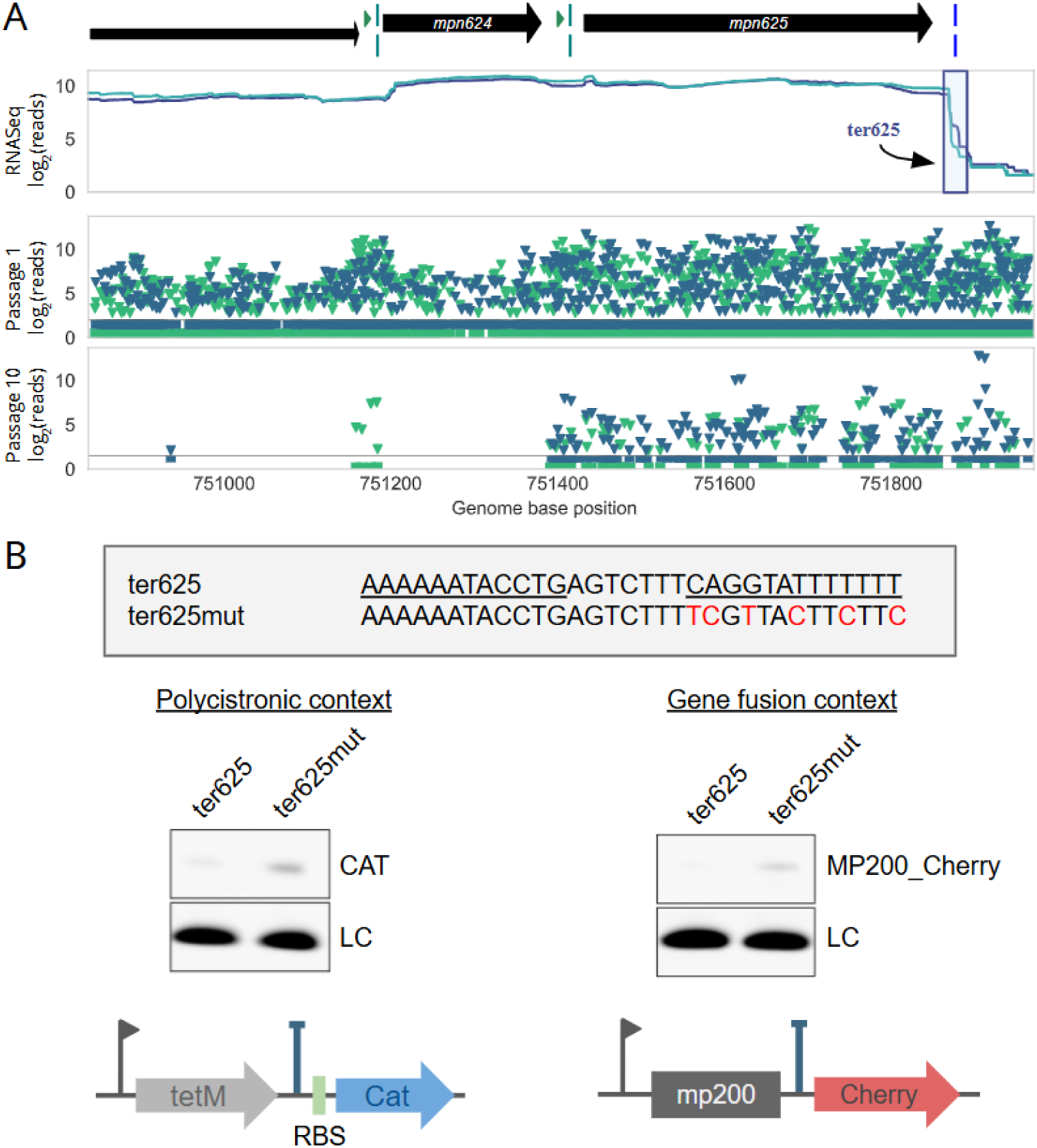
Genomic context and activity of the endogenous ter625 terminator. **A)** Scheme showing the position of the ter625 terminator sequence (downstream blue box, blue vertical line, representing the start of the hairpin) in its natural genomic context. Green arrowheads and vertical lines indicate the position of predicted promoters and TSS, respectively. Below is shown the RNAseq expression profile (n=2), and transposon insertion mapping at passage 1 and 10 along the genomic region. Green and blue triangles represent transposon insertions containing promoters or terminators, respectively. Note the enrichment of transposons containing terminators after 10 passages next to the position of the endogenous ter625 terminator. **B)** Termination activity of the ter625 sequence in two different genetic contexts using gene reporters expressed as polycistronic or fusion transcripts. On top of the panel is shown the wild-type ter625 sequence compared to a mutated version affecting the terminator hairpin structure (underlined). The expression of *cat* or cherry gene reporters was assessed by Western blot analysis in the two genetic configurations shown below the panel. The arrow indicates the promoter of the transcriptional unit, while the position of the ter625 sequence is shown as a T, just before a ribosome binding site (RBS, left panel) or after the mp200 polypeptide fusion (right panel).

**Figure S2.**
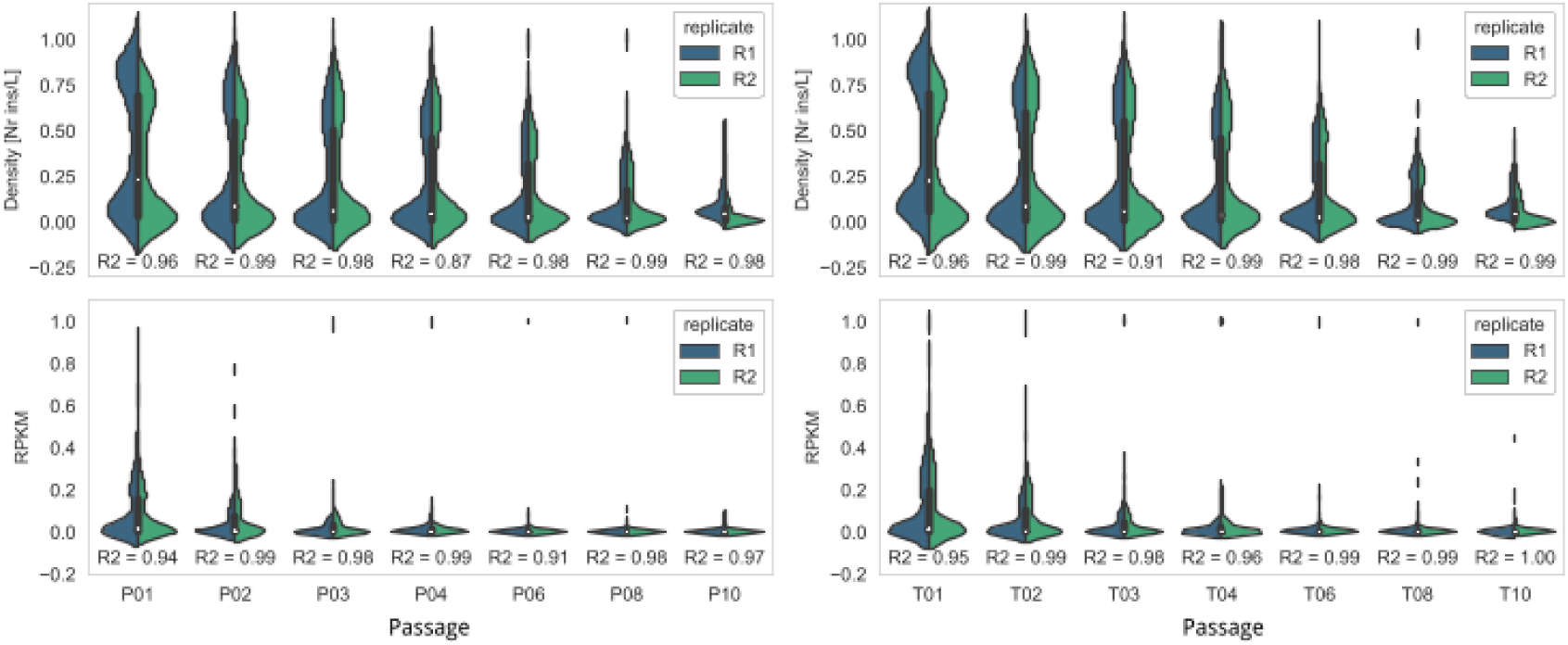
Comparative violin plot between replicates of the P (left) and the T libraries (right) at gene level. Top row represents the linear densities (total insertions normalized by gene length) across passages, while RPKM is represented at the bottom row. X-axes represent the increasing number of passages for each library condition.

**Figure S3.**
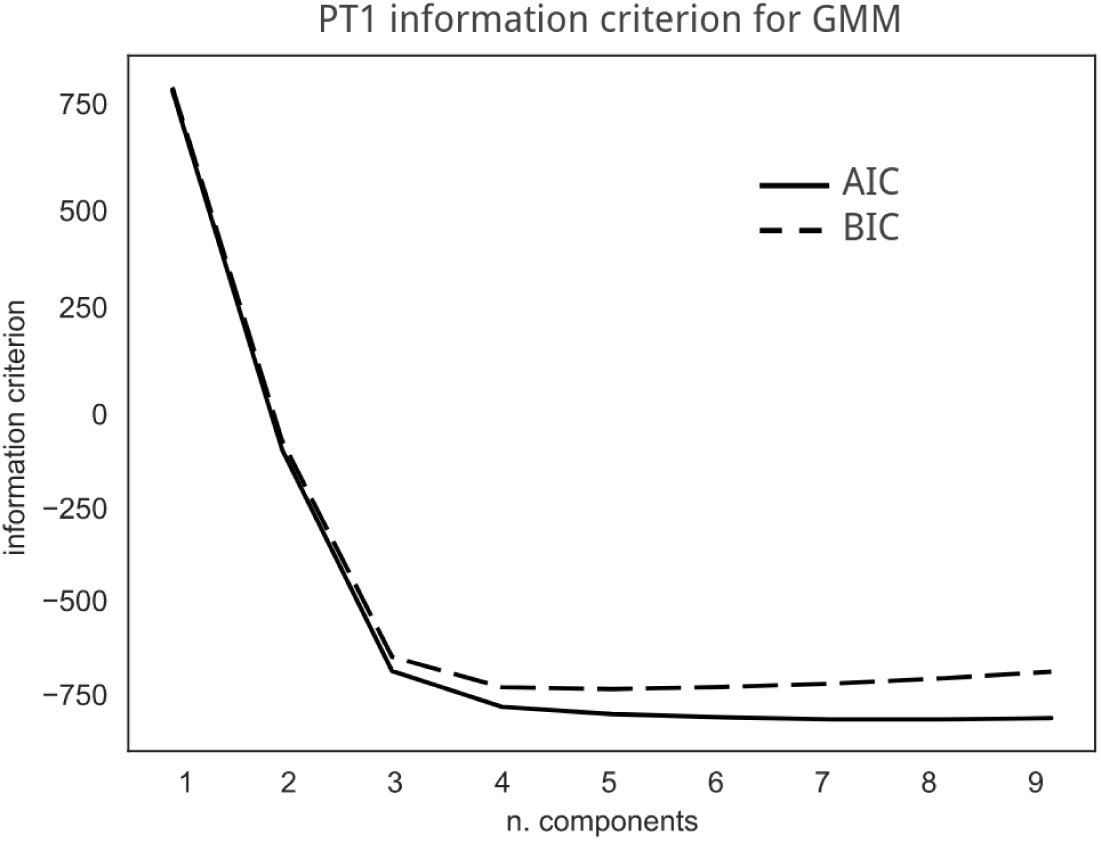
AIC and BIC for GMM of 4 components in PT1. For each number of components tested (X-axis) the information criterion (IC; Y-axis) as Akaike Information criterion (AIC) and Bayesian Information Criterion (BIC) is measured. Ideally, a good classification needs as many components as required to minimize these values and reach the plateau to avoid overfitting.

**Figure S4.**
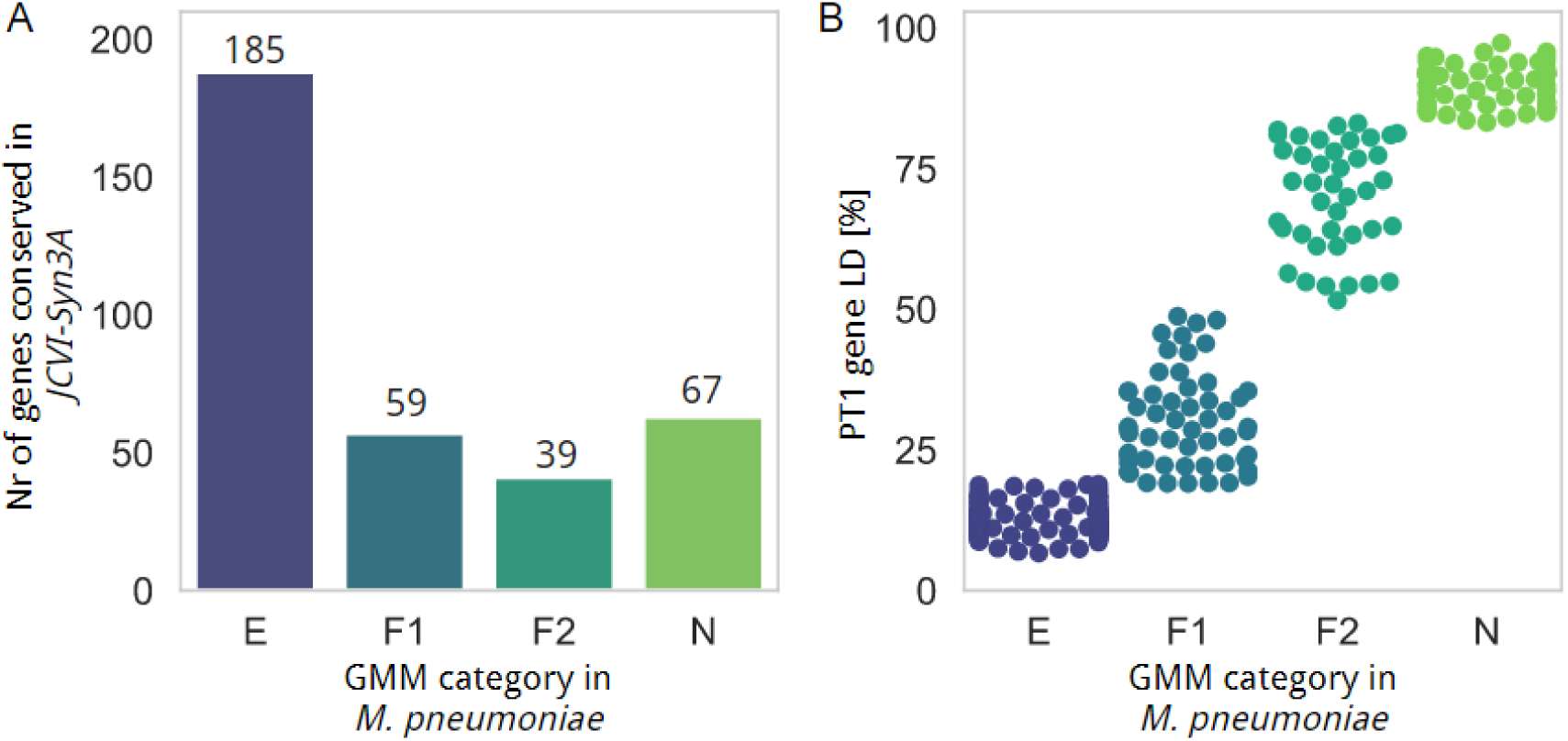
Essentiality analysis of *M. pneumoniae* protein coding genes conserved in the Synthetic *M. mycoides* JCVI-Syn3A strain. **A)** Number of genes conserved in *M. pneumoniae and M. mycoides* JCVI-Syn3A (Y-axis) by BlastP homology, and represented by their essentiality category in *M. pneumoniae* at PT1 condition (X-axis). **B)** Swarm plot where each dot represents the linear density at PT1 (Y-axis) of a conserved gene colored by essentiality categories (X-axis).

**Figure S5.**
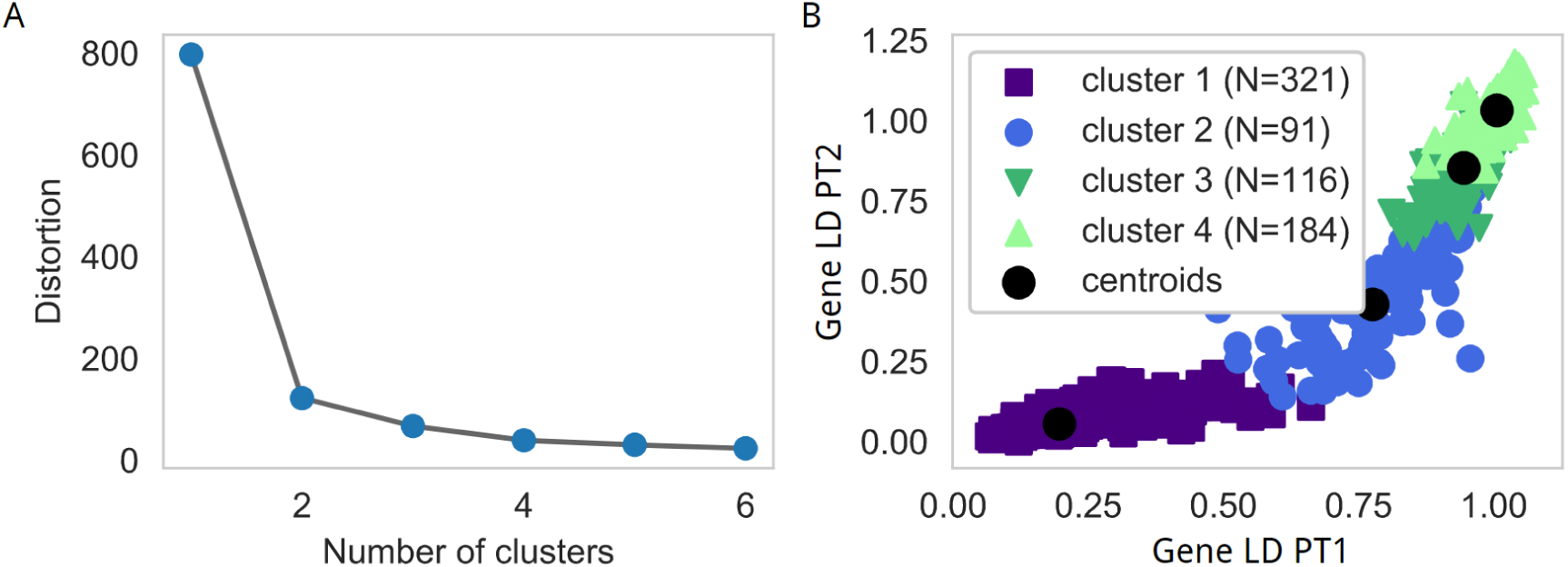
K-means decay clustering for protein coding genes using dataset PT1 to PT8. **A)** Change in the distortion (Y-axis; it measures the difference between elements within proposed clusters) by testing the model from 1 to 6 clusters (X-axis). **B)** Scatter plot relating the LDs measured at passage 1 (X-axis) and passage 2 (Y-axis) colored by the cluster assigned by k-means, from cluster 1 (purple; genes that do not maintain insertions) to 4 (light green; genes that maintain insertions). Legend shows the cluster name, color assigned, and total number of genes grouped in each cluster. Note that clusters 1 and 2 are mainly different due to the persistence of insertions after the first passage.

**Figure S6.**
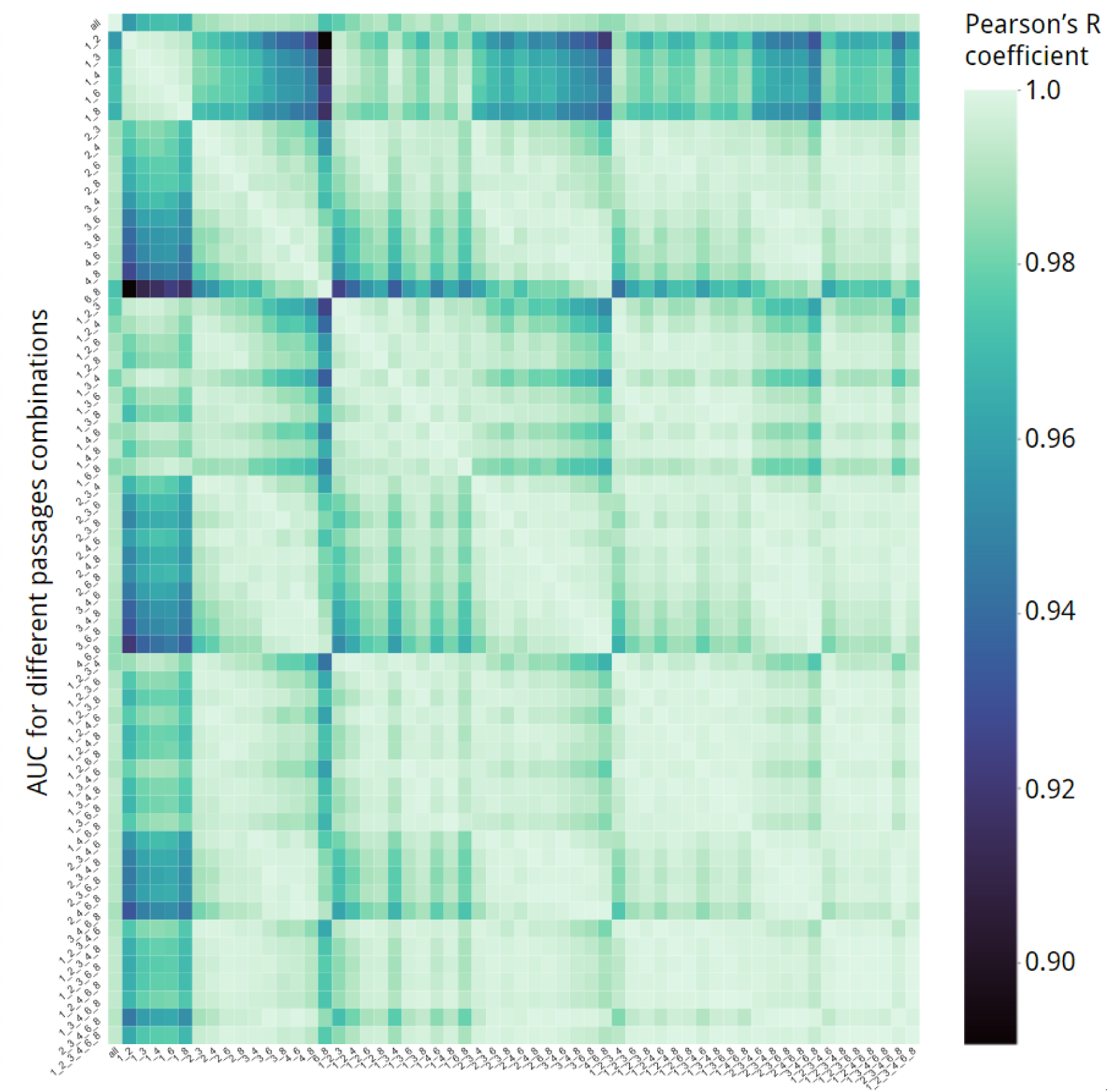
Heatmap representing Pearson’s standard correlation coefficients between AUC calculations made by subsetting different combinations of PT passage profiles, from pairs (*e.g.*, 1_2 means AUC is calculated only between passages 1 and 2) to all (‘all’). Color is lighter for higher correlation values which are always above 0.85 for every comparison.

**Figure S7.**
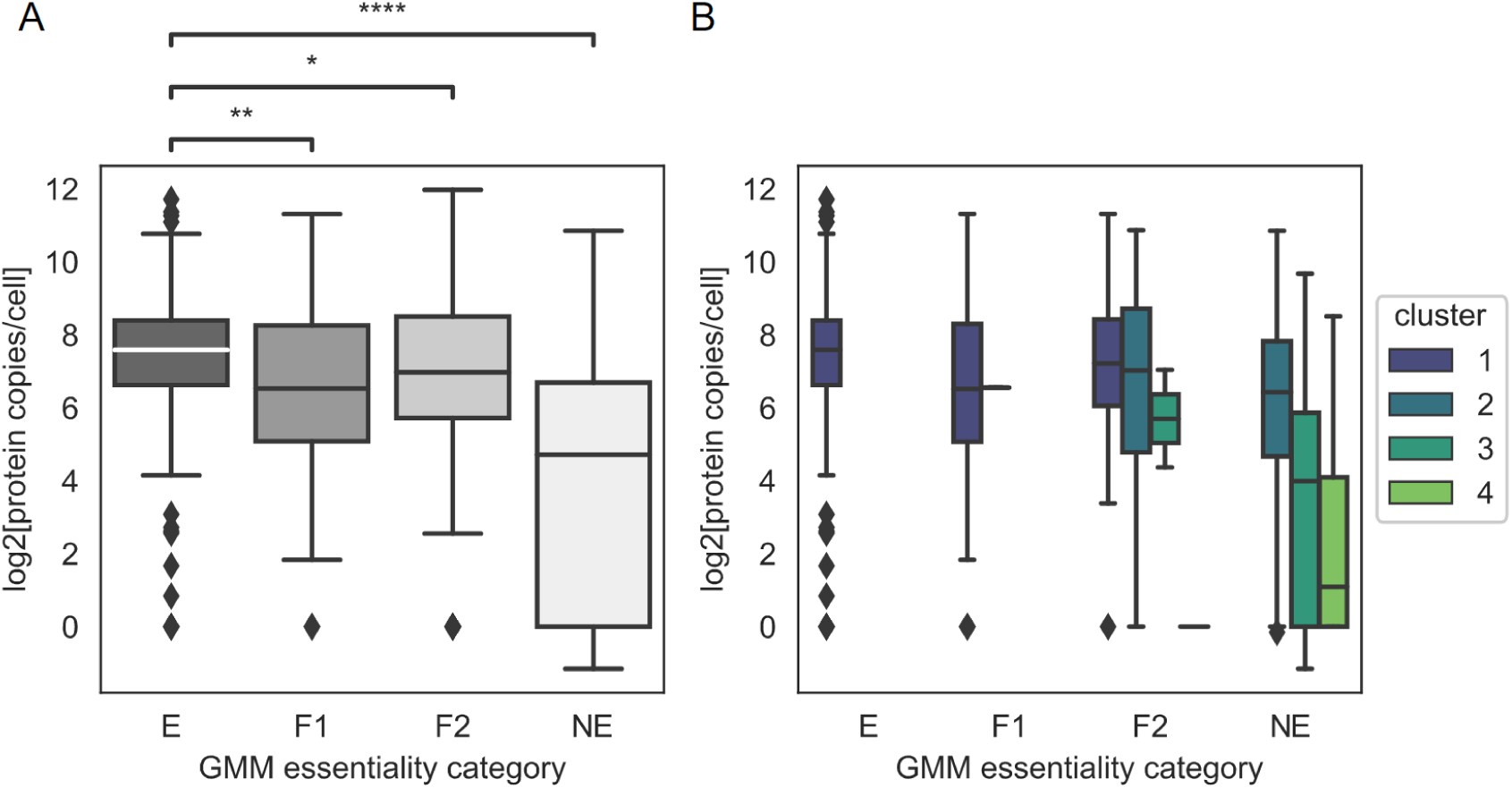
Box plots showing the relation between GMM class and k-means clusters in relation to protein copy numbers. **A)** For each predicted essentiality category at PT1 (X-axis) we relate the log_2_ transform copies per cell of the proteins in each group (Y-axis). E genes present significantly higher copies/cell levels than F1, F2 and NE genes (‘*’ Mann-Whitney *P* < 0.05; ‘****’ Mann-Whitney *P* < 0.0001). NE genes present the lowest copies per cell. **B)** Same representation as panel A, but separating genes by the cluster assigned by the k-means applied on the LD decays between PT1 and PT8. It can be observed that F2 and NE genes assigned to higher clusters (*i.e.,* more stable insertions) present lower copies per cell.

**Figure S8.**
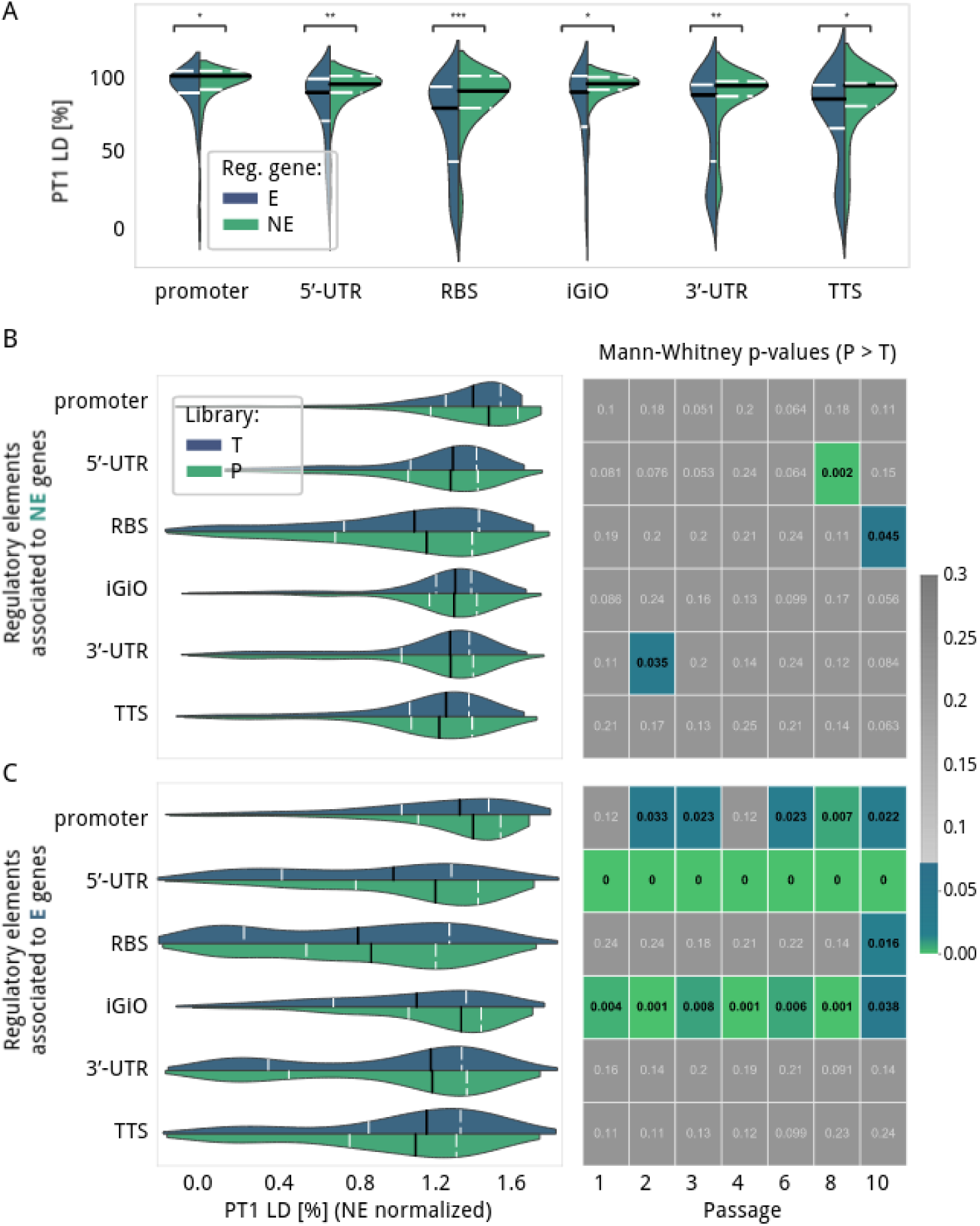
Exploration of regulatory regions by linear density. **A)** Comparative violin plot of LD measured in PT1 library for different regulatory elements (X-axis) associated with E/F1 or F2/NE regulated genes (blue and green, respectively). **B)** and **C)** Linear density comparison between P and T libraries for regulatory elements associated with NE (panel B) or E (panel C) genes. Left violin plots show the distribution of LD (X-axis) at P1 (green) and T1 (blue). Within the violins, white dashed lines represent the 1^st^ and 3^rd^ quartiles while the median is represented in black. Values are normalized by the LD found in NE regions. On the right, heatmaps with the *P* values from one-tailed Mann-Whitney tests are provided for each passage (X-axis) comparing P and T libraries. If significant by Mann-Whitney one-tailed test, the cell is colored while they remain gray otherwise.

**Figure S9.**
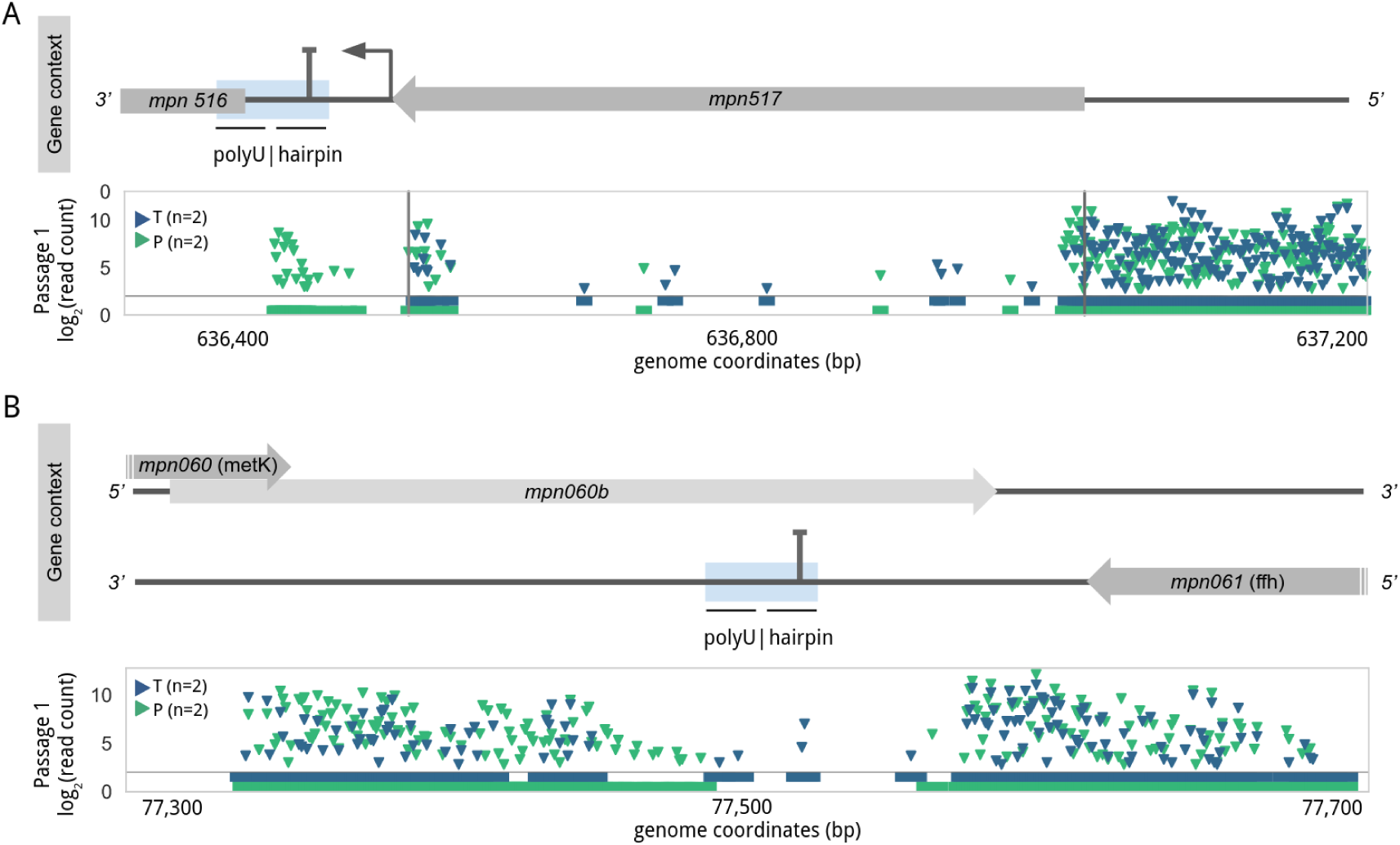
Examples of essential 3’UTR and terminator signals of *mpn517* (A) and *mpn061* (B) essential genes. For each panel, the insertion profile of the region of interest is shown. Transposons containing promoters or terminators are depicted in blue and green triangles, respectively. The genetic context is also shown on top of each panel, including orientation of the genetic elements. Predicted intrinsic terminator sequences are labeled in blue boxes showing hairpin and poly-U sequences. The 3’UTR are defined from stop codons to the terminators. It should be noticed that the 3’UTR region of *mpn517* partially overlaps with the 5’UTR of *mpn516*, which is essential in the T library but not in the P library, thus making it difficult to assess whether it is essential the terminator or the 5’UTR.

**Figure S10.**
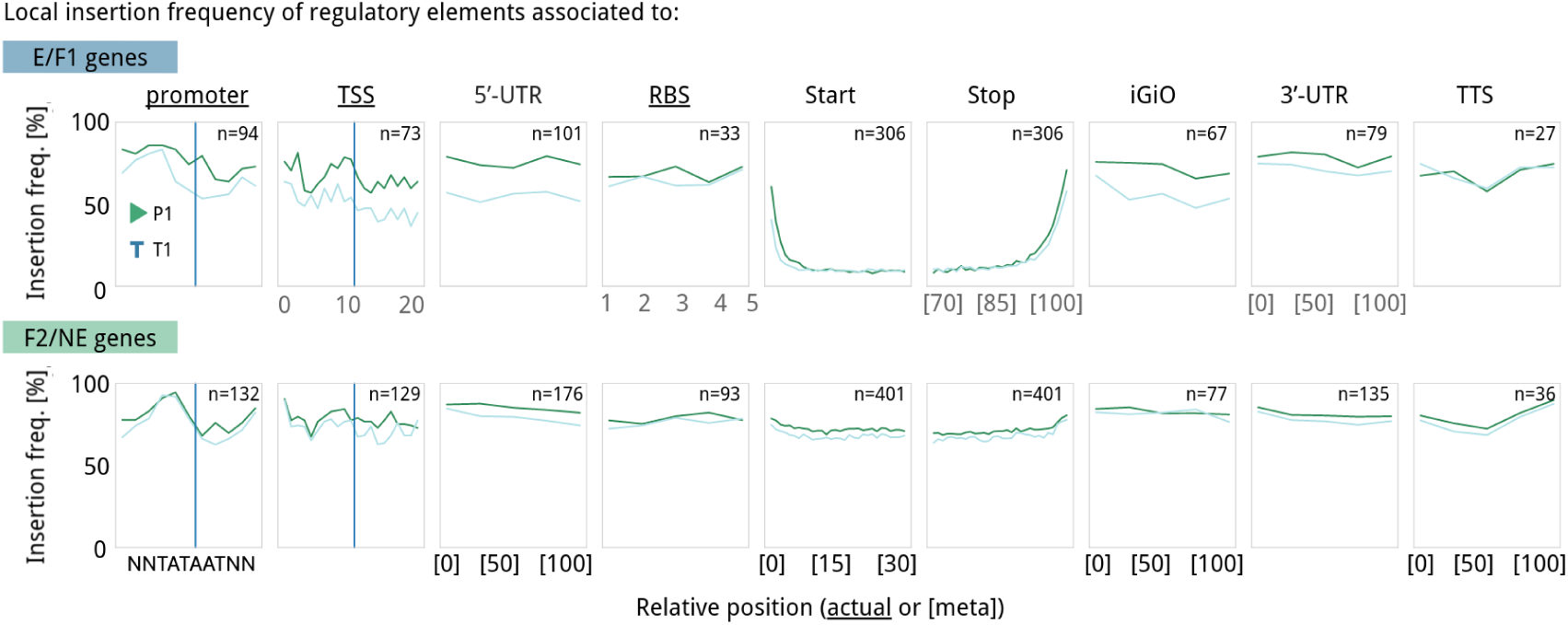
Local transposon insertion frequency in different regulatory elements. The local frequency of transposon insertions containing promoters (green) or terminators (blue) at passage 1 is shown for different regulatory elements, (including start and stop codons) associated with E/F1 (top panel) or F2/NE genes (bottom panel). Regulatory elements with a fixed sequence length are underlined and have a solid blue line representing the center of the element. Regulatory elements with a variable length, the X-axis represents the relative percentage of the covered region (minimum 5 bins for regulatory elements, 100 bins for genes and then we represent the 30 first and 30 last to get the start and stop regions). While no difference is observed for F2/NE genes, E and F1 genes are characterized for an enrichment of transposons containing promoters in upstream regulatory elements except for RBS. Downstream regulatory elements (3’-UTR and TTS) do not show global significant differences regardless of the essentiality of the associated gene. Note that N- and C-terminal regions of E genes differ in the length of transposon disruption tolerance, with C-terminal regions showing a higher tolerance for extended regions.

**Figure S11.**
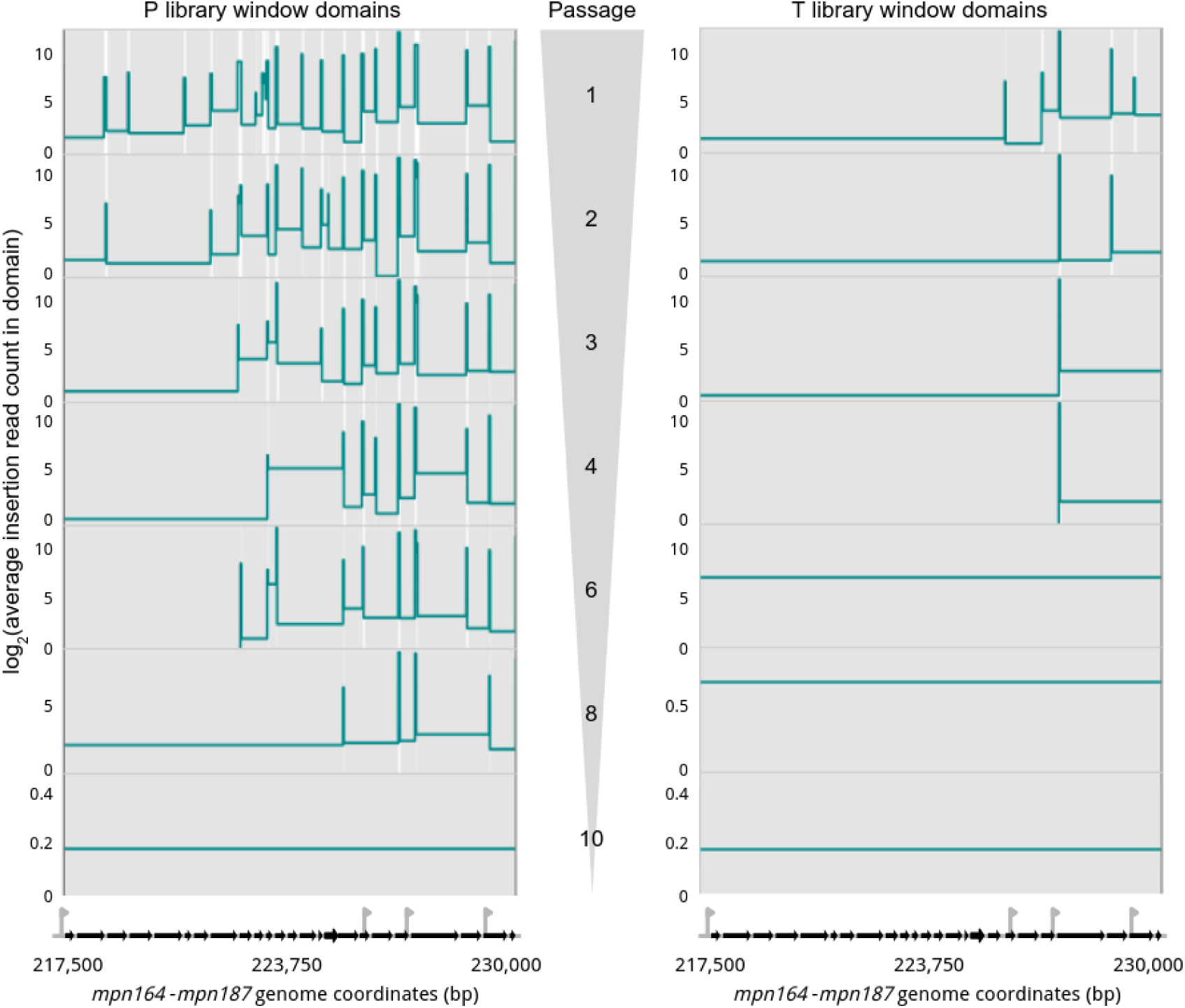
Transposon insertion windows domains from passage 1 to 10 across the operon containing genes *mpn164* to *mpn187* mainly encoding ribosomal proteins. This corresponds to the same representation as panel C in Fig. 2, but showing the window essentiality domains across increasing passages (top to bottom), highlighting the differential persistence of each transposon library depending on their proximity to the operon TSS. Gray and white areas represent E and NE regions, respectively. The height of each domain corresponds to the log2 transform of the average of transposon read count per insertion.

**Figure S12.**
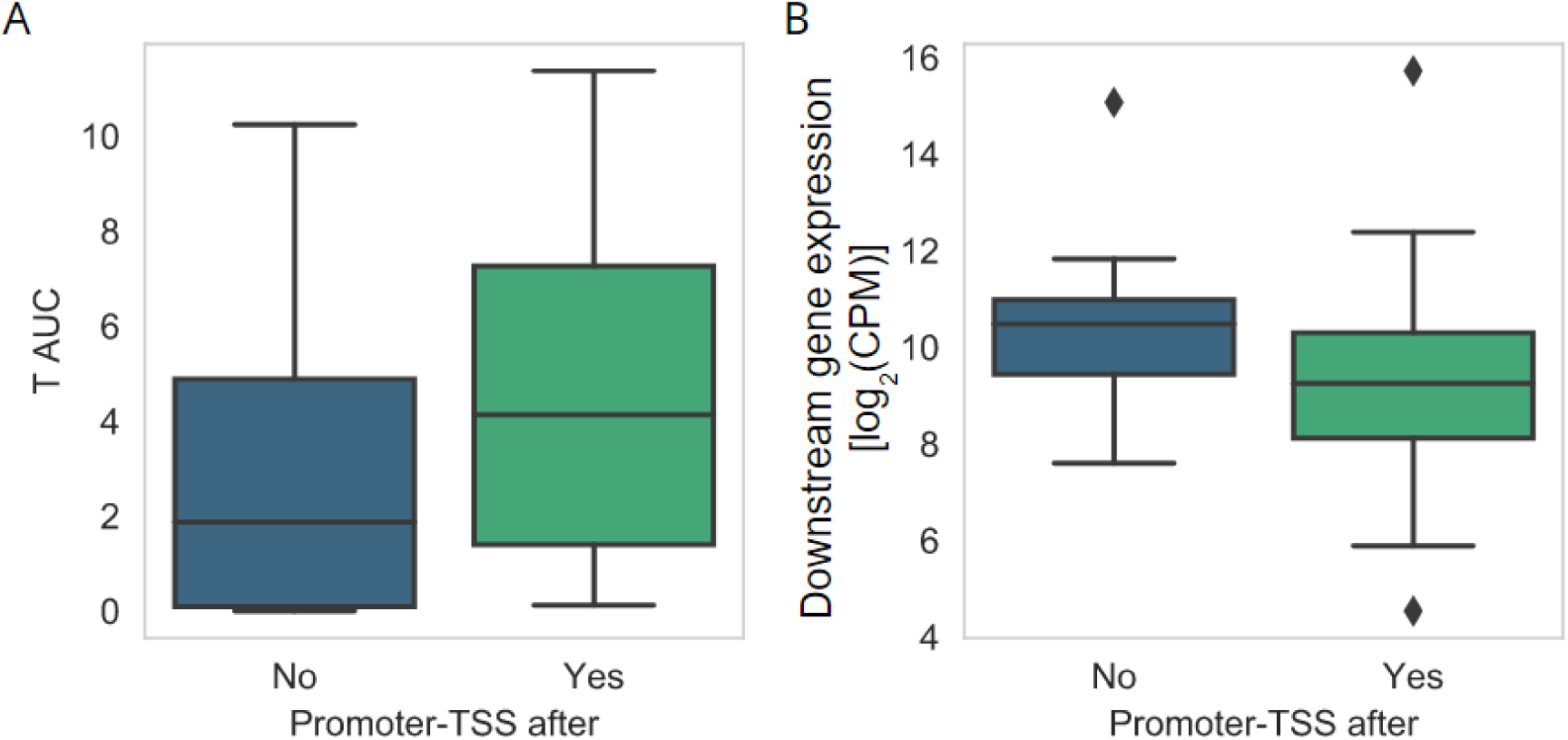
Tolerance of transcription termination depending on the transcriptional context. **A)** Box plot comparing the persistence (AUC in Y-axis) of transposons containing terminators in iGiO elements preceding E genes and separated in the X-axis by the presence (green box) or not (blue box) of a promoter and TSS that could rescue transcription. **B)** Box plot comparing the RNA expression levels (Y-axis, as log_2_-transformed read counts normalized per million reads) of E genes downstream to the same iGiO elements considered in panel A and separated by the presence/absence of rescuing promoter and TSS.

**Figure S13.**
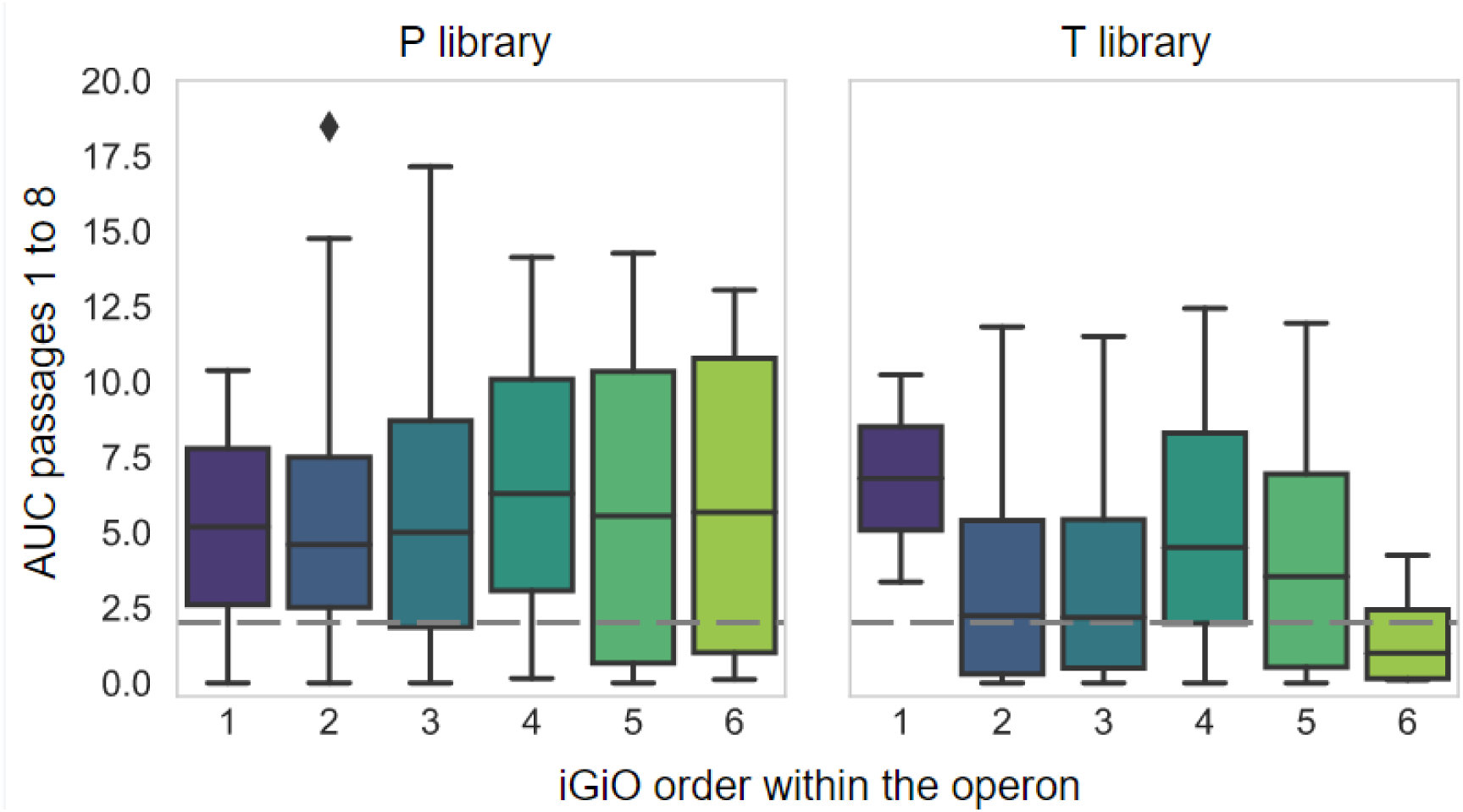
Transposon insertion persistence across passages (1 to 8) for P and T libraries considering iGiO elements preceding E/F1 genes. Box plots show the persistence (AUC in Y-axis) of transposons containing promoters (left) or terminators (right) in iGiOs elements depending on their relative position (X-axis) in the transcriptional unit (*e.g.*, 3 means it is the third iGiO within a transcriptional unit).

**Figure S14.**
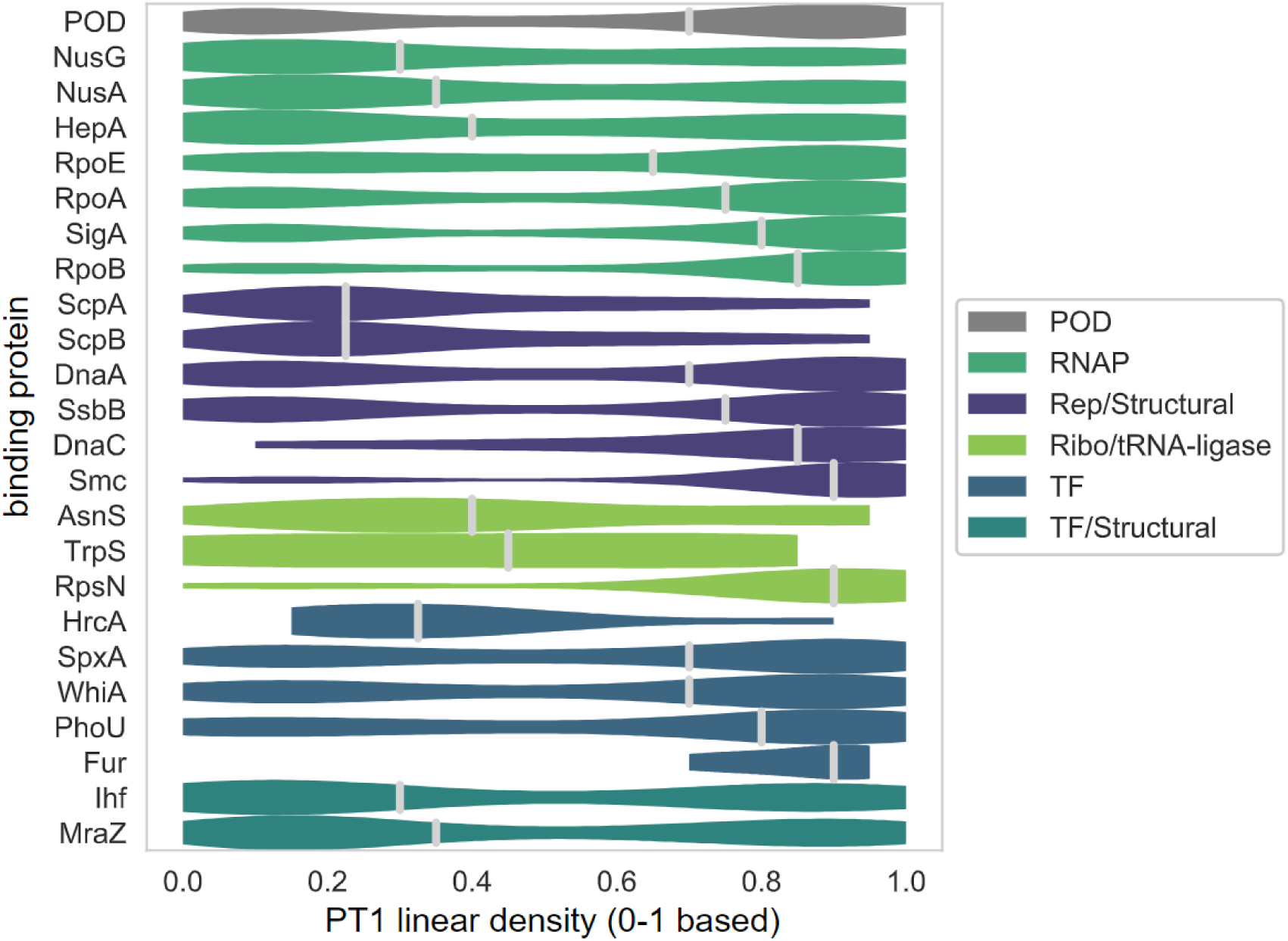
Linear densities distribution at passage 1 in library PT for DNA protection (POD) data and ChIP-seq of specific DNA-binding proteins. Horizontal violin plot shows the transposon linear density distribution for different binding and structural motifs (Y-axis), colored by the type (see legend).

**Figure S15.**
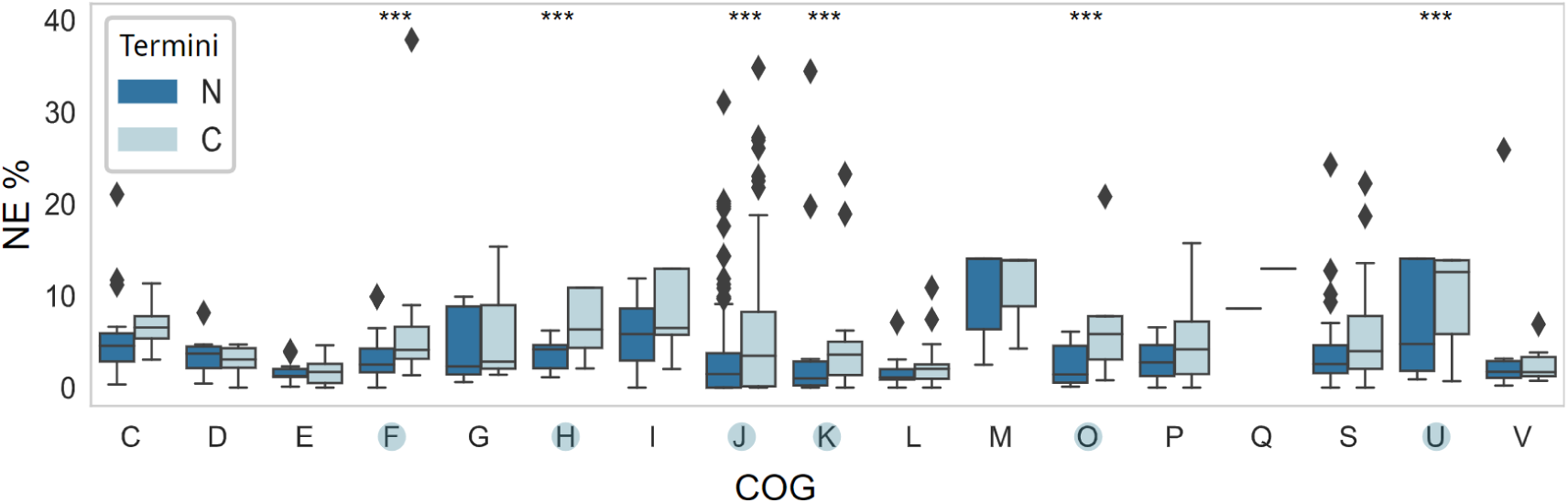
COG category analysis of N- and C-terminal extensions for E and F1 genes. Box plots represent the percentages of NE segments at PT1 condition within the N- or C-terminal extensions of E and F1 genes, which are classified by COG category (X-axis). Notice that some COG categories are missing as no E/F1 are included in those groups. COG categories presenting significant differences (Wilcoxon Test, P-value < 0.05) between both termini are marked with the asterisks and in a blue circle in the X-axis labels. These are: F - Nucleotide metabolism and transport; H - Coenzyme metabolism; J - Translation; K - Transcription; O - Post-translational modification, protein turnover, chaperone functions; and U - Intracellular trafficking and secretion. Note ribosomal genes (J) present the largest values for both ends.

**Figure S16.**
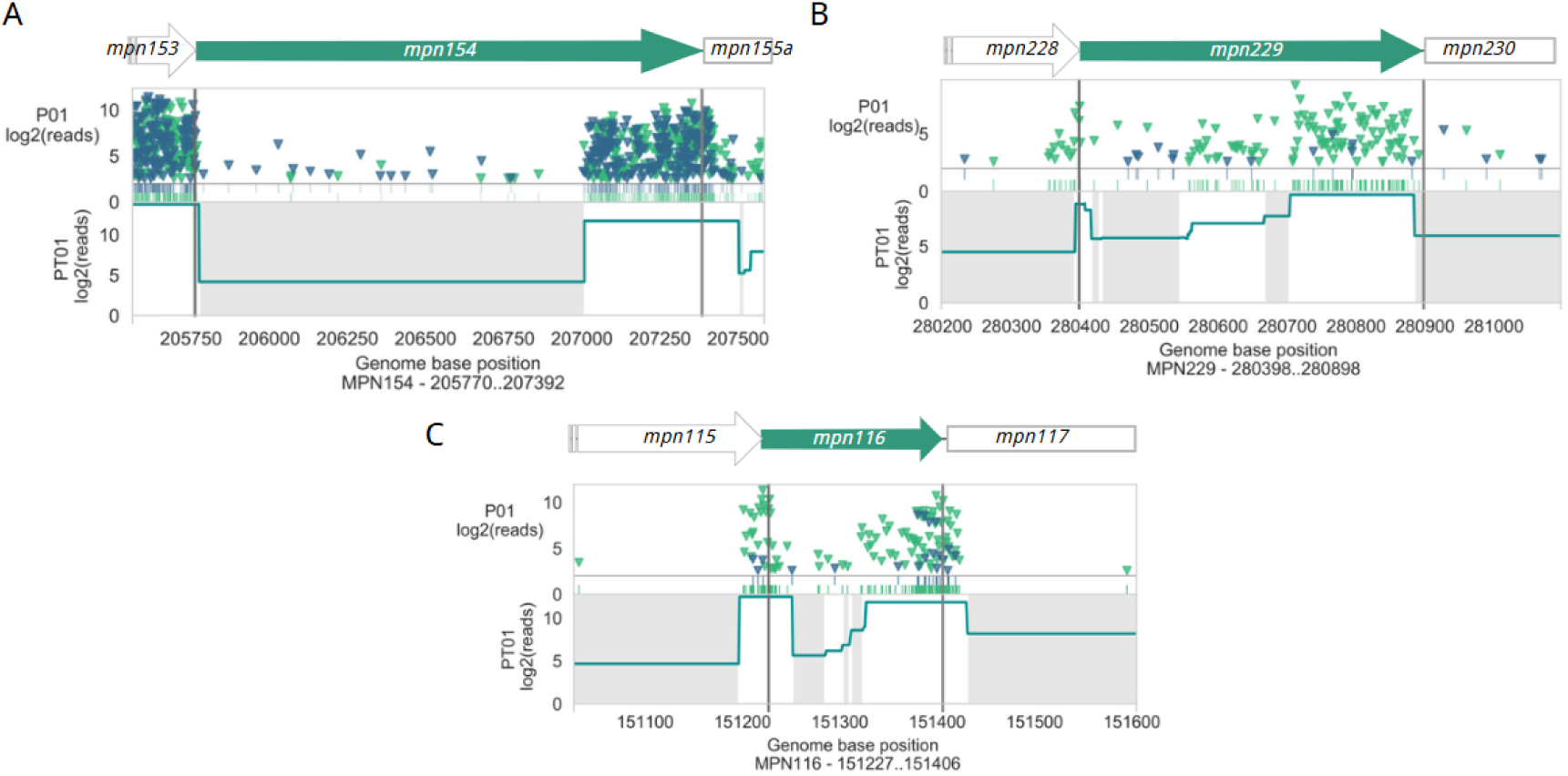
Examples of extended termini regions in E genes. Panels correspond to genes *mpn154* (A), *mpn229* (B) and *mpn116* (C). For each panel, the insertion profile of the region of interest at passage 1 is shown. Transposons containing promoters or terminators are depicted in blue and green triangles, respectively. Vertical lines show start and stop codons of the gene, also including their genomic context on top.

**Figure S17.**
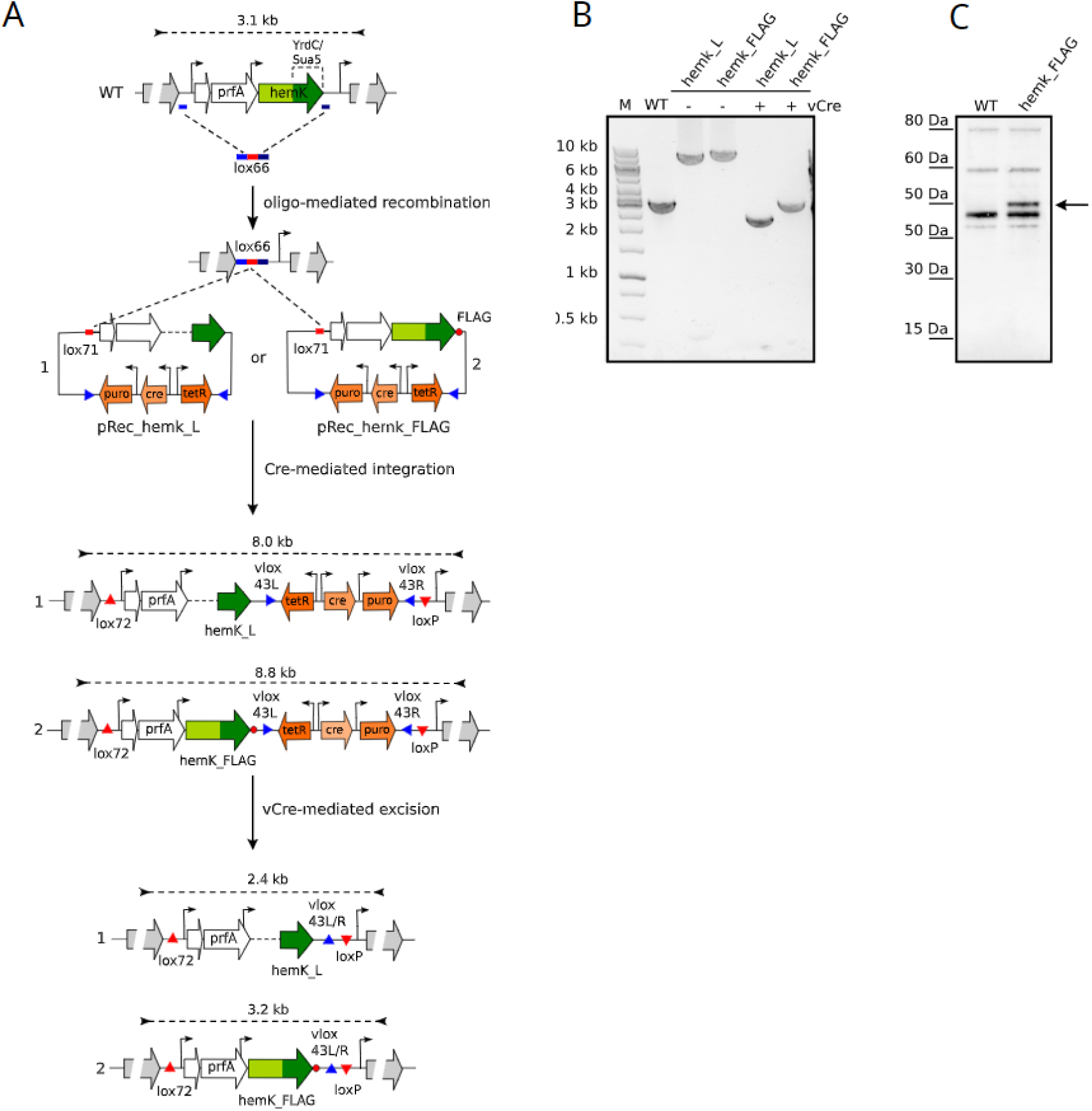
Construction of M. pneumoniae strains expressing *mpn362* gene variants to assess the requirement of the two predicted functional domains for cell viability. **A)** Schematic diagram showing the constructs and SURE-editing procedure used to generate *M. pneumoniae* strains expressing two *mpn362* gene (hemK) variants. Plasmids pRec_hemk_L and pRec_hemk_FLAG were constructed to replace the endogenous *hemK* (*mpn362*) locus by variants lacking the N-terminal Hemk domain (hemk_L), or containing a C-terminal flag (hemk_FLAG), respectively. Note that both plasmids also contain the *mpn360* (*rpmE*) and *mpn361* (*prfA)* genes to maintain the same genetic architecture after complementation. Briefly, deletion of the *rpmE*-*prfA-hemK* (*mpn360*-*mpn361*-*mpn362*) endogenous locus is mediated by oligo-recombineering using an oligo containing a lox66 recombination site (in red) flanked by homologous regions (in blue). Gene complementation with the mutant variants is then mediated by Cre-mediated integration of the plasmids mentioned above in the lox66 site, generating lox72 and loxP sites. Plasmid backbone (containing tetR repressor, Cre and puromycin resistance marker) is then removed from the genome by vCre-mediated recombination using the vlox sites present in the plasmid sequence. Sizes of the PCR products expected for each intermediate strain are shown above. **B)** PCR analyses using genomic DNA of the WT and intermediate strains before (labeled as “-”) and after (labeled as “+”) vCre excision are shown. The size of the expected PCR products is shown in panel A **C)** Western blot analysis of cell lysates of the WT and hemk_FLAG mutant using anti-FLAG antibodies. The arrow shows the protein band expected for the expression of the full-length *mpn362* coding-sequence, indicating that it is expressed as a fusion protein.

**Figure S18.**
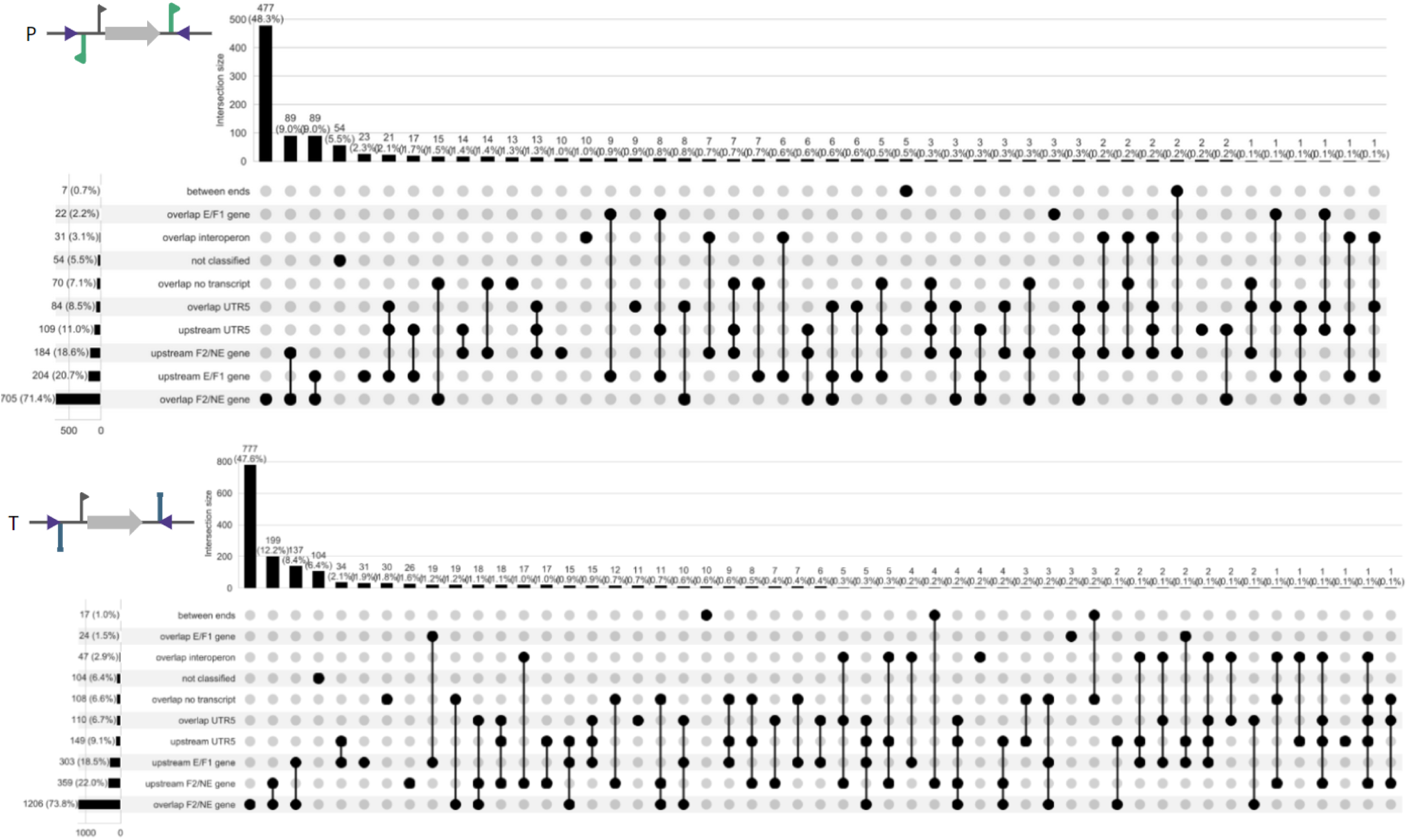
Upset plots representing the different contexts where unique P or T insertions are maintained at nucleobase level. P- and T-selected positions are represented in the top and bottom panels, respectively (associated to construct schemes). Left bar plots account for the different contexts where these insertions are found, represented as total number and percentage (between parentheses). Then, an intersection graph is presented to capture all the possible scenarios with a black solid circle linked with lines when present in several contexts. Top bar plots on each figure account for the number of events in each set.

**Supplementary Table 1.**
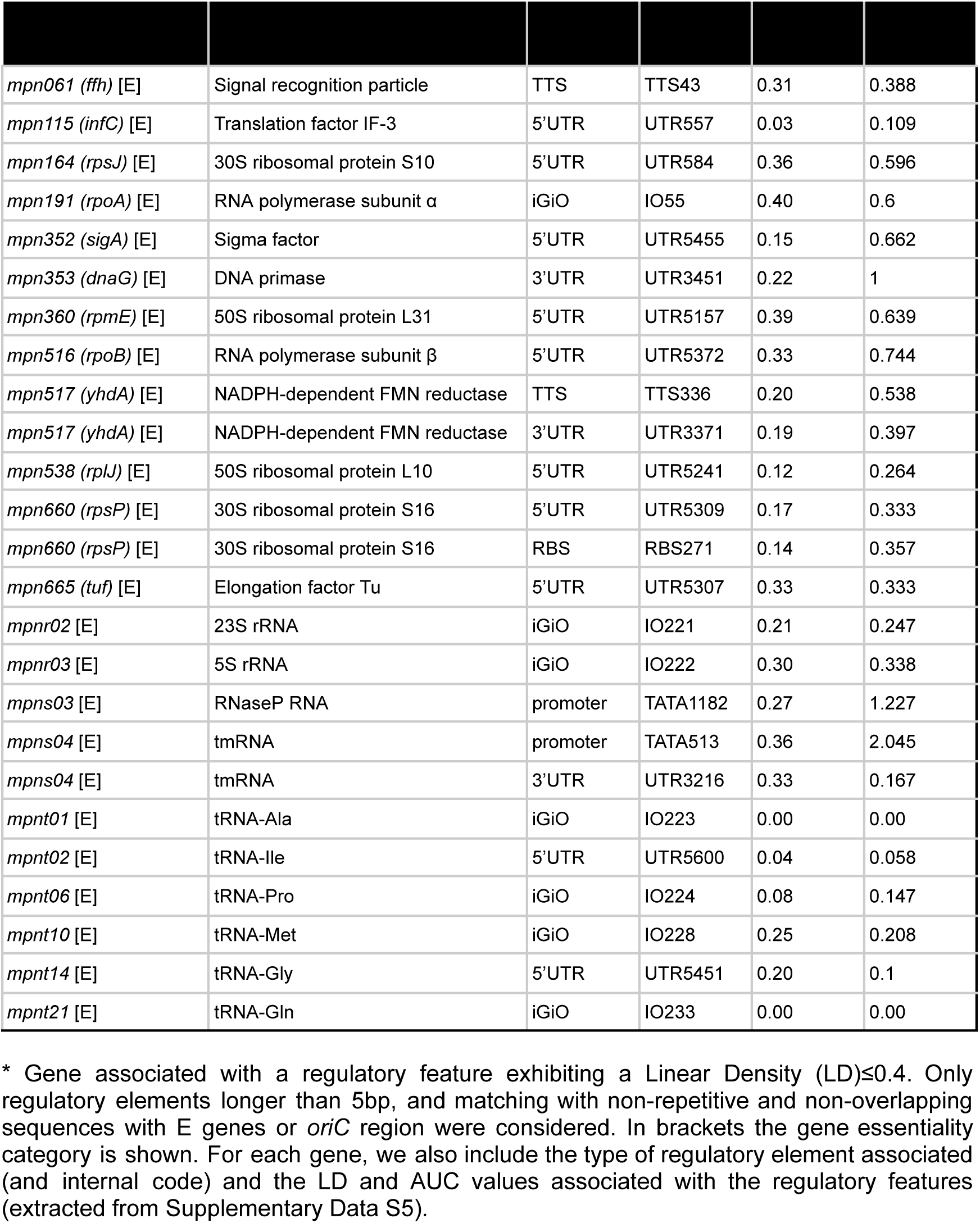
Essential regulatory features.

| *mpn061 (ffh)* [E] | Signal recognition particle | TTS | TTS43 | 0.31 | 0.388 |
| --- | --- | --- | --- | --- | --- |
| *mpn115 (infC)* [E] | Translation factor IF-3 | 5'UTR | UTR557 | 0.03 | 0.109 |
| *mpn164 (rpsJ)* [E] | 30S ribosomal protein S10 | 5'UTR | UTR584 | 0.36 | 0.596 |
| *mpn191 (rpoA)* [E] | RNA polymerase subunit $\alpha$ | iGiO | IO55 | 0.40 | 0.6 |
| *mpn352 (sigA)* [E] | Sigma factor | 5'UTR | UTR5455 | 0.15 | 0.662 |
| *mpn353 (dnaG)* [E] | DNA primase | 3'UTR | UTR3451 | 0.22 | 1 |
| *mpn360 (rpmE)* [E] | 50S ribosomal protein L31 | 5'UTR | UTR5157 | 0.39 | 0.639 |
| *mpn516 (rpoB)* [E] | RNA polymerase subunit $\beta$ | 5'UTR | UTR5372 | 0.33 | 0.744 |
| *mpn517 (yhdA)* [E] | NADPH-dependent FMN reductase | TTS | TTS336 | 0.20 | 0.538 |
| *mpn517 (yhdA)* [E] | NADPH-dependent FMN reductase | 3'UTR | UTR3371 | 0.19 | 0.397 |
| *mpn538 (rplJ)* [E] | 50S ribosomal protein L10 | 5'UTR | UTR5241 | 0.12 | 0.264 |
| *mpn660 (rpsP)* [E] | 30S ribosomal protein S16 | 5'UTR | UTR5309 | 0.17 | 0.333 |
| *mpn660 (rpsP)* [E] | 30S ribosomal protein S16 | RBS | RBS271 | 0.14 | 0.357 |
| *mpn665 (tuf)* [E] | Elongation factor Tu | 5'UTR | UTR5307 | 0.33 | 0.333 |
| *mpnr02* [E] | 23S rRNA | iGiO | IO221 | 0.21 | 0.247 |
| *mpnr03* [E] | 5S rRNA | iGiO | IO222 | 0.30 | 0.338 |
| *mpns03* [E] | RNaseP RNA | promoter | TATA1182 | 0.27 | 1.227 |
| *mpns04* [E] | tmRNA | promoter | TATA513 | 0.36 | 2.045 |
| *mpns04* [E] | tmRNA | 3'UTR | UTR3216 | 0.33 | 0.167 |
| *mpnt01* [E] | tRNA-Ala | iGiO | IO223 | 0.00 | 0.00 |
| *mpnt02* [E] | tRNA-Ile | 5'UTR | UTR5600 | 0.04 | 0.058 |
| *mpnt06* [E] | tRNA-Pro | iGiO | IO224 | 0.08 | 0.147 |
| *mpnt10* [E] | tRNA-Met | iGiO | IO228 | 0.25 | 0.208 |
| *mpnt14* [E] | tRNA-Gly | 5'UTR | UTR5451 | 0.20 | 0.1 |
| *mpnt21* [E] | tRNA-Gln | iGiO | IO233 | 0.00 | 0.00 |

\* Gene associated with a regulatory feature exhibiting a Linear Density (LD) ≤ 0.4. Only regulatory elements longer than 5bp, and matching with non-repetitive and non-overlapping sequences with E genes or *oriC* region were considered. In brackets the gene essentiality category is shown. For each gene, we also include the type of regulatory element associated (and internal code) and the LD and AUC values associated with the regulatory features (extracted from Supplementary Data S5).

**Supplementary Table 2.** Genes encoding protein domains or protein extensions with different essentiality profiles.

| *mpn030* | NusB-like fold protein | F1 | 0,775 | 34468 | 34560 | extension | 31 | 1,382 | 34561 | 34971 | PF09185 | 137 | 0,373 |
| --- | --- | --- | --- | --- | --- | --- | --- | --- | --- | --- | --- | --- | --- |
| *mpn031* | Hypothetical protein | F1 | 0,831 | 34974 | 35456 | PF17534 | 161 | 0,35 | 35457 | 35582 | extension | 42 | 1,687 |
| *mpn133* | Cytotoxic nuclease | F2 | 2,653 | 172246 | 172938 | COG1525 | 231 | 1,001 | 172939 | 173148 | extension | 70 | 4,65 |
| *mpn154* | NusA | F1 | 0,669 | 205770 | 207017 | PF08529<br>PF13184 | 416 | 0,26 | 207018 | 207389 | extension | 124 | 1,46 |
| *mpn173* | 50S ribosomal protein L29 | F1 | 0,607 | 223005 | 223184 | PF00831 | 60 | 0,2 | 223185 | 223337 | extension | 51 | 0,902 |
| *mpn229* | Single-stranded DNA-binding protein | F1 | 0,963 | 280398 | 280709 | PF00436 | 104 | 0,407 | 280710 | 280895 | extension | 62 | 1,04 |
| *mpn231* | 50S ribosomal protein L9 | F1 | 0,74 | 281205 | 281477 | PF01281 | 91 | 0,425 | 281478 | 281651 | PF03948 | 58 | 0,845 |
| *mpn300* | DHFR / ScpA | F1 | 1,175 | 353498 | 354037 | PF00186 | 180 | 1,424 | 354038 | 355015 | PF02616 | 326 | 0,529 |
| *mpn351* | tRNA methyltransferase (TmrK) | F1 | 0,709 | 418720 | 418280 | PF04816 | 147 | 0,31 | 418279 | 418082 | extension | 66 | 1,215 |
| *mpn362* | methyltransferase (HemK) | F1 | 0,8 | 431670 | 432458 | PF05175 | 263 | 0,833 | 432459 | 433028 | PF01300 | 190 | 0,559 |
| *mpn431* | CbiQ family ECF transporter | F1 | 0,762 | 519797 | 518991 | PF02361 | 269 | 0,498 | 518990 | 518847 | extension | 48 | 1,74 |
| *mpn563* | GTP-binding protein Obg | F1 | 0,57 | 684489 | 683521 | PF01018 | 323 | 0,362 | 683520 | 683191 | PF09269 | 110 | 0,971 |
| *mpn597* | F0F1 ATP synthase subunit | F1 | 2,656 | 719508 | 719260 | PF02823 | 83 | 0,518 | 719259 | 719110 | extension | 50 | 3,973 |
| *mpn618* | DNA polymerase III subunit | F1 | 0,429 | 742307 | 741243 | PF13177 | 355 | 0,301 | 741242 | 740265 | extension | 326 | 0,392 |
| *mpn683* | ABC transporter | F1 | 1,299 | 806853 | 806563 | extension | 97 | 2,694 | 806562 | 805837 | PF00005 | 242 | 0,32 |
| *mpn688\** | ParA family protein | F2 | 3,165 | 816301 | 816155 | PF01656 | 49 | 1,17 | 816154 | 815492 | PF01656 | 221 | 2,367 |

<sup>a</sup> Protein domains are defined based on differential essentiality profiles.
<sup>b</sup> Conserved protein domains are assigned to COG or Pfam families. When the protein domain is not conserved and only present in mycoplasma proteins, the domain is annotated as protein “extension”. Only protein-coding genes exhibiting “extensions” covering more than 15% of the total protein length are presented.
<sup>c</sup> Amino acid length of the protein domain.
<sup>d</sup> Domain essentiality estimation based on linear density at PT1. AUC fitness score is shown in parenthesis.
\* The 49 amino acid N-terminal region of MPN688 may overlap with putative sequences corresponding to the *oriC* region.

**Supplementary Table 3.** Putative essential protein-coding genes that can be functionally split.

| *mpn034 (rpoC)* | DNA polymerase III subunit $\alpha$ | E | 0,39 | 40548 | 40591 | M193 | 1,364 |
| --- | --- | --- | --- | --- | --- | --- | --- |
| *mpn081 (glnQ)* | Gln transport ATP-binding protein | F1 | 0,757 | 100751 | 100759 | L65 | 3,111 |
|  |  |  |  | 100819 | 100940 | L114 | 2,004 |
| *mpn106 (pheT)* | Phenylalanyl-tRNA synthetase subunit $\beta$ | E | 0,407 | 137322 | 137353 | M208 | 2,609 |
| *mpn123 (parC)* | DNA topoisomerase 4 subunit A | E | 0,442 | 159490 | 159511 | M387 | 1,886 |
| *mpn160 (HP)* | Hypothetical protein | F1 | 0,617 | 213490 | 213514 | V96 | 2,94 |
| *mpn338 (HP)* | Hypothetical protein | E | 0,36 | 402288 | 402299 | V332 | 1,75 |
| *mpn357 (ligA)* | DNA ligase | E | 0,378 | 425716 | 425749 | M574 | 1,824 |
| *mpn378 (dnaE)* | DNA polymerase III subunit $\alpha$ | E | 0,349 | 454329 | 454342 | M249 | 0,857 |
| *mpn556 (argS)* | Arginyl-tRNA synthetase | E | 0,413 | 676187 | 676252 | M356 | 0,886 |

<sup>a</sup> Coordinates correspond to the genome sequence of the M129 lab strain (M129\_LSR) used in this study.
<sup>b</sup> Predicted first residue of the C-terminal fragment. When several alternative start sites were found in the disruptable region, we indicate the most likely position based on the persistence of transposon insertions.
<sup>c</sup> AUC value calculated for the detected disruptable region within the essential sequence of the protein coding gene.

**Supplementary Table 4.**
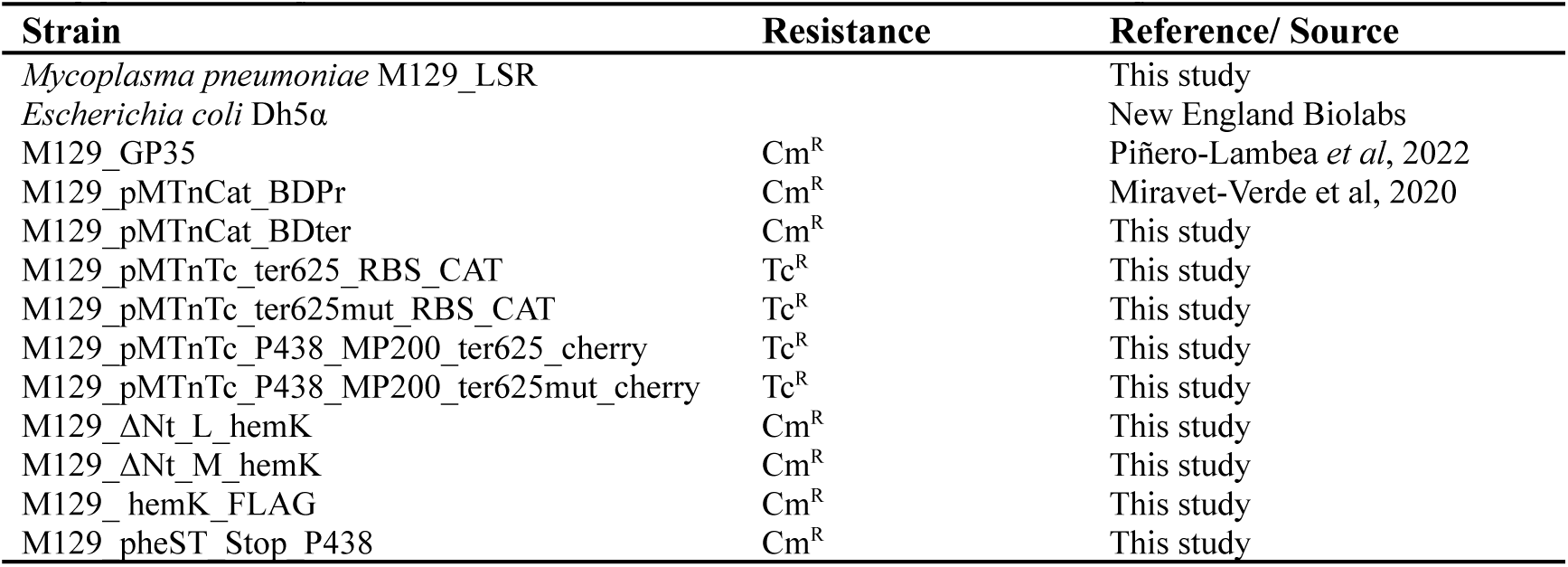
Bacterial strains used in this study.

**Supplementary Table 5.**
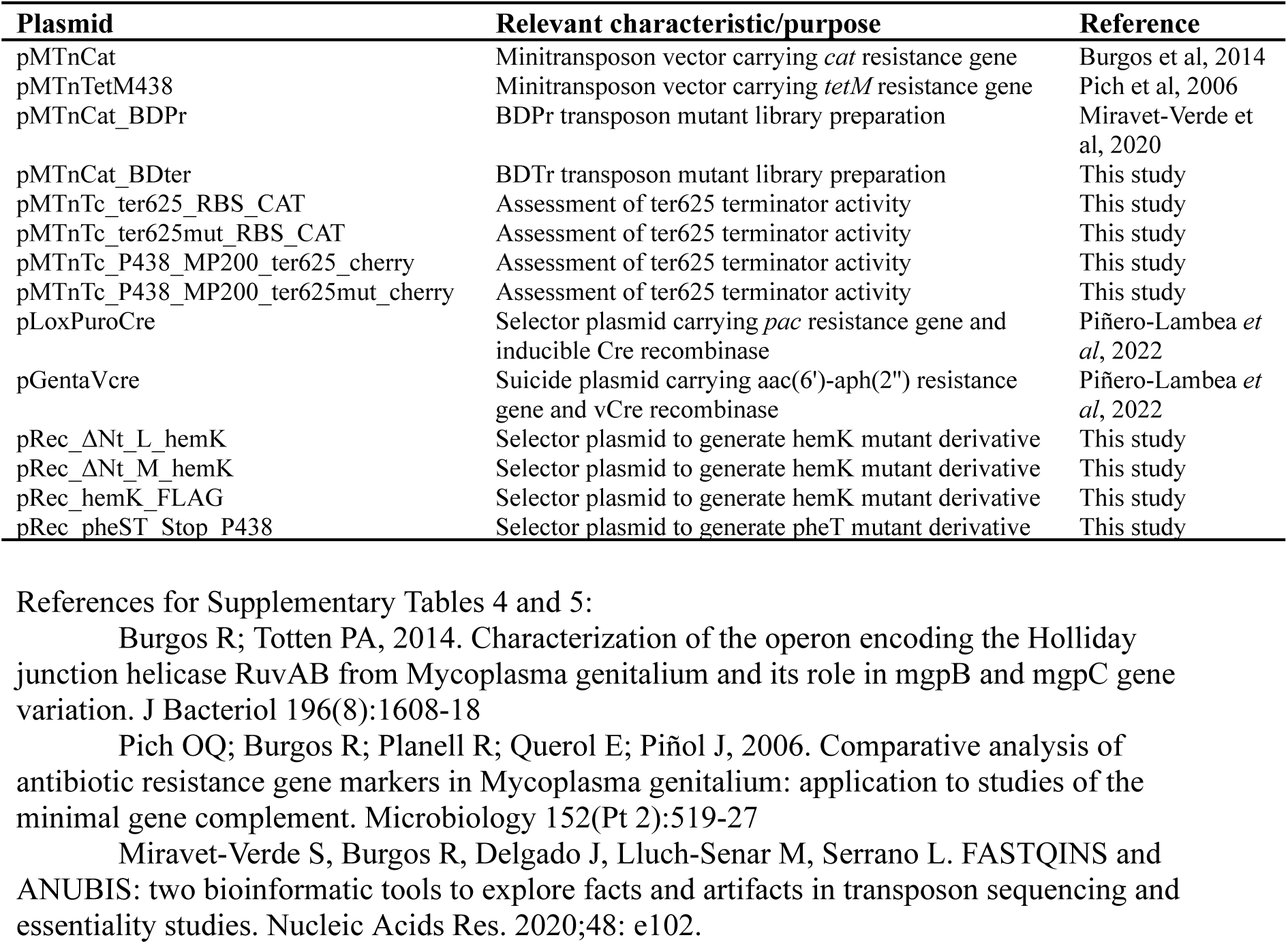
Plasmids used in this study.

| Plasmid | Relevant characteristic/purpose | Reference |
| --- | --- | --- |
| pMTnCat | Minitransposon vector carrying *cat* resistance gene | Burgos *et al*, 2014 |
| pMTnTetM438 | Minitransposon vector carrying *tetM* resistance gene | Pich *et al*, 2006 |
| pMTnCat_BDPr | BDPr transposon mutant library preparation | Miravet-Verde *et al*, 2020 |
| pMTnCat_BDter | BDTr transposon mutant library preparation | This study |
| pMTnTc_ter625_RBS_CAT | Assessment of ter625 terminator activity | This study |
| pMTnTc_ter625mut_RBS_CAT | Assessment of ter625 terminator activity | This study |
| pMTnTc_P438_MP200_ter625_cherry | Assessment of ter625 terminator activity | This study |
| pMTnTc_P438_MP200_ter625mut_cherry | Assessment of ter625 terminator activity | This study |
| pLoxPuroCre | Selector plasmid carrying *pac* resistance gene and inducible Cre recombinase | Piñero-Lambea *et al*, 2022 |
| pGentaVcre | Suicide plasmid carrying aac(6')-aph(2'') resistance gene and vCre recombinase | Piñero-Lambea *et al*, 2022 |
| pRec_ΔNt_L_hemK | Selector plasmid to generate hemK mutant derivative | This study |
| pRec_ΔNt_M_hemK | Selector plasmid to generate hemK mutant derivative | This study |
| pRec_hemK_FLAG | Selector plasmid to generate hemK mutant derivative | This study |
| pRec_pheST_Stop_P438 | Selector plasmid to generate pheT mutant derivative | This study |

References for Supplementary Tables 4 and 5:
Burgos R; Totten PA, 2014. Characterization of the operon encoding the Holliday junction helicase RuvAB from *Mycoplasma genitalium* and its role in *mgpB* and *mgpC* gene variation. *J Bacteriol* 196(8):1608-18
Pich OQ; Burgos R; Planell R; Querol E; Piñol J, 2006. Comparative analysis of antibiotic resistance gene markers in *Mycoplasma genitalium*: application to studies of the minimal gene complement. *Microbiology* 152(Pt 2):519-27
Miravet-Verde S, Burgos R, Delgado J, Lluch-Senar M, Serrano L. FASTQINS and ANUBIS: two bioinformatic tools to explore facts and artifacts in transposon sequencing and essentiality studies. *Nucleic Acids Res.* 2020;48: e102.

**Supplementary Table 6.**
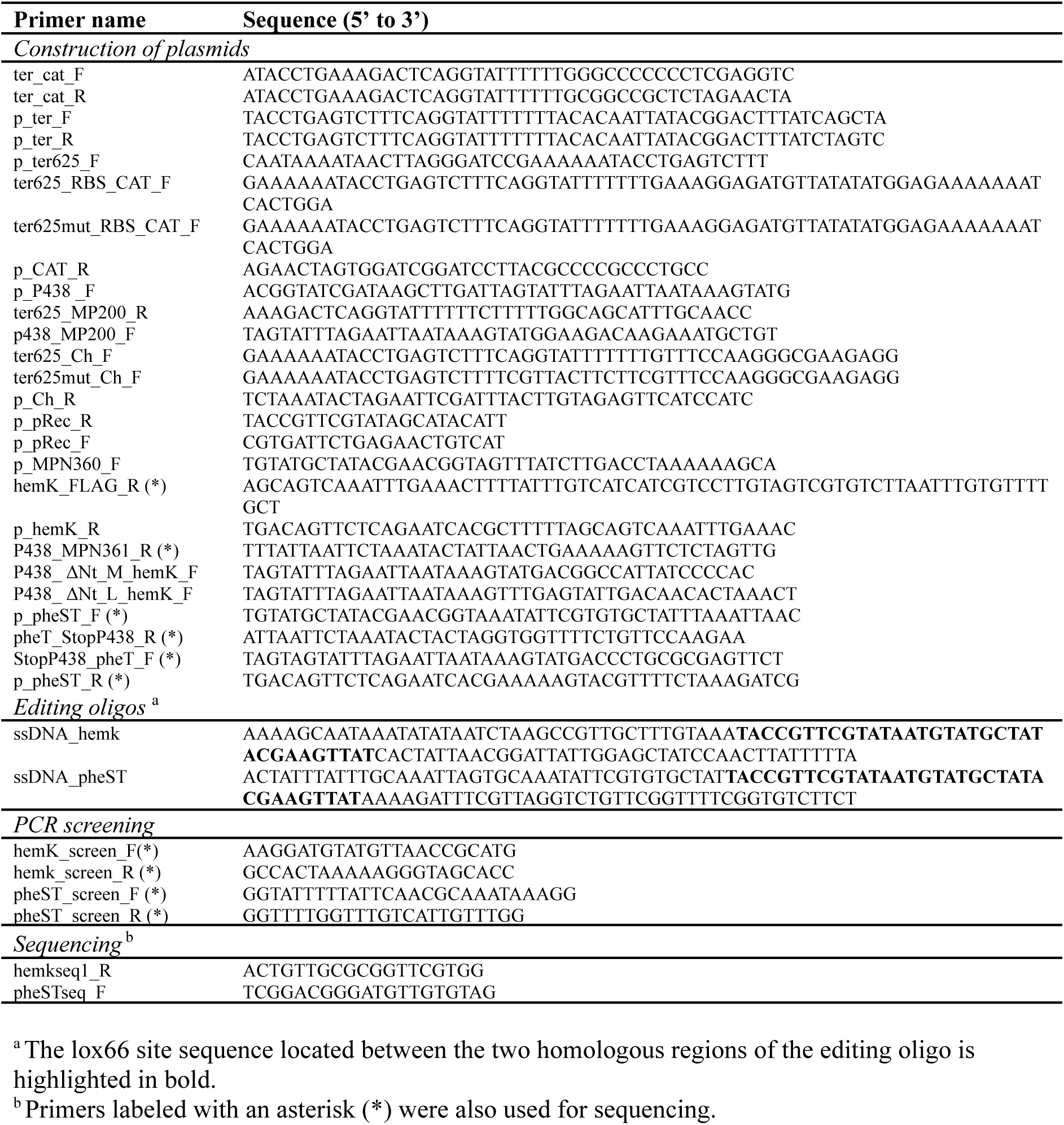
Primers used in this study.

**Additional Supplementary Data description**

**Supplementary Data S1 - Raw insertion profile data.** For each nucleotide position of the *M. pneumoniae* genome (column *A,* index in 0-base) we include the number of insertion events detected for each sample analyzed in this work (column name). Samples of the P library are named as PXX_RY, where X refers to the passage (1, 2, 3, 4, 6, 8, or 10) and Y to the replicate batch (1, 2, 3, or 4). For samples of the T library we use the nomenclature TXX_RY, while for the combination of both plasmid libraries we use PTXX_RY. When no replicate is associated, the sample represents the addition of the different replicates for that condition.

## REFERENCES

1. Yus E, Lloréns-Rico V, Martínez S, Gallo C, Eilers H, Blötz C, et al. Determination of the Gene Regulatory Network of a Genome-Reduced Bacterium Highlights Alternative Regulation Independent of Transcription Factors. Cell Syst. 2019;9: 143–158.e13.

2. Trussart M, Yus E, Martinez S, Baù D, Tahara YO, Pengo T, et al. Defined chromosome structure in the genome-reduced bacterium Mycoplasma pneumoniae. Nat Commun. 2017;8: 14665.

3. Hamer L, DeZwaan TM, Montenegro-Chamorro MV, Frank SA, Hamer JE. Recent advances in large-scale transposon mutagenesis. Curr Opin Chem Biol. 2001;5: 67–73.

4. Basta DW, Bergkessel M, Newman DK. Identification of Fitness Determinants during Energy-Limited Growth Arrest in Pseudomonas aeruginosa. mBio. 2017. doi:10.1128/mbio.01170-17

5. Price MN, Zane GM, Kuehl JV, Melnyk RA, Wall JD, Deutschbauer AM, et al. Filling gaps in bacterial amino acid biosynthesis pathways with high-throughput genetics. PLoS Genet. 2018;14: e1007147.

6. Salama NR, Shepherd B, Falkow S. Global transposon mutagenesis and essential gene analysis of Helicobacter pylori. J Bacteriol. 2004;186: 7926–7935.

7. Christen B, Abeliuk E, Collier JM, Kalogeraki VS, Passarelli B, Coller JA, et al. The essential genome of a bacterium. Mol Syst Biol. 2011;7: 528.

8. Langridge GC, Phan M-D, Turner DJ, Perkins TT, Parts L, Haase J, et al. Simultaneous assay of every Salmonella Typhi gene using one million transposon mutants. Genome Research. 2009. pp. 2308–2316. doi:10.1101/gr.097097.109

9. Barquist L, Boinett CJ, Cain AK. Approaches to querying bacterial genomes with transposon-insertion sequencing. RNA Biol. 2013;10: 1161–1169.

10. Mann B, van Opijnen T, Wang J, Obert C, Wang Y-D, Carter R, et al. Control of virulence by small RNAs in Streptococcus pneumoniae. PLoS Pathog. 2012;8: e1002788.

11. Capel E, Zomer AL, Nussbaumer T, Bole C, Izac B, Frapy E, et al. Comprehensive Identification of Meningococcal Genes and Small Noncoding RNAs Required for Host Cell Colonization. mBio. 2016. doi:10.1128/mbio.01173-16

12. Barquist L, Langridge GC, Turner DJ, Phan M-D, Turner AK, Bateman A, et al. A comparison of dense transposon insertion libraries in the Salmonella serovars Typhi and Typhimurium. Nucleic Acids Res. 2013;41: 4549–4564.

13. Zhang YJ, Ioerger TR, Huttenhower C, Long JE, Sassetti CM, Sacchettini JC, et al. Global assessment of genomic regions required for growth in Mycobacterium tuberculosis. PLoS Pathog. 2012;8: e1002946.

14. Coe KA, Lee W, Stone MC, Komazin-Meredith G, Meredith TC, Grad YH, et al. Multi-strain Tn-Seq reveals common daptomycin resistance determinants in Staphylococcus aureus. PLoS Pathog. 2019;15: e1007862.

15. Güell M, van Noort V, Yus E, Chen W-H, Leigh-Bell J, Michalodimitrakis K, et al. Transcriptome Complexity in a Genome-Reduced Bacterium. Science. 2009. pp. 1268–1271. doi:10.1126/science.1176951

16. Yus E, Maier T, Michalodimitrakis K, van Noort V, Yamada T, Chen W-H, et al. Impact of Genome Reduction on Bacterial Metabolism and Its Regulation. Science. 2009. pp. 1263–1268. doi:10.1126/science.1177263

17. Kühner S, van Noort V, Betts MJ, Leo-Macias A, Batisse C, Rode M, et al. Proteome Organization in a Genome-Reduced Bacterium. Science. 2009. pp. 1235–1240. doi:10.1126/science.1176343

18. Maier T, Schmidt A, Güell M, Kühner S, Gavin A, Aebersold R, et al. Quantification of mRNA and protein and integration with protein turnover in a bacterium. Molecular Systems Biology. 2011. p. 511. doi:10.1038/msb.2011.38

19. Maier T, Marcos J, Wodke JAH, Paetzold B, Liebeke M, Gutiérrez-Gallego R, et al. Large-scale metabolome analysis and quantitative integration with genomics and proteomics data in Mycoplasma pneumoniae. Mol Biosyst. 2013;9: 1743–1755.

20. Chen W-H, van Noort V, Lluch-Senar M, Hennrich ML, Wodke JAH, Yus E, et al. Integration of multi-omics data of a genome-reduced bacterium: Prevalence of post-transcriptional regulation and its correlation with protein abundances. Nucleic Acids Res. 2016;44: 1192–1202.

21. Lluch-Senar M, Luong K, Lloréns-Rico V, Delgado J, Fang G, Spittle K, et al. Comprehensive methylome characterization of Mycoplasma genitalium and Mycoplasma pneumoniae at single-base resolution. PLoS Genet. 2013;9: e1003191.

22. Burgos R, Weber M, Martinez S, Lluch-Senar M, Serrano L. Protein quality control and regulated proteolysis in the genome-reduced organism Mycoplasma pneumoniae. Molecular Systems Biology. 2020. doi:10.15252/msb.20209530

23. Lluch-Senar M, Delgado J, Chen W-H, Lloréns-Rico V, O’Reilly FJ, Wodke JA, et al. Defining a minimal cell: essentiality of small ORFs and ncRNAs in a genome-reduced bacterium. Mol Syst Biol. 2015;11: 780.

24. Burgos R, Totten PA. Characterization of the operon encoding the Holliday junction helicase RuvAB from Mycoplasma genitalium and its role in mgpB and mgpC gene variation. J Bacteriol. 2014;196: 1608–1618.

25. Lyon BR, May JW, Skurray RA. Tn4001: a gentamicin and kanamycin resistance transposon in Staphylococcus aureus. Mol Gen Genet. 1984;193: 554–556.

26. Pich OQ, Burgos R, Planell R, Querol E, Piñol J. Comparative analysis of antibiotic resistance gene markers in Mycoplasma genitalium: application to studies of the minimal gene complement. Microbiology. 2006;152: 519–527.

27. Nakagawa S, Niimura Y, Miura K-I, Gojobori T. Dynamic evolution of translation initiation mechanisms in prokaryotes. Proc Natl Acad Sci U S A. 2010;107: 6382–6387.

28. Montero-Blay A, Miravet-Verde S, Lluch-Senar M, Piñero-Lambea C, Serrano L. SynMyco transposon: engineering transposon vectors for efficient transformation of minimal genomes. DNA Res. 2019;26: 327–339.

29. Miravet-Verde S, Burgos R, Delgado J, Lluch-Senar M, Serrano L. FASTQINS and ANUBIS: two bioinformatic tools to explore facts and artifacts in transposon sequencing and essentiality studies. Nucleic Acids Res. 2020;48: e102.

30. Laslett D, Canback B. ARAGORN, a program to detect tRNA genes and tmRNA genes in nucleotide sequences. Nucleic Acids Res. 2004;32: 11–16.

31. Murphy FV 4th, Ramakrishnan V. Structure of a purine-purine wobble base pair in the decoding center of the ribosome. Nat Struct Mol Biol. 2004;11: 1251–1252.

32. Lloréns-Rico V, Cano J, Kamminga T, Gil R, Latorre A, Chen W-H, et al. Bacterial antisense RNAs are mainly the product of transcriptional noise. Sci Adv. 2016;2: e1501363.

33. Hutchison CA 3rd, Chuang R-Y, Noskov VN, Assad-Garcia N, Deerinck TJ, Ellisman MH, et al. Design and synthesis of a minimal bacterial genome. Science. 2016;351: aad6253.

34. Yus E, Güell M, Vivancos AP, Chen W-H, Lluch-Senar M, Delgado J, et al. Transcription start site associated RNAs in bacteria. Mol Syst Biol. 2012;8: 585.

35. Tietze L, Lale R. Importance of the 5’ regulatory region to bacterial synthetic biology applications. Microb Biotechnol. 2021;14: 2291–2315.

36. Shepherd J, Ibba M. Bacterial transfer RNAs. FEMS Microbiol Rev. 2015;39: 280–300.

37. Burgos R, Weber M, Gallo C, Lluch-Senar M, Serrano L. Widespread ribosome stalling in a genome-reduced bacterium and the need for translational quality control. iScience. 2021;24: 102985.

38. Ren G-X, Guo X-P, Sun Y-C. Regulatory 3′ Untranslated Regions of Bacterial mRNAs. Frontiers in Microbiology. 2017. doi:10.3389/fmicb.2017.01276

39. Torres-Puig S, Broto A, Querol E, Piñol J, Pich OQ. A novel sigma factor reveals a unique regulon controlling cell-specific recombination in Mycoplasma genitalium. Nucleic Acids Res. 2015;43: 4923–4936.

40. Reynolds R, Chamberlin MJ. Parameters affecting transcription termination by Escherichia coli RNA. II. Construction and analysis of hybrid terminators. J Mol Biol. 1992;224: 53–63.

41. Blötz C, Lartigue C, Valverde Timana Y, Ruiz E, Paetzold B, Busse J, et al. Development of a replicating plasmid based on the native oriC in Mycoplasma pneumoniae. Microbiology. 2018;164: 1372–1382.

42. Wolański M, Donczew R, Zawilak-Pawlik A, Zakrzewska-Czerwińska J. oriC-encoded instructions for the initiation of bacterial chromosome replication. Front Microbiol. 2015;0. doi:10.3389/fmicb.2014.00735

43. Soppa J, Kobayashi K, Noirot-Gros M-F, Oesterhelt D, Ehrlich SD, Dervyn E, et al. Discovery of two novel families of proteins that are proposed to interact with prokaryotic SMC proteins, and characterization of the Bacillus subtilis family members ScpA and ScpB. Mol Microbiol. 2002;45: 59–71.

44. El Yacoubi B, Lyons B, Cruz Y, Reddy R, Nordin B, Agnelli F, et al. The universal YrdC/Sua5 family is required for the formation of threonylcarbamoyladenosine in tRNA. Nucleic Acids Res. 2009;37: 2894–2909.

45. Heurgue-Hamard V. The hemK gene in Escherichia coli encodes the N5-glutamine methyltransferase that modifies peptide release factors. The EMBO Journal. 2002. pp. 769–778. doi:10.1093/emboj/21.4.769

46. Dinçbas-Renqvist V, Engström A, Mora L, Heurgué-Hamard V, Buckingham R, Ehrenberg M. A post-translational modification in the GGQ motif of RF2 from Escherichia coli stimulates termination of translation. EMBO J. 2000;19: 6900–6907.

47. Piñero-Lambea C, Garcia-Ramallo E, Miravet-Verde S, Burgos R, Scarpa M, Serrano L, et al. SURE editing: combining oligo-recombineering and programmable insertion/deletion of selection markers to efficiently edit the Mycoplasma pneumoniae genome. Nucleic Acids Res. 2022;50: e127.

48. Pich OQ, Burgos R, Ferrer-Navarro M, Querol E, Piñol J. Mycoplasma genitalium mg200 and mg386 genes are involved in gliding motility but not in cytadherence. Mol Microbiol. 2006;60: 1509–1519.

49. Cloward JM, Krause DC. Mycoplasma pneumoniae J-domain protein required for terminal organelle function. Mol Microbiol. 2009;71: 1296–1307.

50. Miravet-Verde S, Ferrar T, Espadas-García G, Mazzolini R, Gharrab A, Sabido E, et al. Unraveling the hidden universe of small proteins in bacterial genomes. Mol Syst Biol. 2019;15: e8290.

51. Miravet-Verde S, Mazzolini R, Segura-Morales C, Broto A, Lluch-Senar M, Serrano L. ProTInSeq: transposon insertion tracking by ultra-deep DNA sequencing to identify translated large and small ORFs. Nat Commun. 2024;15: 2091.

52. Mosyak L, Reshetnikova L, Goldgur Y, Delarue M, Safro MG. Structure of phenylalanyl-tRNA synthetase from Thermus thermophilus. Nature Structural Biology. 1995. pp. 537–547. doi:10.1038/nsb0795-537

53. Roy H, Ibba M. Phenylalanyl-tRNA synthetase contains a dispensable RNA-binding domain that contributes to the editing of noncognate aminoacyl-tRNA. Biochemistry. 2006;45: 9156–9162.

54. Goldgur Y, Mosyak L, Reshetnikova L, Ankilova V, Lavrik O, Khodyreva S, et al. The crystal structure of phenylalanyl-tRNA synthetase from thermus thermophilus complexed with cognate tRNAPhe. Structure. 1997;5: 59–68.

55. Larrimore KE, Rancati G. The conditional nature of gene essentiality. Curr Opin Genet Dev. 2019;58–59: 55–61.

56. Danchin A, Fang G. Unknown unknowns: essential genes in quest for function. Microb Biotechnol. 2016;9: 530–540.

57. Luhua S, Hegie A, Suzuki N, Shulaev E, Luo X, Cenariu D, et al. Linking genes of unknown function with abiotic stress responses by high-throughput phenotype screening. Physiol Plant. 2013;148: 322–333.

58. Cain AK, Barquist L, Goodman AL, Paulsen IT, Parkhill J, van Opijnen T. A decade of advances in transposon-insertion sequencing. Nat Rev Genet. 2020;21: 526–540.

59. Zhu J, Gong R, Zhu Q, He Q, Xu N, Xu Y, et al. Genome-Wide Determination of Gene Essentiality by Transposon Insertion Sequencing in Yeast Pichia pastoris. Sci Rep. 2018;8: 1–13.

60. Ghosh A, Komar AA. Eukaryote-specific extensions in ribosomal proteins of the small subunit: Structure and function. Translation (Austin). 2015;3: e999576.

61. Zhou X, Liao W-J, Liao J-M, Liao P, Lu H. Ribosomal proteins: functions beyond the ribosome. J Mol Cell Biol. 2015;7: 92–104.

62. Aseev LV, Boni IV. [Extraribosomal functions of bacterial ribosomal proteins]. Mol Biol. 2011;45: 805–816.

63. O’Reilly FJ, Xue L, Graziadei A, Sinn L, Lenz S, Tegunov D, et al. In-cell architecture of an actively transcribing-translating expressome. Science. 2020;369: 554–557.

64. Smith N, Wilson MA. Structural Biology of the DJ-1 Superfamily. Adv Exp Med Biol. 2017;1037: 5–24.

65. Rodríguez-Rojas A, Blázquez J. The Pseudomonas aeruginosa pfpI Gene Plays an Antimutator Role and Provides General Stress Protection. Journal of Bacteriology. 2009. pp. 844–850. doi:10.1128/jb.01081-08

66. Sturm M, Schroeder C, Bauer P. SeqPurge: highly-sensitive adapter trimming for paired-end NGS data. BMC Bioinformatics. 2016;17: 1–7.

67. Bankevich A, Nurk S, Antipov D, Gurevich AA, Dvorkin M, Kulikov AS, et al. SPAdes: a new genome assembly algorithm and its applications to single-cell sequencing. J Comput Biol. 2012;19: 455–477.

68. Gurevich A, Saveliev V, Vyahhi N, Tesler G. QUAST: quality assessment tool for genome assemblies. Bioinformatics. 2013;29: 1072–1075.

69. Khelik K, Lagesen K, Sandve GK, Rognes T, Nederbragt AJ. NucDiff: in-depth characterization and annotation of differences between two sets of DNA sequences. BMC Bioinformatics. 2017;18: 338.

70. Lloréns-Rico V, Lluch-Senar M, Serrano L. Distinguishing between productive and abortive promoters using a random forest classifier in Mycoplasma pneumoniae. Nucleic Acids Res. 2015;43: 3442–3453.

71. Naville M, Ghuillot-Gaudeffroy A, Marchais A, Gautheret D. ARNold: A web tool for the prediction of Rho-independent transcription terminators. RNA Biology. 2011. pp. 11–13. doi:10.4161/rna.8.1.13346

72. Weber M, Burgos R, Yus E, Yang J-S, Lluch-Senar M, Serrano L. Impact of C-terminal amino acid composition on protein expression in bacteria. Mol Syst Biol. 2020;16: e9208.

73. Garreta R, Moncecchi G. Learning scikit-learn: Machine Learning in Python. Packt Publishing Ltd; 2013.

74. Harris CR, Millman KJ, van der Walt SJ, Gommers R, Virtanen P, Cournapeau D, et al. Array programming with NumPy. Nature. 2020;585: 357–362.

